# Origin Specific Genomic Selection: a simple process to optimize the favourable contribution of parents to progeny

**DOI:** 10.1101/2019.12.13.875690

**Authors:** Chin Jian Yang, Rajiv Sharma, Gregor Gorjanc, Sarah Hearne, Wayne Powell, Ian Mackay

## Abstract

Modern crop breeding is in constant demand for new genetic diversity as part of the arms race with genetic gain. The elite gene pool has limited genetic variation and breeders are trying to introduce novelty from unadapted germplasm, landraces and wild relatives. For polygenic traits, currently available approaches to introgression are not ideal, as there is a demonstrable bias against exotic alleles during selection. Here, we propose a partitioned form of genomic selection, called Origin Specific Genomic Selection (OSGS), where we identify and target selection on favourable exotic alleles. Briefly, within a population derived from a bi-parental cross, we isolate alleles originating from the elite and exotic parents, which then allows us to separate out the predicted marker effects based on the allele origins. We validated the usefulness of OSGS using two nested association mapping (NAM) datasets: barley NAM (elite-exotic) and maize NAM (elite-elite), as well as by computer simulation. Our results suggest that OSGS works well in bi-parental crosses, and it is possible to extend the approach to broader multi-parental populations.

## Introduction

There is a general concern that the genetic base of elite varieties of many crops has become very narrow, diminishing the ability of the farming landscape to respond positively and quickly to new challenges. To continue to introduce novel, high value genetic variation into the elite gene pool, breeding programmes can select from crosses between their germplasm and materials from plant genetic resources; including wild species, landraces, and improved germplasm that are unadapted to the target environment. In these exotic crosses, marker assisted selection and backcrossing can effectively track a limited number of QTL accounting for a large proportion of the genetic variation for traits such as disease resistances. For highly polygenic traits, the introgression of novel variation from exotic sources is more complex for a number of reasons. Firstly, QTL mapping for polygenic traits is ineffective or may capture only a small proportion of the genetic variation. Secondly, the breeding scheme and population size needed to effectively pyramid many QTLs are unmanageable. Thirdly, loci at which the exotic lines carry a favourable allele are often linked in repulsion with loci at which the elite lines carry a favourable allele. Consequently, selection on segregating populations from elite-exotic crosses tends to select for the elite background and favourable contribution from the exotic may be lost through linkage drag, or equivalently through hitchhiking. Further, it can be difficult to phenotype adaptive traits, including yield, in populations with a high proportion of unadapted/exotic germplasm. For this reason, selection is often restricted to populations derived from the first or second backcross to the elite parent, in which the average exotic contribution is one quarter or one eighth. However, this practice increases further the risk of loss of favourable alleles introduced from the exotic parent.

To overcome these problems, additional generations of crossing among progeny prior to selection can be made to reduce repulsion linkage. Recurrent selection programmes have also been proposed to increase the frequency of favourable alleles from both elite and exotic donors over several generations (Hallauer and Carena 2012). Bernardo (2009) proposed genomic recurrent selection starting in the F_2_ before deriving recombinant lines and found this to be more effective than the conventional practice of selecting among lines derived from the backcross to the elite parent. More recently, Gorjanc et al. (2016) proposed genomic selection on a population established among exotic accessions to increase the frequency of favourable alleles prior to making crosses between the elite and (improved) exotic population. In simulation, this reduced the loss of favourable alleles from exotic sources compared to direct crossing. However, there is a risk that selective effort is wasted in increasing favourable allele frequencies in the improved exotic pool that are already at high frequency among elite lines. Ru and Bernardo (2019a and 2019b) proposed backcrossing favourable linkage groups instead of QTLs from exotic parent into elite parent using soybean nested association mapping (NAM) data as an example. Recently, Allier et al. (2019a) proposed treating parental genome contribution as a trait in its own right, and suggested index or truncation selection on this and agronomic traits as a means of reducing the loss of favourable exotic alleles. In addition, Allier et al. (2019b) proposed a method to identify exotic candidates that can provide the most benefit in elite-exotic crosses through maximizing favourable contributions from exotic parents.

The problems associated with introgression programmes for quantitative traits also manifest in mainstream breeding programmes. In a cross between two elite inbred lines, the favourable alleles at loci determining a polygenic trait are unlikely to be distributed equally between the two parents. For genetic progress, descendant lines must be selected in which both parents contribute favourable alleles, since only then can the performance of descendants exceed that of the best parent. Assuming for simplicity that all gene effects are equal, the selected line must be fixed for more favourable alleles than the best parent. However, selection among progeny may still result in a disproportionate contribution from the genome of the best parent. For example, Fradgley et al. (2019) found it common for an elite wheat line to share over 80% of its genetic material with one of its two elite parents.

In this paper we propose a simple process to quantify and therefore control the favourable contribution of parents to progeny with a technique called Origin Specific Genomic Selection (OSGS). We achieve this by partitioning a genomic prediction equation into two components: the first component is the contribution from markers where the favourable allele is carried by the primary (often elite) parent and the second component is the contribution from markers where the favourable allele is carried by the secondary (often exotic) parent. We test this method by within-cross prediction in two NAM datasets. The first is a barley NAM of backcross derived lines from an elite variety (Barke) and 25 wild barleys (Maurer et al. 2015). The second is the maize NAM of F_2_ derived lines from crosses between the inbred B73 and 25 lines selected to sample diversity among elite maize germplasm (Yu et al. 2008). We validate our results by computer simulations and discuss the implications of our results for introgression and pre-breeding together with broader applications in plant breeding, including the use of OSGS in multi-parental populations.

## Materials and Methods

### 1. Genomic Selection (GS) and Origin Specific Genomic Selection (OSGS)

The mixed linear model commonly used in the training population of genomic selection (GS) can be generalized as:

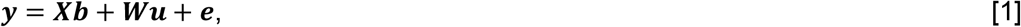

where ***y*** is a vector of observed trait values for each individual,

***X*** is a design matrix associating fixed effects with trait observations,
***b*** is a vector of fixed effects,
***W*** is a design matrix associating marker effects with trait observations,
***u*** is a vector of marker effects with an assumed distribution of 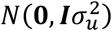,
***e*** is a vector of residuals with an assumed distribution of 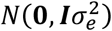.

Then, once the marker effects are estimated (***û***), we can predict breeding values (***â***) for genotyped individuals (even non-phenotyped) as:

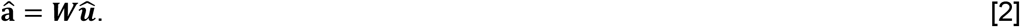

In a bi-parental cross, provided marker data are available on the parents, marker regression coefficients **û** can be partitioned into those that pertain to the favoured alleles of the primary parent **û**_1_ and those that pertain to favoured alleles of the secondary parent **û**_2_ such that **û** = **û**_1_ + **û**_2_. We define the primary parent as the better performing line, the elite parent in an introgression programme. The prediction equation [2] can then be partitioned into:

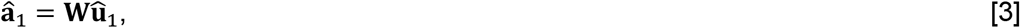

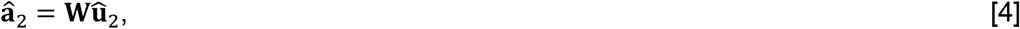

and

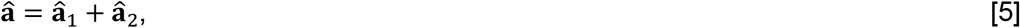

where **â**_1_ is the contribution from the primary parent and analogously **â**_2_ is the contribution from the secondary parent.

Among any set of individuals, we can then select based on **â**, or on any index of **â**_1_ and **â**_2_. Thus, we refer to the former method as GS and the latter method as OSGS. Table 1 provides a simple example of the computation of **â**, **â**_1_ and **â**_2_ in which three favourable alleles out of ten are contributed by the exotic parent.

**Table 1.**
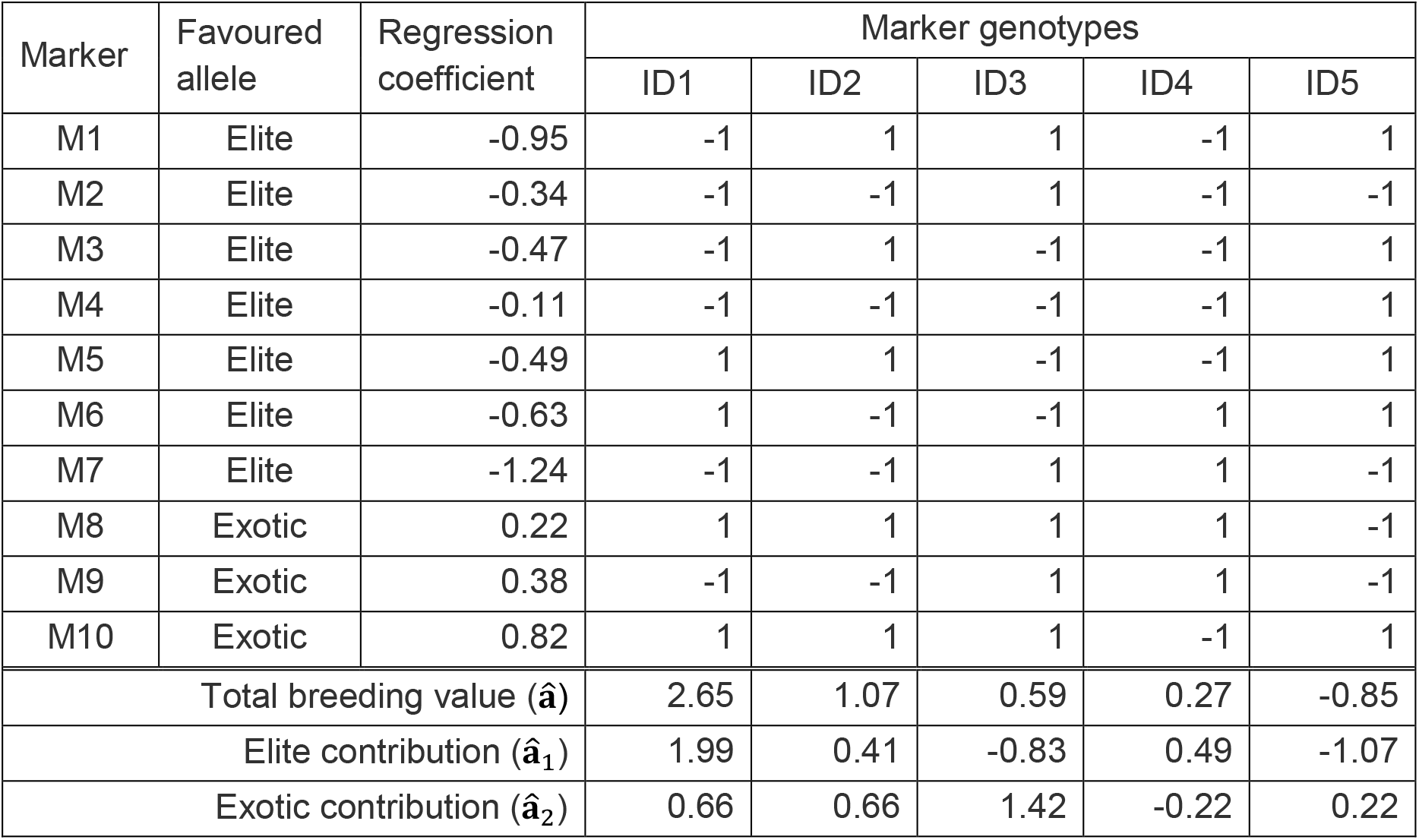
An example of OSGS for ten unlinked markers segregating among inbred lines derived from the F_2_ cross of an elite and exotic parent. At each marker, elite and exotic homozygotes are respectively coded −1 and +1. Negative regression coefficient indicates the increasing allele for the trait is carried by the elite parent and a positive coefficient that the increasing allele is carried by the exotic parent. Here, seven favourable alleles originate from the elite and three from the exotic parent. For each individual (ID1-5), the sum of the products of marker genotypes and regression coefficients gives an estimate of the total breeding value, **â**. Totalling products over the first three and last seven markers partitions the breeding value into contribution respectively from the elite (**â**_1_) and exotic (**â**_2_) parent. For the coefficients given, the expected correlation between **â** and **â**_1_ is 0.89 and between **â** and **â**_2_ is 0.45. The expected correlation between **â**_1_ and **â**_2_ is zero, since the markers are not linked in this example.

| Marker | Favoured allele | Regression coefficient | Marker genotypes |  |  |  |  |
| --- | --- | --- | --- | --- | --- | --- | --- |
|  |  |  | ID1 | ID2 | ID3 | ID4 | ID5 |
| M1 | Elite | -0.95 | -1 | 1 | 1 | -1 | 1 |
| M2 | Elite | -0.34 | -1 | -1 | 1 | -1 | -1 |
| M3 | Elite | -0.47 | -1 | 1 | -1 | -1 | 1 |
| M4 | Elite | -0.11 | -1 | -1 | -1 | -1 | 1 |
| M5 | Elite | -0.49 | 1 | 1 | -1 | -1 | 1 |
| M6 | Elite | -0.63 | 1 | -1 | -1 | 1 | 1 |
| M7 | Elite | -1.24 | -1 | -1 | 1 | 1 | -1 |
| M8 | Exotic | 0.22 | 1 | 1 | 1 | 1 | -1 |
| M9 | Exotic | 0.38 | -1 | -1 | 1 | 1 | -1 |
| M10 | Exotic | 0.82 | 1 | 1 | 1 | -1 | 1 |
| Total breeding value ($\hat{a}$) | | | 2.65 | 1.07 | 0.59 | 0.27 | -0.85 |
| Elite contribution ($\hat{a}_1$) | | | 1.99 | 0.41 | -0.83 | 0.49 | -1.07 |
| Exotic contribution ($\hat{a}_2$) | | | 0.66 | 0.66 | 1.42 | -0.22 | 0.22 |

### 2. Data Analysis

No modification of an existing method for genomic prediction is required for OSGS provided the method estimates ***u***. OSGS requires only that (i) allele origins are identified and (ii) marker estimates are partitioned into two classes with favourable alleles carried by the primary or by secondary parent. If marker genotypes are centered and coded −1, 0, 1 with 0 as the heterozygous class, this partition is simply on the basis of the sign of the regression coefficients. Also, identifying allele origin is trivial in plant breeding scenarios with inbred parents.

Performance differences among various genomic prediction methods are generally minimal, especially if predictions are among closely related material over a limited number of generations (Daetwyler et al, 2013). In this paper therefore, we compare only three standard methods: ridge regression (Hoerl and Kennarad 1976) as implemented in rrBLUP (Endelmann 2011), LASSO (Tibshirani 1996) as implemented in glmnet (Friedman et al. 2010), and BayesCπ (Habier et al. 2011) as implemented in BGLR (Perez and de los Campos 2014) to test that OSGS is robust to the choice of method. All three methods are available as R packages and all analyses were performed with R 3.6.0 (R Core Team 2019).

For each method of analysis, we estimated correlation coefficients between the observed 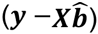 and predicted trait values estimated from (a) all markers (**â**), (b) favourable alleles carried by the primary parent (**â**_1_), and (c) favourable alleles carried by the secondary parent (**â**_2_). The relative importance of the primary and secondary parents as contributors of favourable genetic variation was quantified by the correlations between the pairs of **â**, **â**_1_ and **â**_2_, and by the number and distribution of favourable marker effects among the two parents.

In genomic selection, regression coefficients are typically estimated from one population of lines, the training or reference population, and the prediction equation is applied to a set of candidates with no trait information. Here, the emphasis is different since we are primarily interested in partitioning the observed phenotype of individuals into unobserved contributions from the two parents. For the data analysis here, we partitioned the barley and maize NAMs into training and testing populations as summarised in Table 2. Type 1 ignores the variations among the 25 families. Type 2 accounts for the variations among families by the inclusion of a fixed effect. Type 3 tests the prediction accuracy in the family that was excluded from the training population. We included the type 3 to test if, in addition to partitioning favourable contributions between the two parents, we may also identify which future crosses to make solely from parental information. Type 4 treats each of the 25 crosses separately. For types 3 and 4, we repeated each analysis 25 times: once for each target family.

**Table 2.** Partition of NAMs into training and testing populations to evaluate prediction accuracy from OSGS.

| Type | Number of Families |  | Family (Fixed Effect) |
| --- | --- | --- | --- |
|  | Training Pop. | Testing Pop. |  |
| 1 | 25 | 25 | No |
| 2 | 25 | 25 | Yes |
| 3 | 24 | 1 | No |
| 4 | 1 | 1 | N/A |

In addition, we compared the distributions of favourable primary and secondary marker effects. We first extracted the favourable primary and secondary marker effects based on the signs of rrBLUP coefficients and the favourable direction for each trait. We then converted the marker effects into absolute values and compared the two distributions using the Kolmogorov-Smirnov test as implemented in the *ks.test* function in R (R Core Team 2019). We showed the results as −log_10_(p).

### 3. Barley NAM population

We analysed two polygenic traits in the barley NAM population: days to heading (DTH) and yield (YLD), which were respectively taken from Herzig et al. (2018) and Sharma et al. (2018). Since only raw data on DTH and YLD were provided, we calculated the least squares means of DTH and YLD for 1,420 lines based on the fixed effects of location, nitrogen treatment and year.

We also obtained the accompanying marker genotype data from Maurer et al. (2015), which consisted of 1,427 lines and 5,709 polymorphic markers. We removed five markers that did not map to reference genome, resulting in 5,704 markers. The markers were initially coded as 0 for homozygous elite allele, 1 for heterozygous, 2 for homozygous wild allele and 5 for non-polymorphic within family. To maintain consistency between the barley and maize NAM data, we set all the markers coded as 5 to missing and imputed these missing markers using the same method as for the maize NAM (Buckler et al. 2009), where any missing data were imputed as an average of two non-missing flanking markers. Markers with missing data at the start and end of each chromosome were imputed to be the same as the nearest markers. Finally, we converted the marker from 0/1/2 to −1/0/1 format.

The trait and marker data combined resulted in 1,371 lines for analysis.

### 4. Maize NAM population

We analysed two polygenic traits in the maize NAM population that are comparable to DTH and YLD in the barley NAM population: days to silking (DTS) and cob length (CL), which were taken from Buckler et al. (2009) and Brown et al. (2011) respectively. Instead of calculating the least squares means of DTS and CL, we used the available best linear unbiased predictions of DTS for 4,699 lines and CL for 4,892 lines.

We also obtained the accompanying marker genotype data from McMullen et al. (2009) for 4,699 lines and 1,106 polymorphic markers. The marker data is fully imputed and phased, so we only converted the marker format from 0/1/2 to −1/0/1 format.

The trait and marker data combined resulted in 4,699 lines for analysis.

### 5. Computer simulations

For comparison with the barley and maize NAMs, we simulated three bi-parental populations: (1) F_2_-derived, (2) BC_1_-derived and (3) reverse BC_1_-derived (secondary line as the recurrent parent), all of which were selfed for 4 generations prior to GS/OSGS. For each population, we simulated traits with varying ratios of favourable primary to secondary QTL (P:S = 50:50, 55:45, 60:40, 70:30, 80:20, 90:10) and with QTL densities of 2cM/QTL or 20cM/QTL.

We used rrBLUP to calculate the marker effects in the F_6_, BC_1_S_4_ and rBC_1_S_4_ populations. These were used for determining the breeding values of each line using GS and OSGS methods. For OSGS, we applied five different weighting schemes to favourable alleles originating from the primary or secondary parents (5:5, 6:4, 7:3, 8:2, 9:1). We crossed the top 10 lines (identified by GS/OSGS) in a half diallel and kept 10 progeny from each cross for the next selection cycle using the previously calculated marker effects to predict their breeding values. We repeated this selection process for 5 cycles. Details on the selection process can be found in Supplementary Figure 1. All simulations were repeated 100 times.

All simulations were performed with R 3.6.0 (R Core Team 2019), in which marker data were generated using AlphaSim (Faux et al. 2016) and trait data were generated using custom R scripts. For all populations, we simulated diploid individuals with 10 chromosomes and 7,750 markers distributed evenly across a total genetic distance of 1,550 cM. The markers were coded as −1 for the primary parent and 1 for the secondary parent. QTL positions were randomly sampled from a uniform distribution of all markers. QTL effects for the primary and secondary parent alleles were simulated from a half-normal distribution such that the QTL marker variances are equal between primary and secondary parent alleles, and the aggregated QTL marker variance is equal to *p^-1^*, where *p* is the total number of QTL (Supplementary Figure 2A). Markers selected as QTL markers were left in these analyses since their removal with such high marker density would have little effect, and our purpose is to compare the performance of OSGS and GS and not to test differences in prediction accuracy due to marker-QTL linkage. For any generation, the true breeding value of each line was calculated from its QTL marker genotypes and QTL effects, and the phenotypic trait value of each line was calculated by adding residual value drawn from a standard normal distribution with mean of 0 and variance of 1. Since we fixed the QTL marker variance and residual variance, the simulated trait heritabilities range from 0.40 to 0.95 depending on the proportion of favourable primary and secondary parent alleles and number of QTL markers (Supplementary Figure 2B).

All R scripts and datasets used can be found at doi.org/10.6084/m9.figshare.11343239.

## Results

### 1. Maize and barley NAM data analysis

First, we compared the prediction accuracies of OSGS across four types of analysis (Table 2) and found that, unsurprisingly, OSGS works best when the prediction equation for a testing population contains the same individuals in the training population (Figure 1A-D, Supplementary Figure 3-26). Predictions with either all markers or just the recurrent (primary) parent markers are less variable across the four analysis types than the predictions with the secondary parent markers, which is likely because the 25 secondary parents have different genetic architectures for each trait. However, the prediction accuracies for type 1 and 2 are highly similar, suggesting that having the family as fixed effects was insufficient to account for the variations across families. Overall, best prediction accuracies were observed in type 4, followed by type 2, 1 and 3.

**Figure 1.**
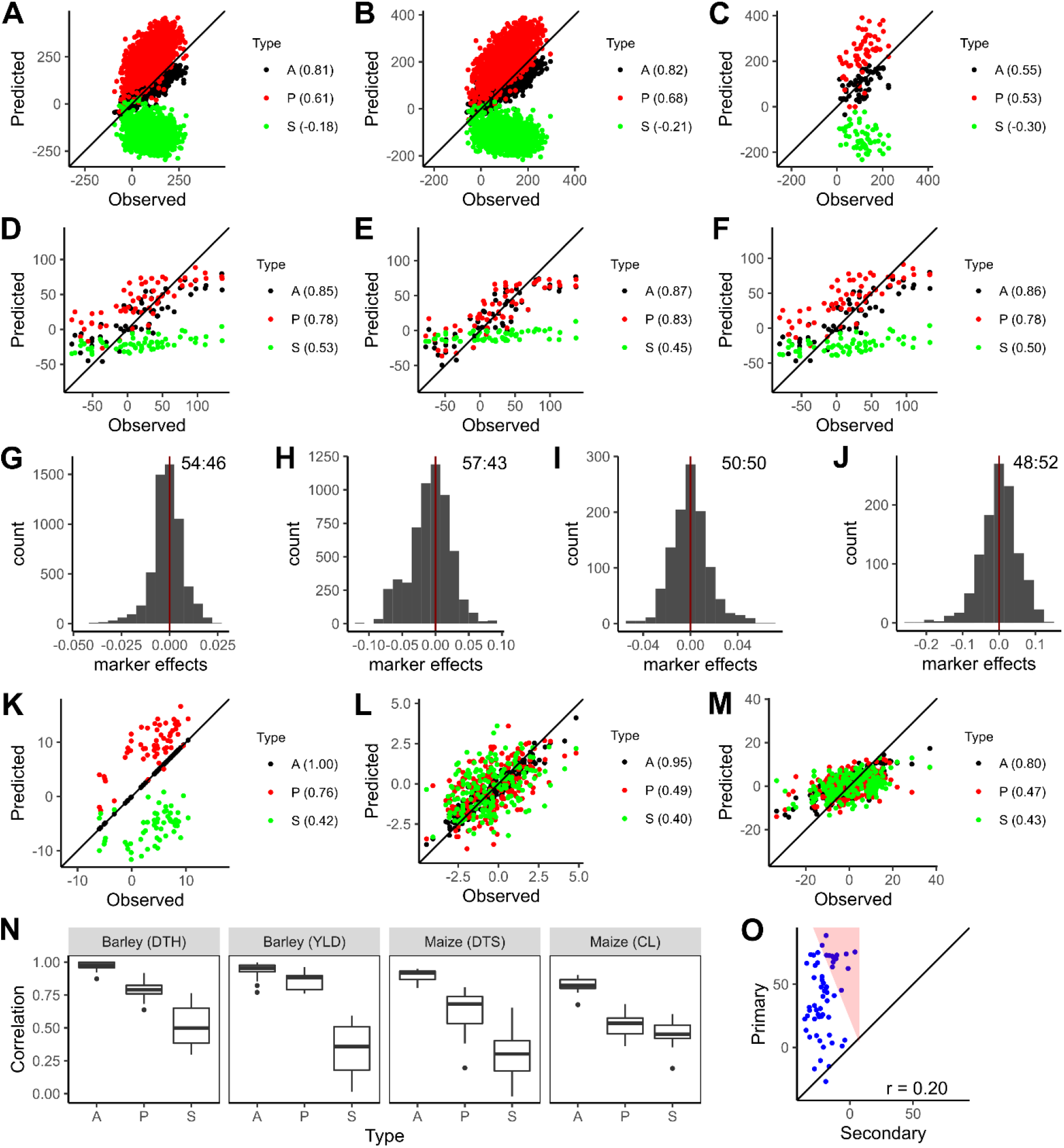
Evaluation of OSGS performances across various considered variables. [**A**] Predictions on YLD with type 1 using all (A), primary (P) and secondary (S) parent markers from rrBLUP. [**B**] Predictions on YLD with type 2 from rrBLUP. [**C**] Predictions on YLD with type 3 on family 1 from rrBLUP. [**D**] Predictions on YLD with type 4 on family 1 from rrBLUP. [**E**] Predictions on YLD with type 4 on family 1 from LASSO. [**F**] Predictions on YLD with type 4 on family 1 from BayesCπ. [**G**] rrBLUP coefficients for DTH with type 4 on family 1. [**H**] rrBLUP coefficients for YLD with type 4 on family 1. [**I**] rrBLUP coefficients for DTS with type 4 on family 1. [**J**] rrBLUP coefficients for CL with type 4 on family 1. [**K**] Predictions on DTH with type 4 on family 1 from rrBLUP. [**L**] Predictions on DTS with type 4 on family 1 from rrBLUP. [**M**] Predictions on CL with type 4 on family 1 from rrBLUP. [**N**] Boxplots of all 25 correlations from each family for each trait and prediction source. [**O**] P-vs-S predictions on yield with type 4 on family 1 from rrBLUP, and the highlighted red triangle represents an equal weight index selection between P and S parent predictions.

OSGS is robust to the choice of GS methods as shown using three popular GS methods (rrBLUP, LASSO and BayesCπ) (Figure 1D-F, Supplementary Figure 3-26). There are little differences in prediction accuracies across these methods, and this holds true even when the markers are partitioned into favourable primary and secondary classes. However, there are a few cases where the training population size is small and the LASSO failed to identify any favourable secondary parent alleles, resulting in zero prediction from these alleles (Supplementary Figure 14).

Distribution of marker effect estimates can inform about the proportion of favourable alleles contributed by each parent (Figure 1G-J, Supplementary Figure 27-38). Late flowering in temperate environment (northern Europe) and high yield are favoured in spring barley, while early flowering and large ear size are favoured in maize, thus favourable DTH, YLD and CL are represented by positive marker effects and favourable DTS is represented by negative marker effects. Across all traits, we found variable proportions of favourable alleles. The 95% confidence intervals for the percentages of favourable primary parent alleles across all 25 families estimated from rrBLUP are 50.4 – 53.6% for barley DTH, 60.4 – 65.5% for barley YLD, 41.7 – 46.2% for maize DTS and 49.6 – 52.7% for maize CL. In barley, we find that the primary (elite) parent has slightly more favourable DTH alleles but many more favourable YLD alleles than the secondary (exotic) parents. In maize, we find that the primary parent has fewer favourable DTS alleles but about equal favourable CL alleles compared to the secondary parents. Provided that a trait fits the infinitesimal model, results here suggest that the distribution of marker effects can be used as a reasonable approximation to the true proportions of favourable QTLs.

In addition to the difference in the proportion of favourable primary and secondary marker effects, the two distributions are significantly different, especially in the barley NAM population (Figure 2). By comparing the favourable primary and secondary marker effect distributions for each trait and family using a Kolmogorov-Smirnov test, we found that all 25 barley NAM families but only about half of the maize NAM families had significant differences. The strongest difference in the distributions was observed in barley YLD, followed by barley DTH, maize DTS and maize CL. Our results here suggest that the distributions of primary and secondary marker effects are more likely to be different in elite-exotic crosses (barley NAM) than elite-elite crosses (maize NAM).

**Figure 2.**
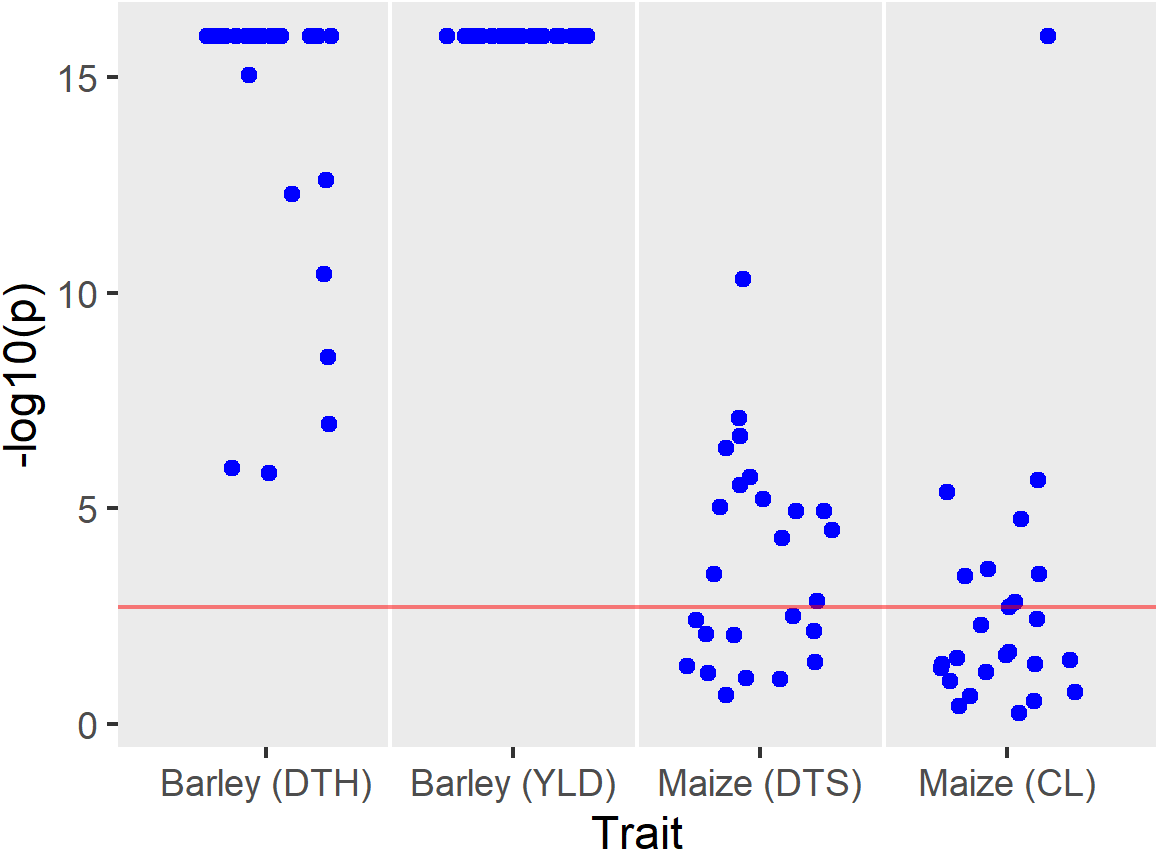
Comparing the distributions of primary (P) and secondary (S) parent marker effects. For each of the 25 NAM families, we separated the rrBLUP coefficients from type 4 into P-favourable and S-favourable, and compared their distributions using Kolmogorov-Smirnov test. The test results are shown as −log_10_(p) and the Bonferroni adjusted threshold of p=0.05 is shown as a red horizontal line.

Across all four traits, the higher the discrepancies in proportion of favourable alleles, the higher the discrepancies in prediction accuracies between primary and secondary parent alleles (Figure 1D, K-N). In the case of YLD in barley, the prediction accuracies from primary parent alleles are much more similar to the prediction accuracies from all markers than from secondary parent alleles, suggesting that the predictions for YLD from all markers are mostly contributed by the primary parent alleles. In contrast, the prediction accuracies for CL in maize are fairly similar between primary and secondary parent alleles, confirming that both primary and secondary parents carry approximately equal proportions of favourable alleles. For the two flowering time traits in barley and maize, the prediction accuracies from primary parent alleles are intermediate between all markers and secondary parent alleles, implying that there is some bias towards primary parent alleles in the predictions but not as severe as observed for YLD.

While a strong positive correlation between the predictions from primary and secondary parent alleles in each family would be ideal for selection, the lack of strong negative correlations (Figure 1O, Supplementary Figure 39-50) implies that we can still select for individuals with good primary and secondary predictions. We can apply index selection based on the ranks of primary and secondary predictions by treating the two predictions as two separate traits. We explored OSGS and index selection further using the simulations described in the next section.

### 2. Computer simulation

First, we compared the effect of QTL density on the performance between GS and OSGS and found that there is more merit to using OSGS when the number of QTLs are large (Figure 3A-B). In the case where the QTL density is low (20cM/QTL), there is little difference between GS and OSGS (Figure 3B). However, as we increased the QTL density to 2cM/QTL, we found that OSGS was able to keep the balance between favourable primary and secondary parent alleles throughout selection, while GS resulted in a larger discrepancy (Figure 3A, Table 3). This highlights the issue with GS in an elite-exotic population as few exotic alleles manage to enter the final breeding population. OSGS can be used to address this issue.

**Figure 3.**
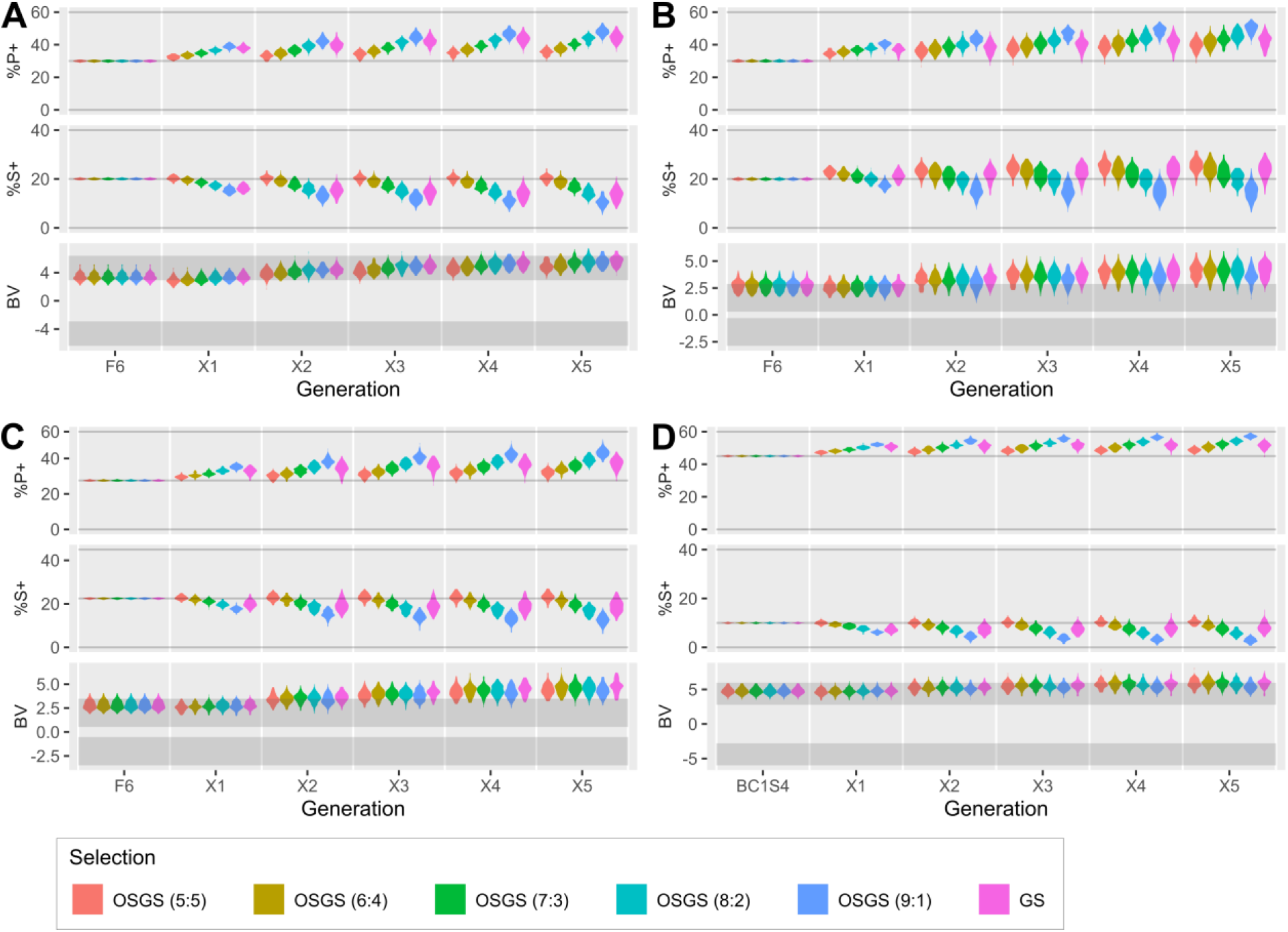
Simulation results comparing OSGS to GS. In each plot, %P+ is the percentage of favourable primary (P) parent alleles, %S+ is percentage of favourable secondary (S) parent alleles, and BV is the breeding values. Densities of %P+ and %S+ are plotted for the mean %P+ and %S+ over 100 simulations. Densities of BV are plotted for the best (maximum) BV over 100 simulations. [**A**] Simulation of an F_2_ population with QTL density of 2cM/QTL and P:S ratio of 60:40. [**B**] Simulation of an F_2_ population with QTL density of 20cM/QTL and P:S ratio of 60:40. [**C**] Simulation of an F_2_ population with QTL density of 2cM/QTL and P:S ratio of 55:45. [**D**] Simulation of a BC_1_ population with QTL density of 2cM/QTL and P:S ratio of 60:40.

**Table 3.**
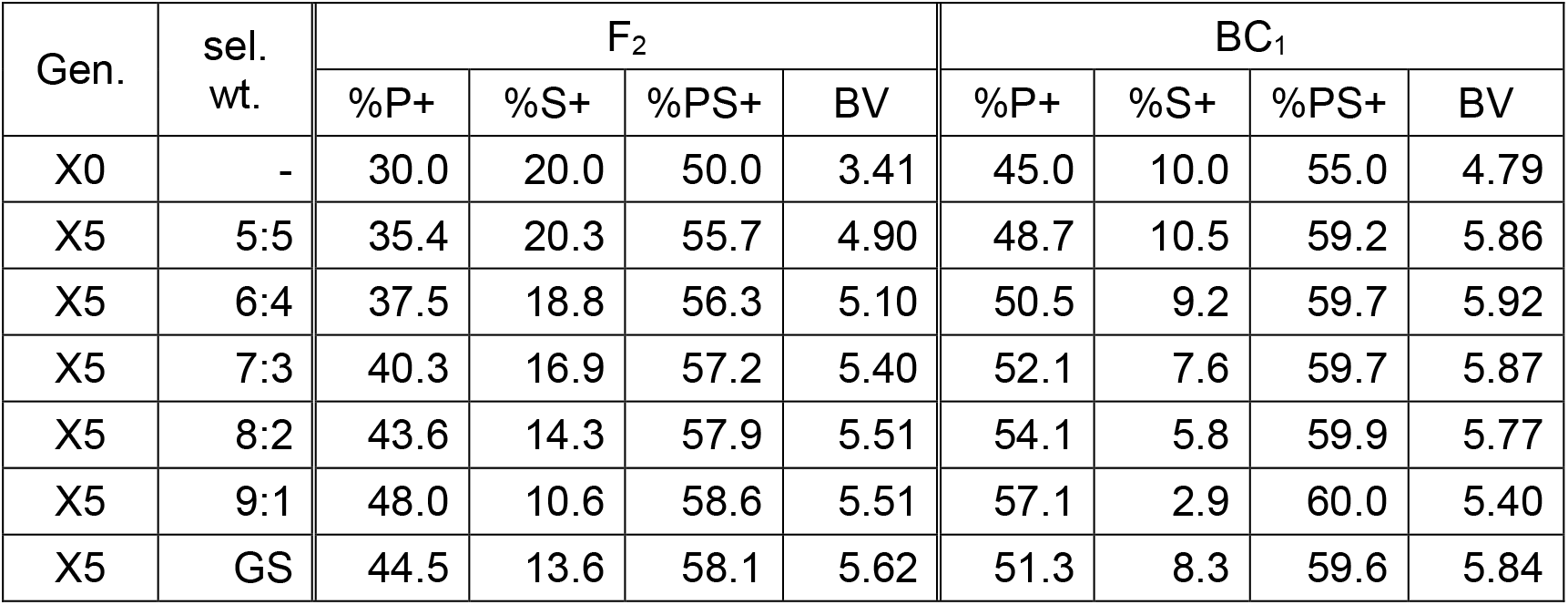
Comparing the changes in proportions of favourable alleles and breeding values depending on starting population type. Means of the percentages of favourable alleles (%P+, %S+, %PS+) and breeding values (BV) from F_2_- and BC_1_-derived populations are calculated and shown for the initial generation (X0 = F_6_ or BC_1_S_4_) and final generation (X5) that underwent OSGS with different selection weights (sel. wt.) or GS. Specifically, in this example, we are showing the simulation results with P:S = 60:40, so the upper limits of %P+ and %S+ are 60% and 40% respectively.

In addition, we found that the performance of OSGS can be optimised depending on the proportions of favourable primary-secondary parent alleles. In the case of a 55:45 QTL proportion (Figure 3C), OSGS resulted in similar breeding values to GS while maintaining a balance between favourable primary and secondary parent alleles. In the case of 60:40 QTL proportion (Figure 3A), if we increase index selection to slightly favour the primary over secondary parent alleles, we can achieve better breeding values at a cost of increasing the discrepancy between favourable primary and secondary parent alleles. Depending on the true QTL proportion, there are specific selection weights for primary and secondary parent that would lead to OSGS being approximately equivalent to GS. The choice of weights for index selection would rest in the user’s preference between allele proportions and breeding values.

Comparing F_2_ and BC_1_-derived populations, OSGS is best performed in an F_2_ population as it begins with an equal proportion of primary and secondary parent alleles (Figure 3A and 3D, Supplementary Figure 51 and 52, Table 3). In a BC_1_ population, there is already a bias in the population towards primary parent alleles as the population has 75% primary parent alleles and 25% secondary parent alleles on average. From a different perspective, without OSGS, one is better off applying GS in a BC_1_ over an F_2_ population as it achieves higher breeding values faster (Table 3). In addition, there is little advantage to implement either OSGS or GS in a backcross population where the exotic parent is used as the recurrent parent as the increase in breeding value comes at a cost of decrease in the proportion of favourable secondary parent alleles (Supplementary Figure 53).

## Discussion

In the recent years, there has been a growing interest in exploring ways for efficient introduction of novel genetic variation from exotic germplasm like landraces and wild relatives into modern breeding populations (Mascher et al. 2019). Even in elite crosses, current selection practices can be strongly biased in favour of one parent (Fradgley 2019), and linkage drag may limit the potential for favourable alleles to be selected from the phenotypically weaker of the two genomes. To circumvent this problem, Gorjanc et al. (2016) suggested an approach to create improved lines from purely exotic materials prior to crossing with the elite materials. Samayoa et al. (2018) suggested a slightly different approach where the exotic improvement is only performed on adaptation-related traits. Alternatively, Han et al. (2017) formulated a method to identify candidate exotic lines for introgressing small numbers of favourable exotic alleles into elite populations. Allier et al. (2019b) further extended this approach for introgressing a larger number of favourable exotic alleles by identifying exotic candidates with higher ratios of favourable over unfavourable alleles. In a slightly different approach, Allier et al. (2019a) proposed the usefulness criterion parental contribution (UCPC) as a metric that combines both the usefulness criterion (Schnell and Utz 1975) and parental genomic contributions in identifying exotic materials for crossing with elite populations. Ru and Bernardo (2019a and 2019b) proposed introgressing linkage groups over QTLs via targeted recombination.

While these approaches seem promising, none of them directly addresses the issues of genomic selection in elite-exotic populations. These approaches focus on identifying the best possible exotic line for crossing, and none attempts to improve the exotic introgression potential after crossing exotic and elite lines. Improvement on solely exotic lines (Gorjanc et al. 2016) may risk selecting for favourable alleles that are already present in elite populations. Selecting for exotic lines with the best combination to the target elite lines (Han et al. 2017; Allier et al. 2019a; Allier et al. 2019b) likely requires accurate predictions on the crosses performances, which calls for large training population and/or close relationships among the selected lines that may not be available.

Here, we propose using OSGS as a generalized framework for partitioning favourable trait contributions among parents. When applied on a single elite-exotic cross population, high prediction accuracies will be possible without requiring a large sized population for phenotyping (Brandariz and Bernardo 2019). This subsequently allows us to partition these predictions into favourable primary and secondary parental contributions with high confidence. OSGS is flexible with respect to the choice of the exotic genome and is complementary to any of the previously described approaches to accommodate those selected exotic lines. In addition, we have demonstrated that OSGS works using the barley and maize NAMs, furthering the potential of community-generated genetic resources as potent breeding tools. Moreover, Bernardo (2009) and our results suggest that it is likely better to use F_2_-derived NAMs to backcross-derived NAMs for this purpose.

In general, OSGS is robust to the choice of a statistical method and should work for other untested methods provided marker effects can be estimated and partitioned into two or more classes. However, one might consider models that are better suited for the presumed trait genetic architectures. For example, LASSO would be a better option for traits regulated by few QTLs since LASSO reduces the effects of most markers to zero.

In this paper we have shown that OSGS can maintain the balance between favourable primary and secondary parent allele proportions over several generations of selection. Hence, OSGS may also play a similar role in optimal contribution selection initially suggested by Meuwissen (1997). Optimal contribution selection aims to maintain genetic diversity in a population under selection by penalizing the estimated breeding values with relationships among selected individuals (Wolliams et al. 2015). In genomic setting this penalty is based on genomic relationships identified from all markers, which does not distinguish between favourable primary and secondary parent alleles. Therefore, OSGS can be complementary to optimal contribution selection as we could partition the kinship matrix into two matrices based on markers carrying favourable primary or secondary parent allele effects. Similar approach has been advocated for optimal contribution selection in rare breeds of livestock in the presence of introgression from cosmopolitan breeds (Wang et al. 2017a, 2017b and 2019).

There are several applications of OSGS remaining to be explored. We found that the distributions of favourable primary and secondary parent effects are different, especially in elite-exotic crosses. This is expected because of the joint action of selection and drift during and after species domestication. OSGS may provide an approach to studying this effect by comparing distributions across populations and species. The application of OSGS could be extended to multi-parental crosses using predictions based on identity-by-descent relationships due to originating parents. Multi-parent populations based on two or more elite lines and a single exotic parent are already in use in pre-breeding (Hao et al. 2019; Singh et al. 2018). There is a strong risk that phenotypic or genomic selection in these populations will discriminate against favourable alleles carried by the exotic parent to an even greater extent than we have shown in bi-parental populations (see also simulations by Gorjanc et al. 2016).

There might also be merits in combining OSGS with other approaches. For example, we can combine the parent selection approaches of Allier et al. (2019b) with OSGS. This may be particularly useful for breeding programs that attempt to use elite and exotic lines with high performance gaps in the traits of interest. In addition, OSGS can be extended to work with gametic variance-based selection (Bijma et al. 2019) by maintaining a balance in the parental contributions on gametic variance.

Lastly, the most promising application of OSGS may be its extension to multi-trait selection. This could be especially useful in elite-exotic crosses where the traits are not unanimously favourable in the elite lines. For example, the exotic parent may carry most favourable alleles for abiotic or biotic stress resistance, but the elite parent mostly for productivity traits.

