## Supplementary Figures 1 - 53 for "Origin Specific Genomic Selection: a simple process to optimize the favourable contribution of parents to progeny"

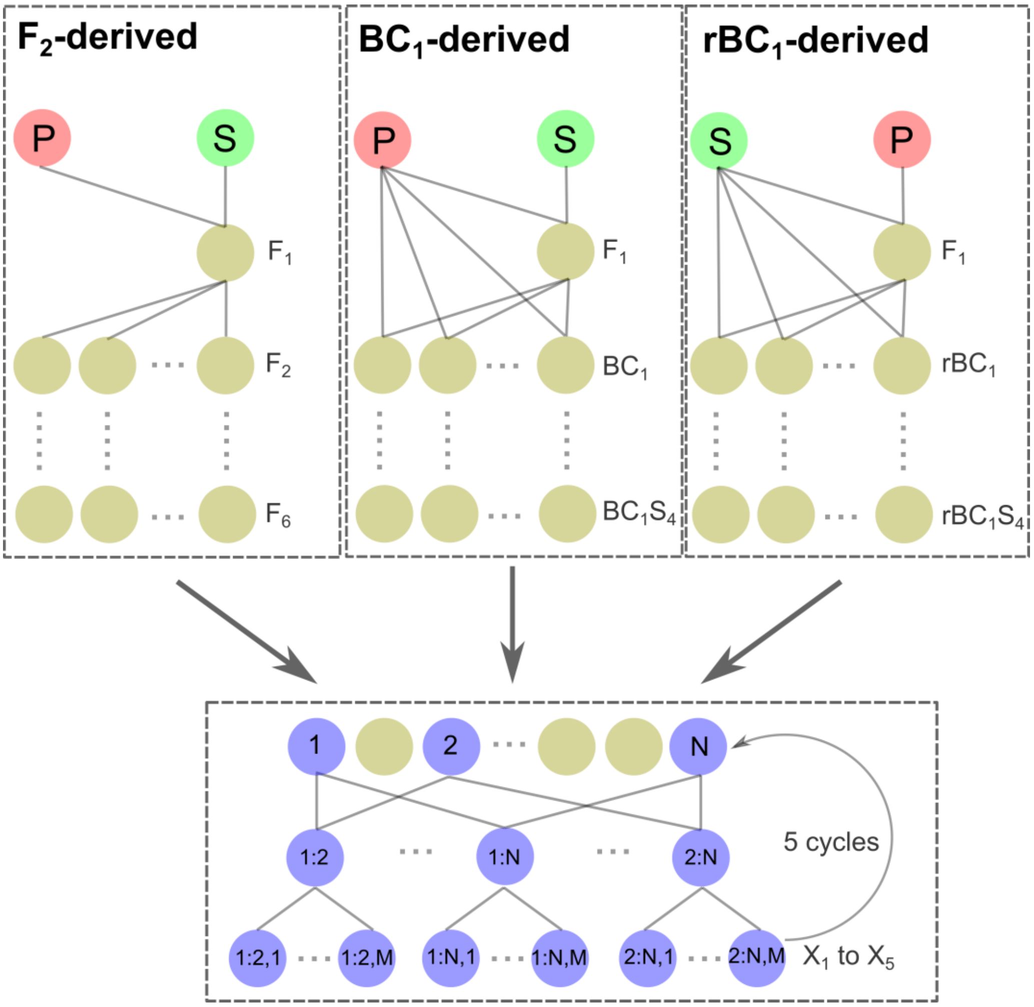


**Supplementary Figure 1: Selection process used in the simulation study**. For each of the $F_2$-, $BC_1$- and $rBC_1$-derived populations, we selected *N* lines based on their breeding values determined from either genomic selection (GS) or origin specific genomic selection (OSGS). All possible half-diallel crosses were made among the *N* selected lines, and *M* progeny were kept from each cross to make up the population referred to as $X_1$, which has $M\sum_i^{N-1} i$ lines that are used for the next cycle of selection. A total of 5 cycles of selection were made, in which the percentage of favourable primary and secondary QTL alleles and the true genetic values for each line were evaluated from $X_1$ to $X_5$. We set *N=10* and *M=10* for simulation with starting population size of 200.

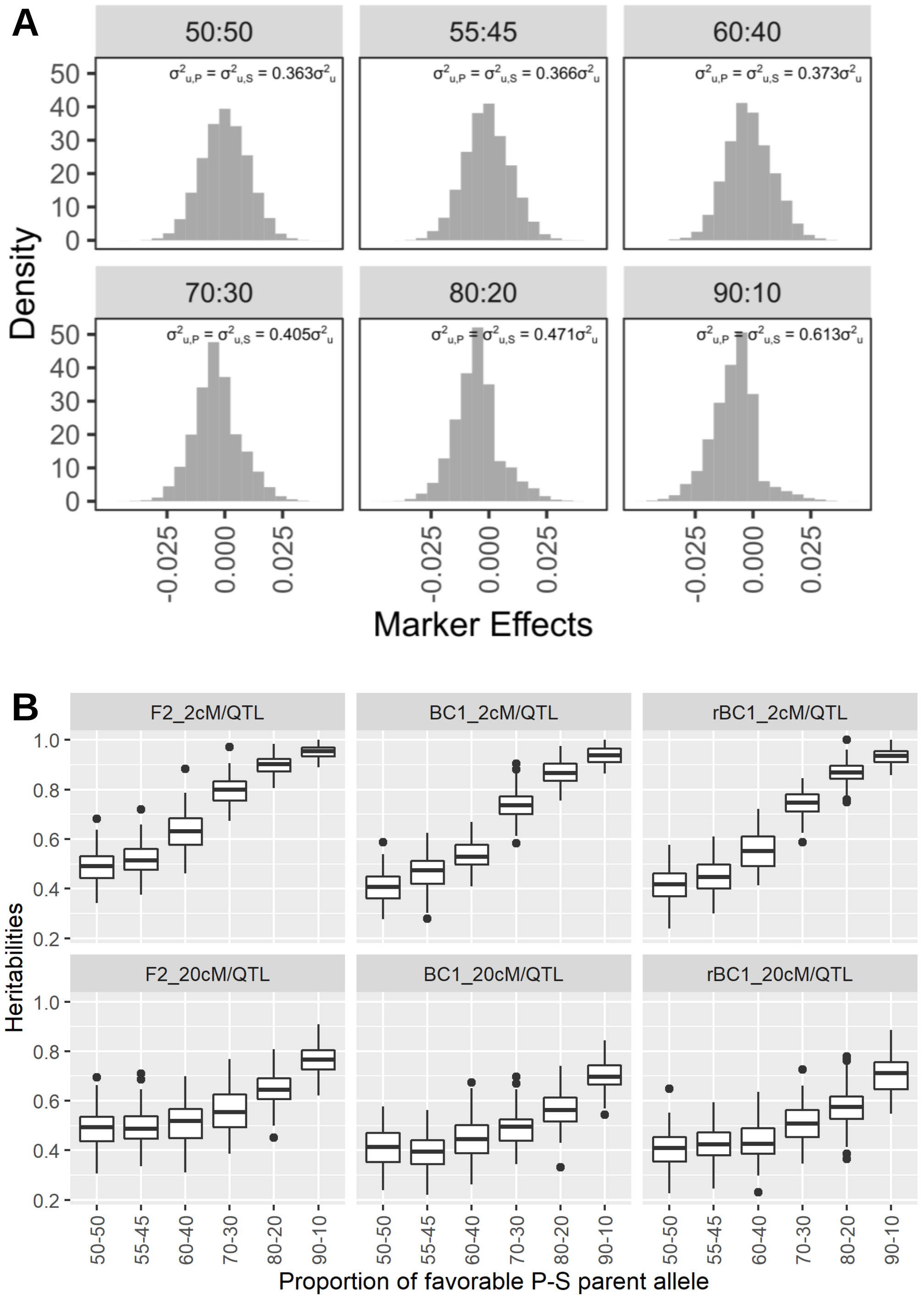


**Supplementary Figure 2. Simulated QTL effect for various P:S**. [**A**] For each P:S, we simulated the QTL marker effects for P and S using a half-normal distribution with the same variance ($\sigma^2_{u,P} = \sigma^2_{u,S}$). In addition, we also restricted the simulated effects such that the aggregated distribution of P and S gives $\sigma^2_u = p^{-1}$, where $p$ is the number of QTLs. This ensures that the genetic variance $\sigma^2_g = 1$ under random mating. [**B**] Actual heritabilities of the simulated phenotypic trait for given population, QTL density, and proportion of favorable P and S alleles.

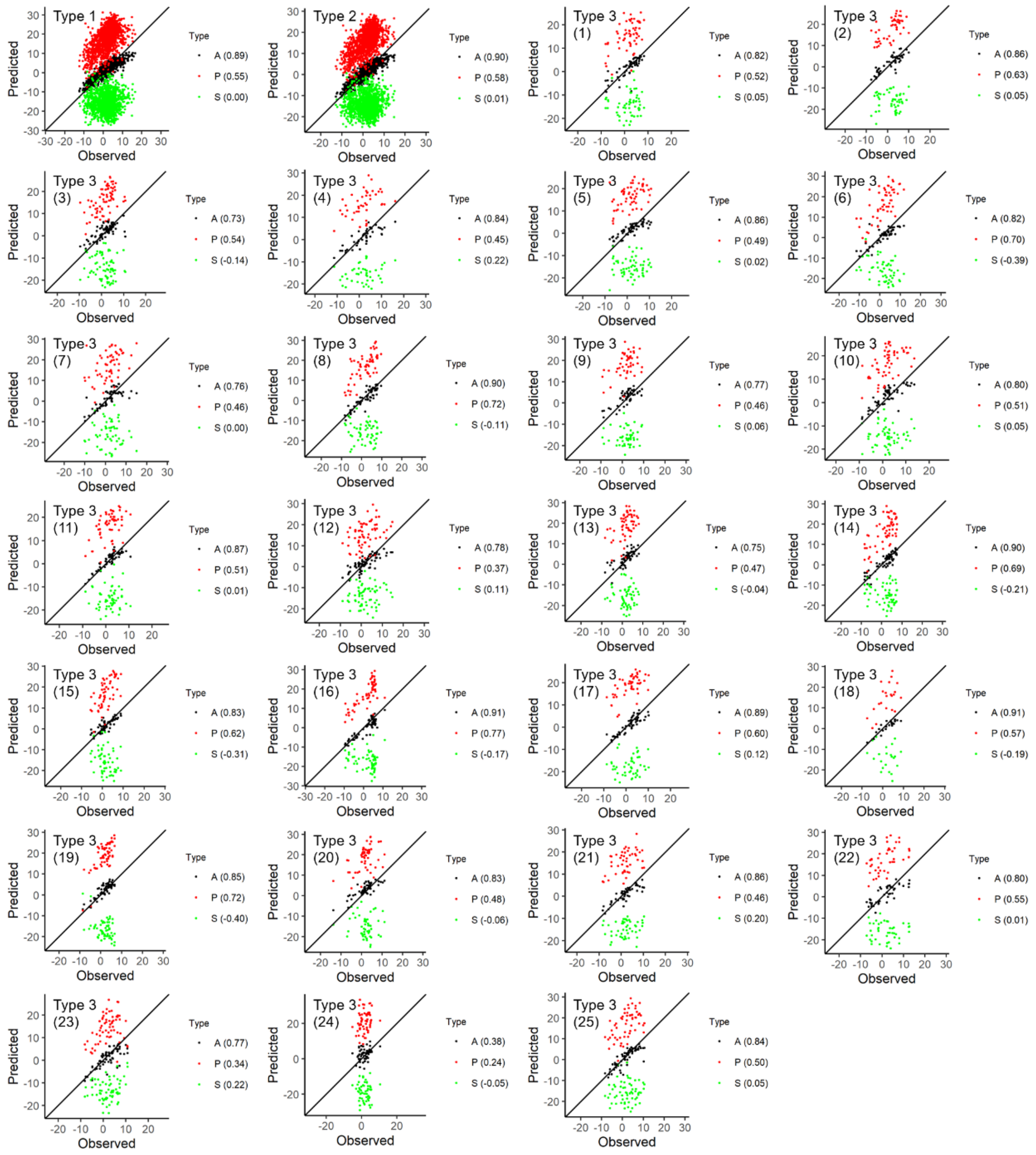


**Supplementary Figure 3. Prediction accuracies on DTH (barley) for analysis type 1-3 using rrBLUP**. Predicted values from all markers (A), favourable primary alleles (P) and favourable secondary alleles (S) are plotted against the observed trait values. Correlations between the predicted and observed values are annotated on each individual plot. Type 1 combines all 25 NAM families, type 2 combines all 25 NAM families while accounting for fixed family effect, and type 3 excludes the testing family in its training set and the testing family is indicated on each individual plot.

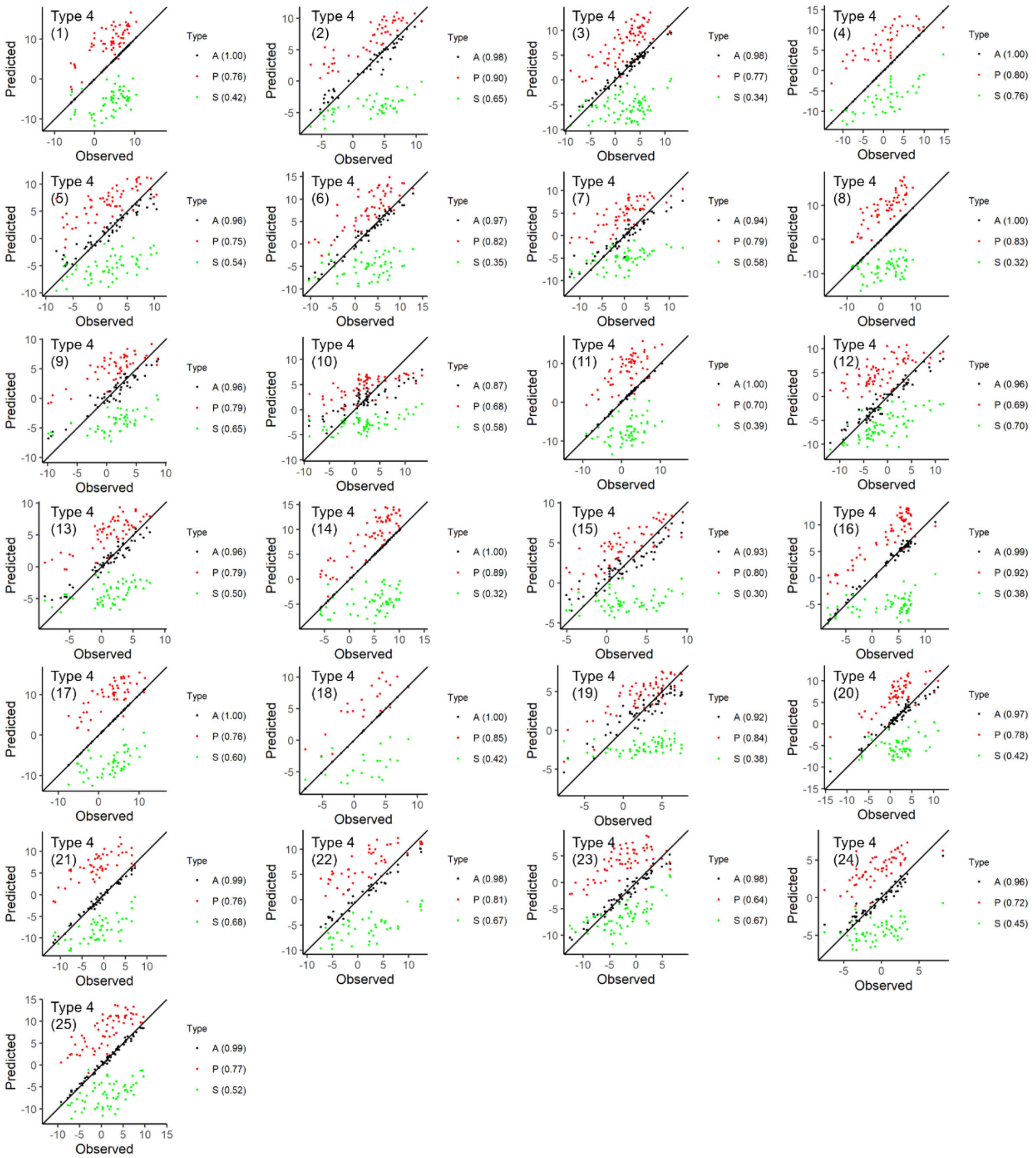


**Supplementary Figure 4. Prediction accuracies on DTH (barley) for analysis type 4 using rrBLUP**. Predicted values from all markers (A), favourable primary alleles (P) and favourable secondary alleles (S) are plotted against the observed trait values. Correlations between the predicted and observed values are annotated on each individual plot. Type 1 combines all 25 NAM families, type 2 combines all 25 NAM families while accounting for fixed family effect, and type 3 excludes the testing family in its training set and the testing family is indicated on each individual plot.

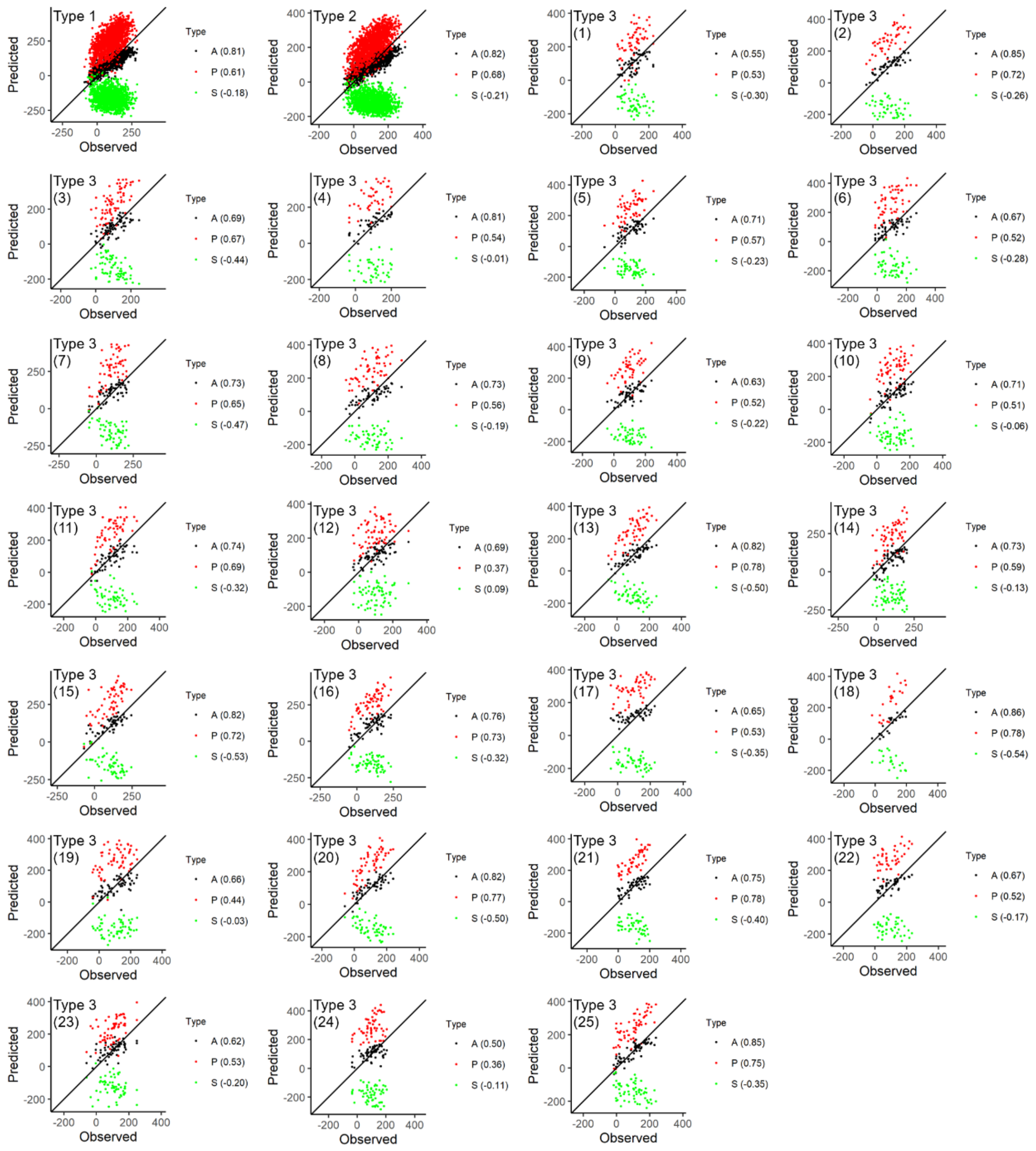

**Supplementary Figure 5. Prediction accuracies on YLD (barley) for analysis type 1-3 using rrBLUP**. Predicted values from all markers (A), favourable primary alleles (P) and favourable secondary alleles (S) are plotted against the observed trait values. Correlations between the predicted and observed values are annotated on each individual plot. Type 1 combines all 25 NAM families, type 2 combines all 25 NAM families while accounting for fixed family effect, and type 3 excludes the testing family in its training set and the testing family is indicated on each individual plot.

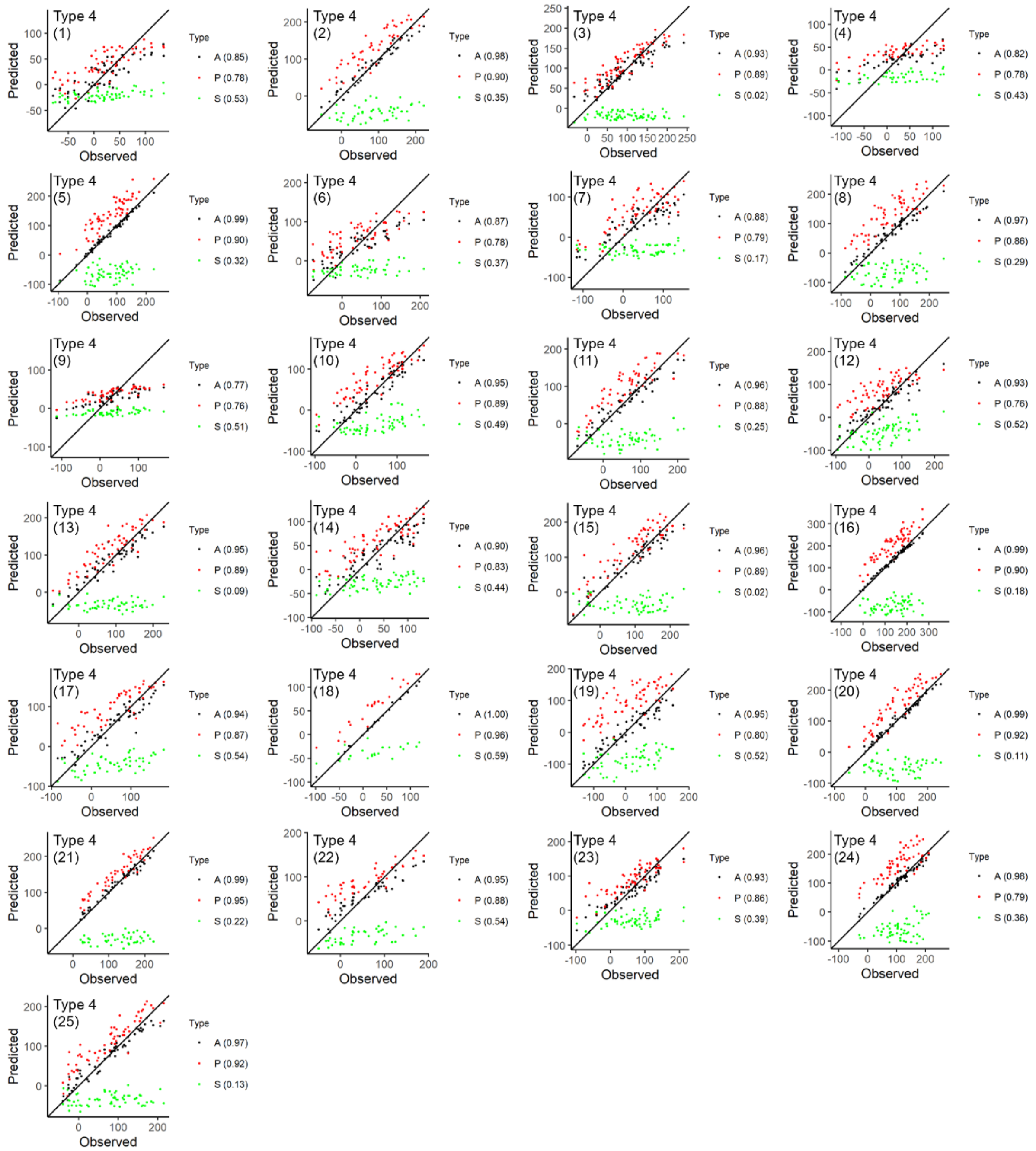


**Supplementary Figure 6. Prediction accuracies on YLD (barley) for analysis type 4 using rrBLUP**. Predicted values from all markers (A), favourable primary alleles (P) and favourable secondary alleles (S) are plotted against the observed trait values. Correlations between the predicted and observed values are annotated on each individual plot. Type 1 combines all 25 NAM families, type 2 combines all 25 NAM families while accounting for fixed family effect, and type 3 excludes the testing family in its training set and the testing family is indicated on each individual plot.

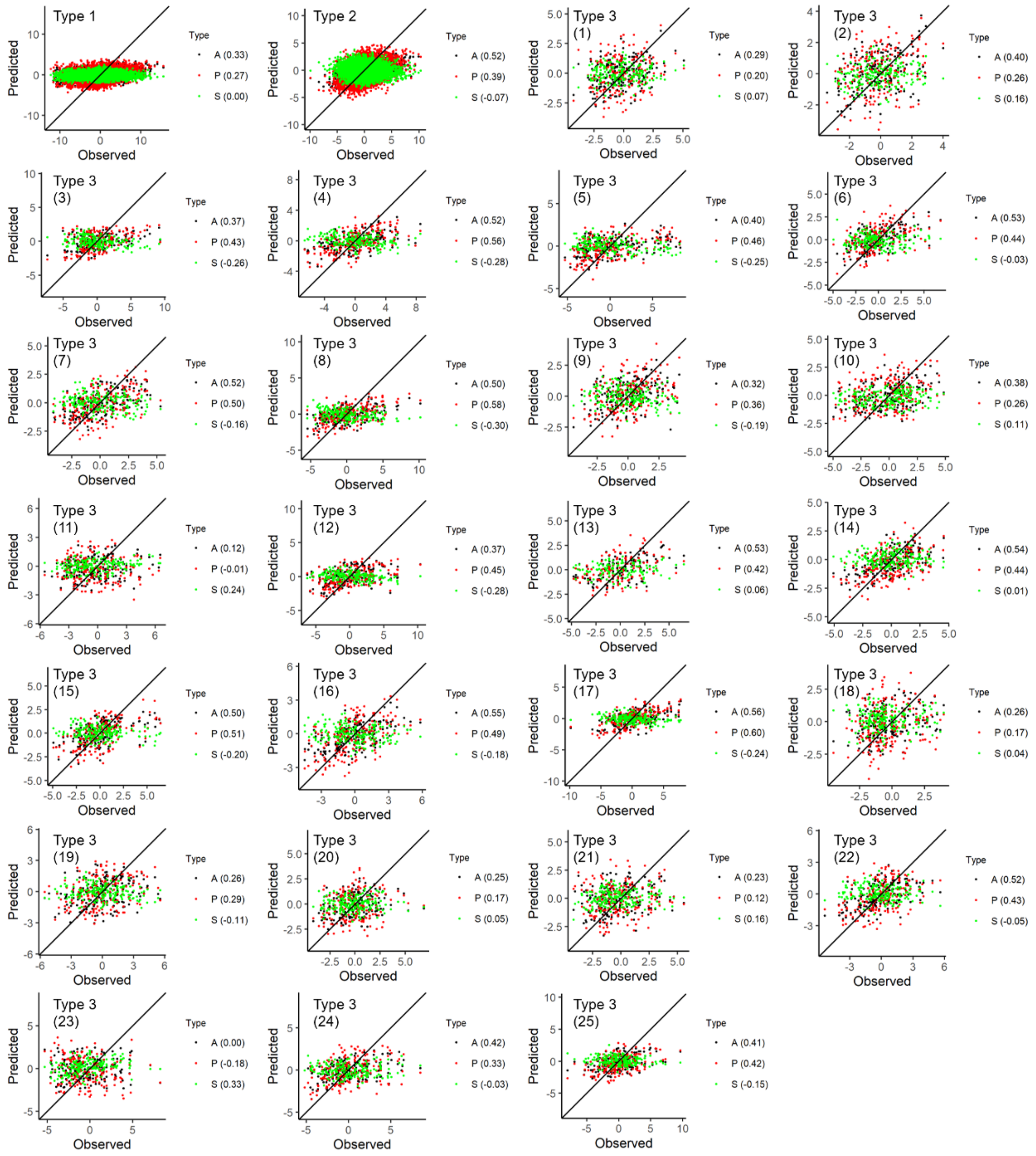


**Supplementary Figure 7. Prediction accuracies on DTS (maize) for analysis type 1-3 using rrBLUP**. Predicted values from all markers (A), favourable primary alleles (P) and favourable secondary alleles (S) are plotted against the observed trait values. Correlations between the predicted and observed values are annotated on each individual plot. Type 1 combines all 25 NAM families, type 2 combines all 25 NAM families while accounting for fixed family effect, and type 3 excludes the testing family in its training set and the testing family is indicated on each individual plot.

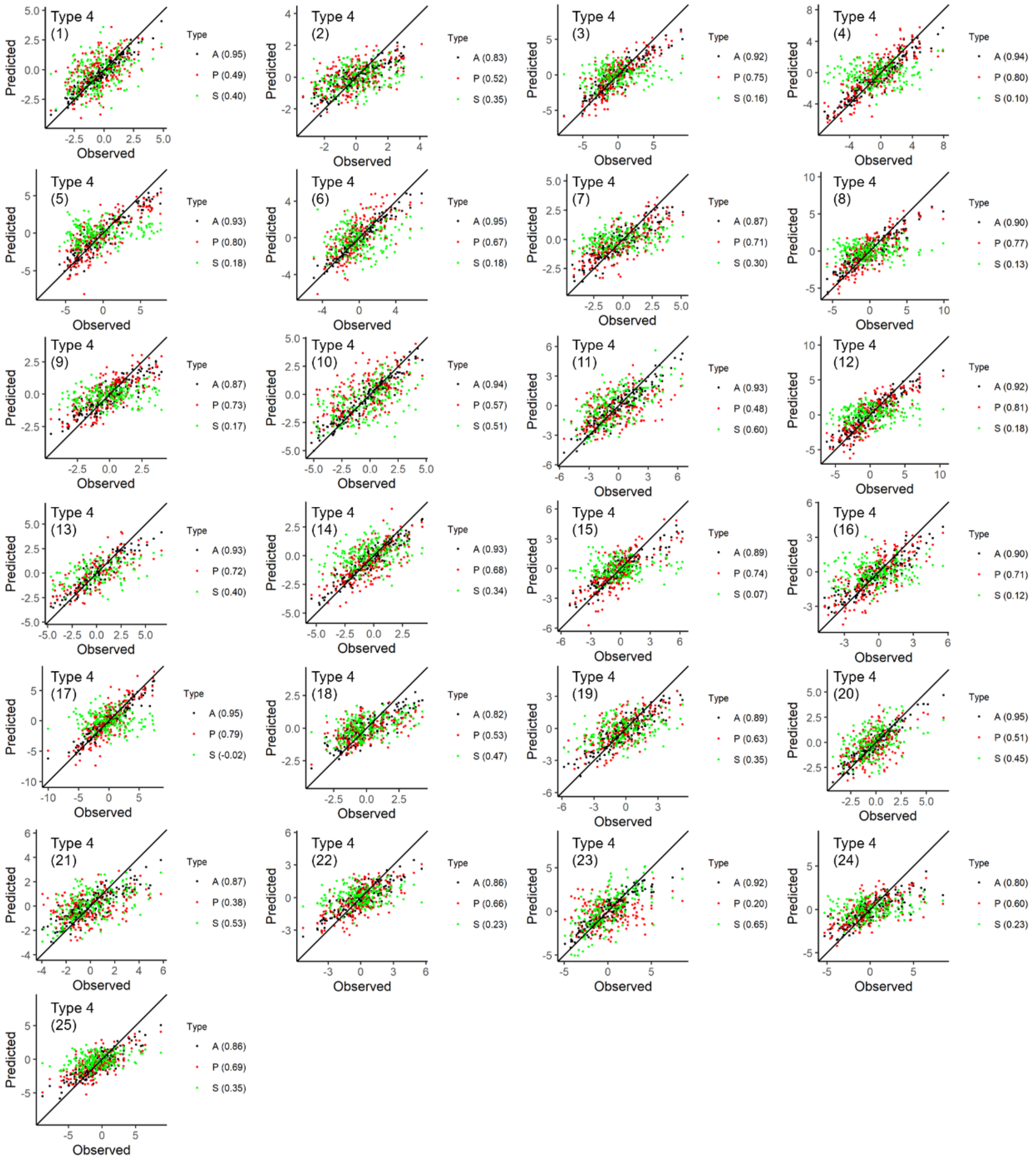


**Supplementary Figure 8. Prediction accuracies on DTS (maize) for analysis type 4 using rrBLUP**. Predicted values from all markers (A), favourable primary alleles (P) and favourable secondary alleles (S) are plotted against the observed trait values. Correlations between the predicted and observed values are annotated on each individual plot. Type 1 combines all 25 NAM families, type 2 combines all 25 NAM families while accounting for fixed family effect, and type 3 excludes the testing family in its training set and the testing family is indicated on each individual plot.

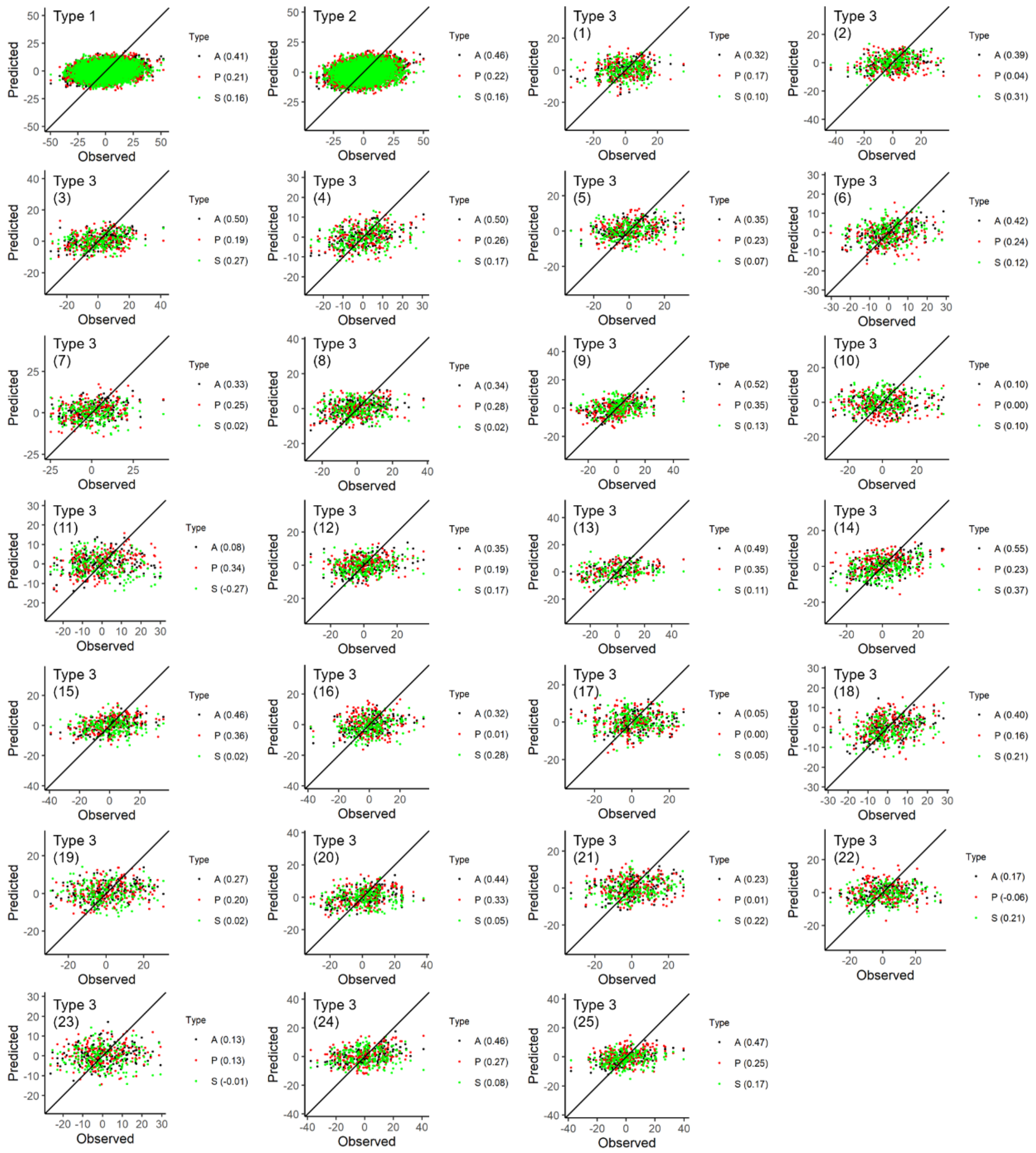


**Supplementary Figure 9. Prediction accuracies on CL (maize) for analysis type 1-3 using rrBLUP**. Predicted values from all markers (A), favourable primary alleles (P) and favourable secondary alleles (S) are plotted against the observed trait values. Correlations between the predicted and observed values are annotated on each individual plot. Type 1 combines all 25 NAM families, type 2 combines all 25 NAM families while accounting for fixed family effect, and type 3 excludes the testing family in its training set and the testing family is indicated on each individual plot.

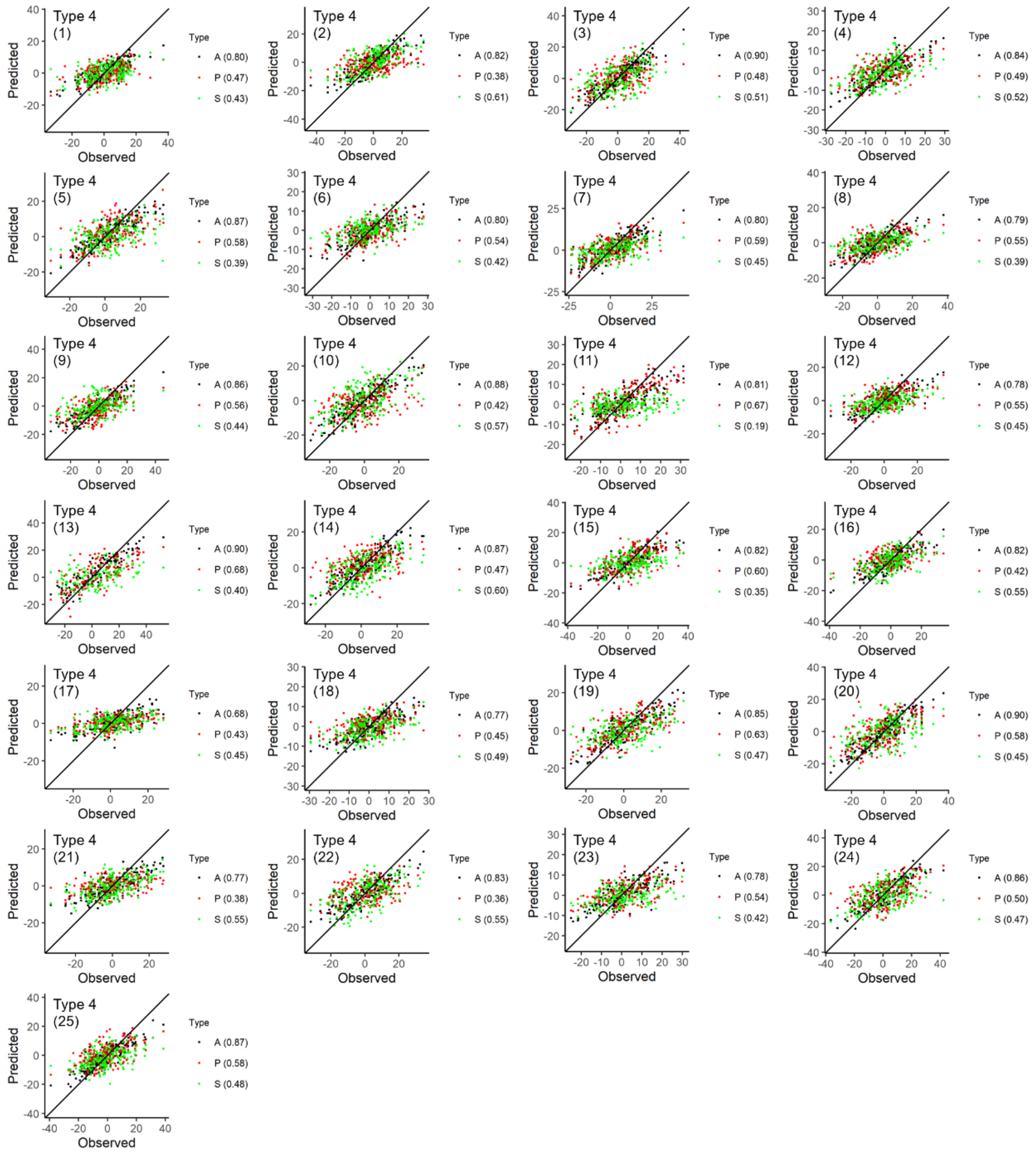


**Supplementary Figure 10. Prediction accuracies on CL (maize) for analysis type 4 using rrBLUP**. Predicted values from all markers (A), favourable primary alleles (P) and favourable secondary alleles (S) are plotted against the observed trait values. Correlations between the predicted and observed values are annotated on each individual plot. Type 1 combines all 25 NAM families, type 2 combines all 25 NAM families while accounting for fixed family effect, and type 3 excludes the testing family in its training set and the testing family is indicated on each individual plot.

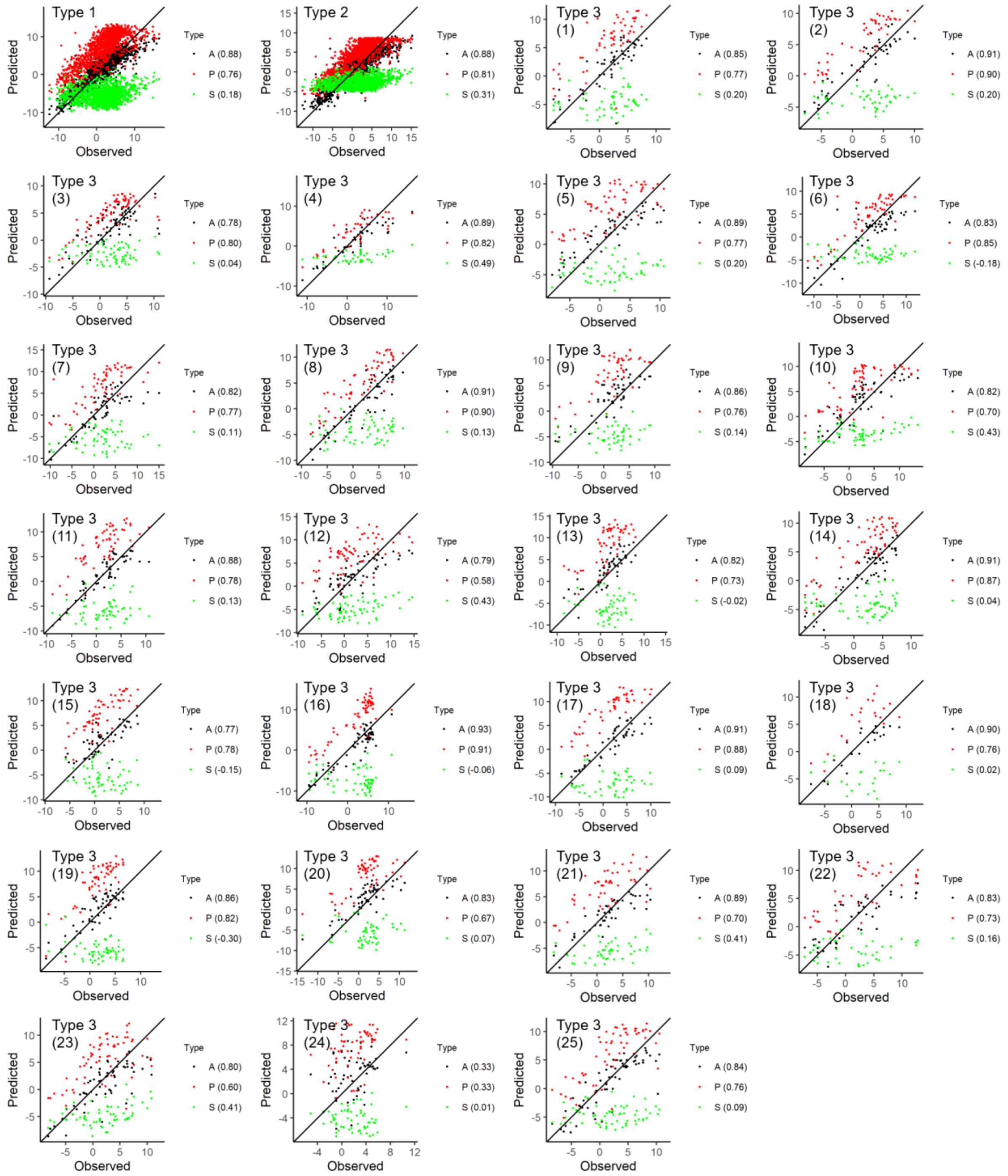


**Supplementary Figure 11. Prediction accuracies on DTH (barley) for analysis type 1-3 using LASSO**. Predicted values from all markers (A), favourable primary alleles (P) and favourable secondary alleles (S) are plotted against the observed trait values. Correlations between the predicted and observed values are annotated on each individual plot. Type 1 combines all 25 NAM families, type 2 combines all 25 NAM families while accounting for fixed family effect, and type 3 excludes the testing family in its training set and the testing family is indicated on each individual plot.

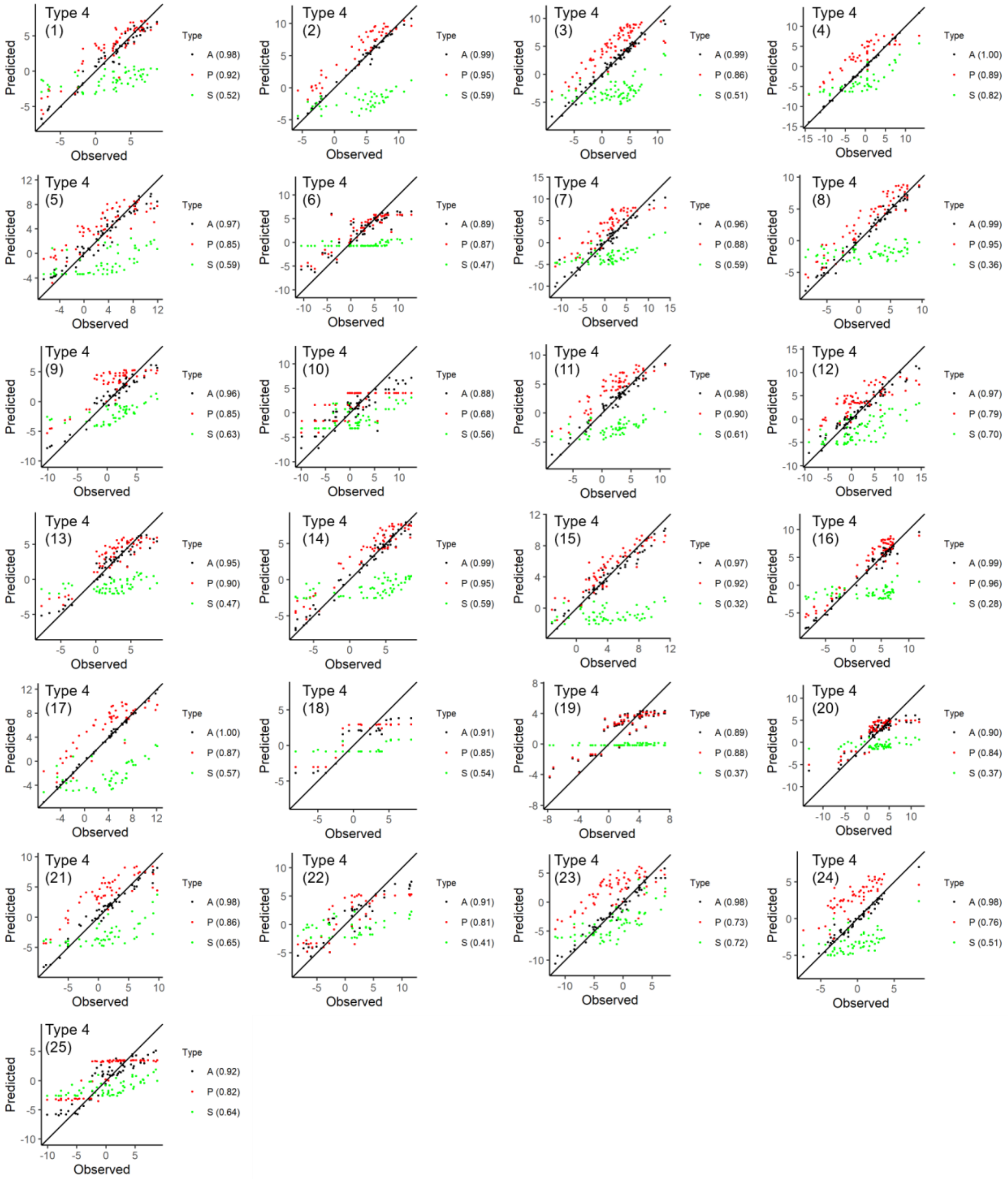


**Supplementary Figure 12. Prediction accuracies on DTH (barley) for analysis type 4 using LASSO**. Predicted values from all markers (A), favourable primary alleles (P) and favourable secondary alleles (S) are plotted against the observed trait values. Correlations between the predicted and observed values are annotated on each individual plot. Type 1 combines all 25 NAM families, type 2 combines all 25 NAM families while accounting for fixed family effect, and type 3 excludes the testing family in its training set and the testing family is indicated on each individual plot.

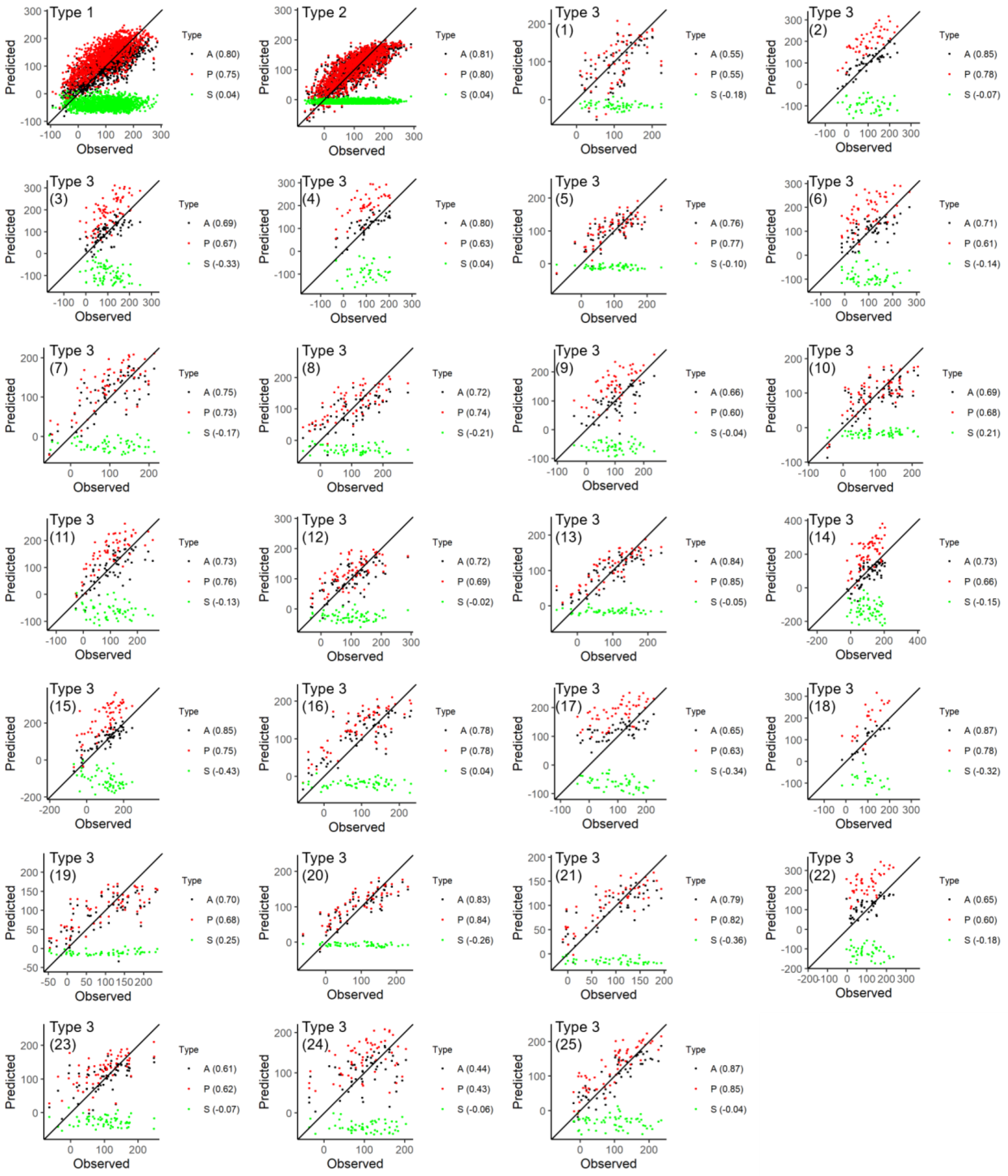


**Supplementary Figure 13. Prediction accuracies on YLD (barley) for analysis type 1-3 using LASSO**. Predicted values from all markers (A), favourable primary alleles (P) and favourable secondary alleles (S) are plotted against the observed trait values. Correlations between the predicted and observed values are annotated on each individual plot. Type 1 combines all 25 NAM families, type 2 combines all 25 NAM families while accounting for fixed family effect, and type 3 excludes the testing family in its training set and the testing family is indicated on each individual plot.

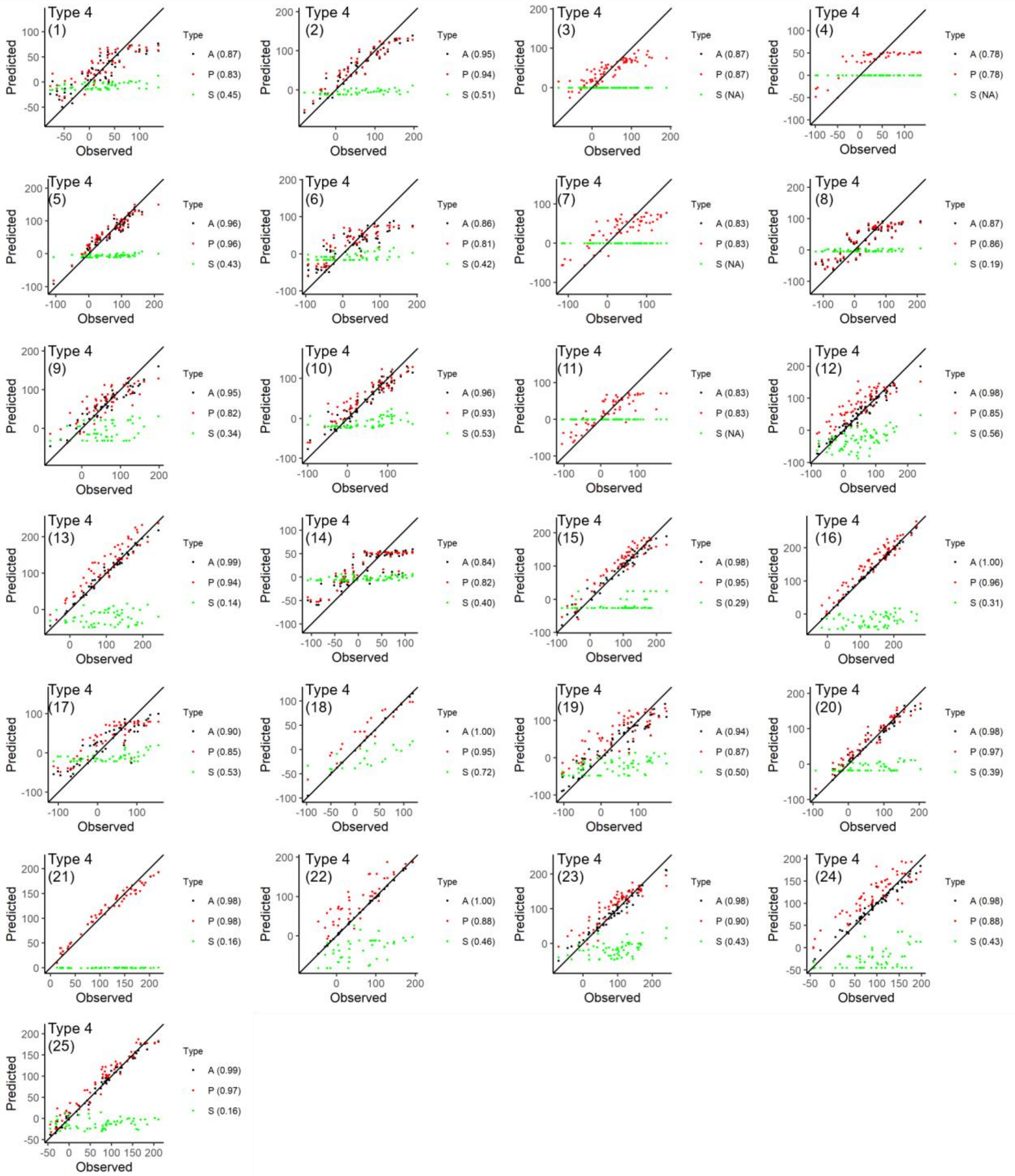


**Supplementary Figure 14. Prediction accuracies on YLD (barley) for analysis type 4 using LASSO**. Predicted values from all markers (A), favourable primary alleles (P) and favourable secondary alleles (S) are plotted against the observed trait values. Correlations between the predicted and observed values are annotated on each individual plot. Type 1 combines all 25 NAM families, type 2 combines all 25 NAM families while accounting for fixed family effect, and type 3 excludes the testing family in its training set and the testing family is indicated on each individual plot.

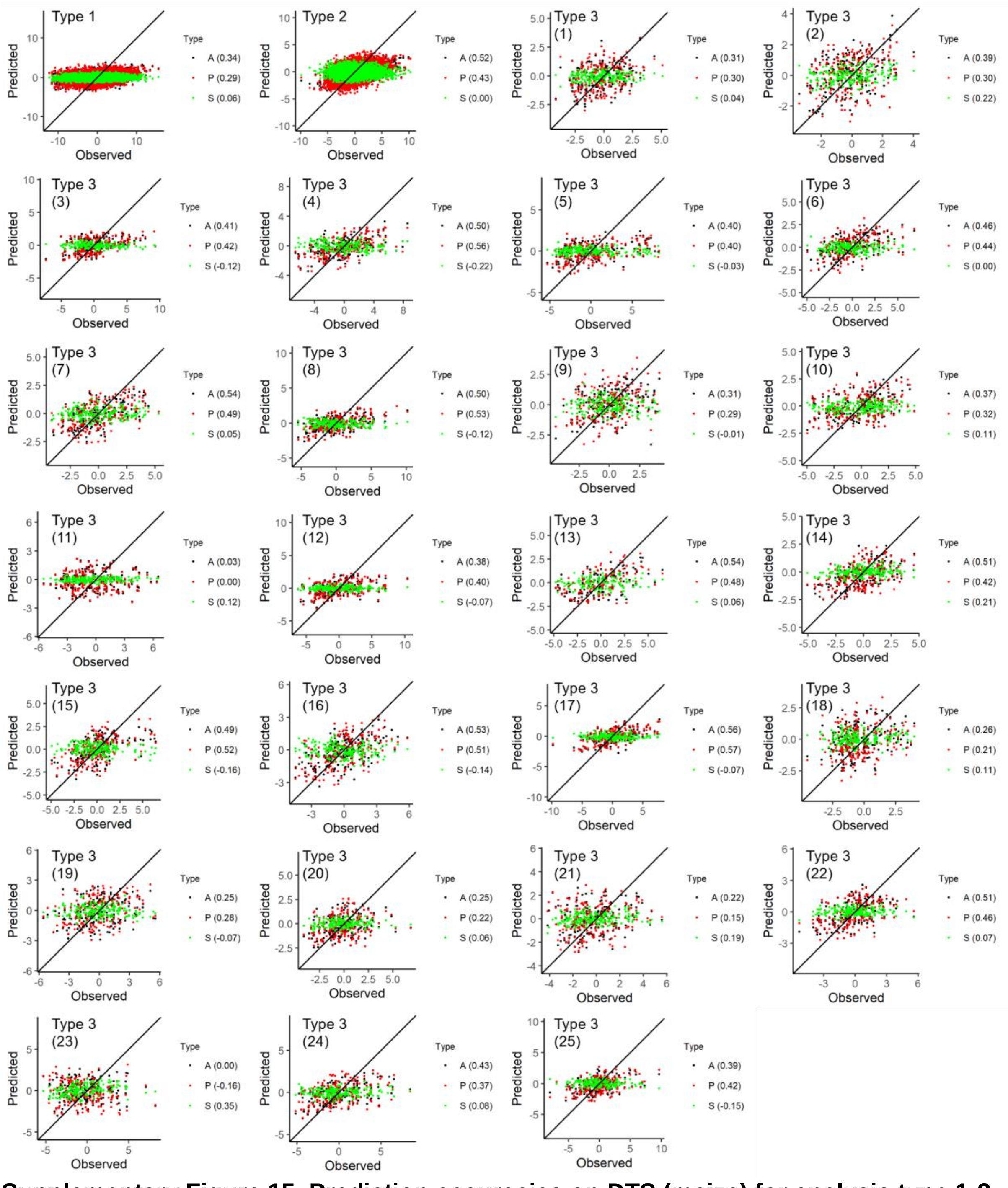


**Supplementary Figure 15. Prediction accuracies on DTS (maize) for analysis type 1-3 using LASSO**. Predicted values from all markers (A), favourable primary alleles (P) and favourable secondary alleles (S) are plotted against the observed trait values. Correlations between the predicted and observed values are annotated on each individual plot. Type 1 combines all 25 NAM families, type 2 combines all 25 NAM families while accounting for fixed family effect, and type 3 excludes the testing family in its training set and the testing family is indicated on each individual plot.

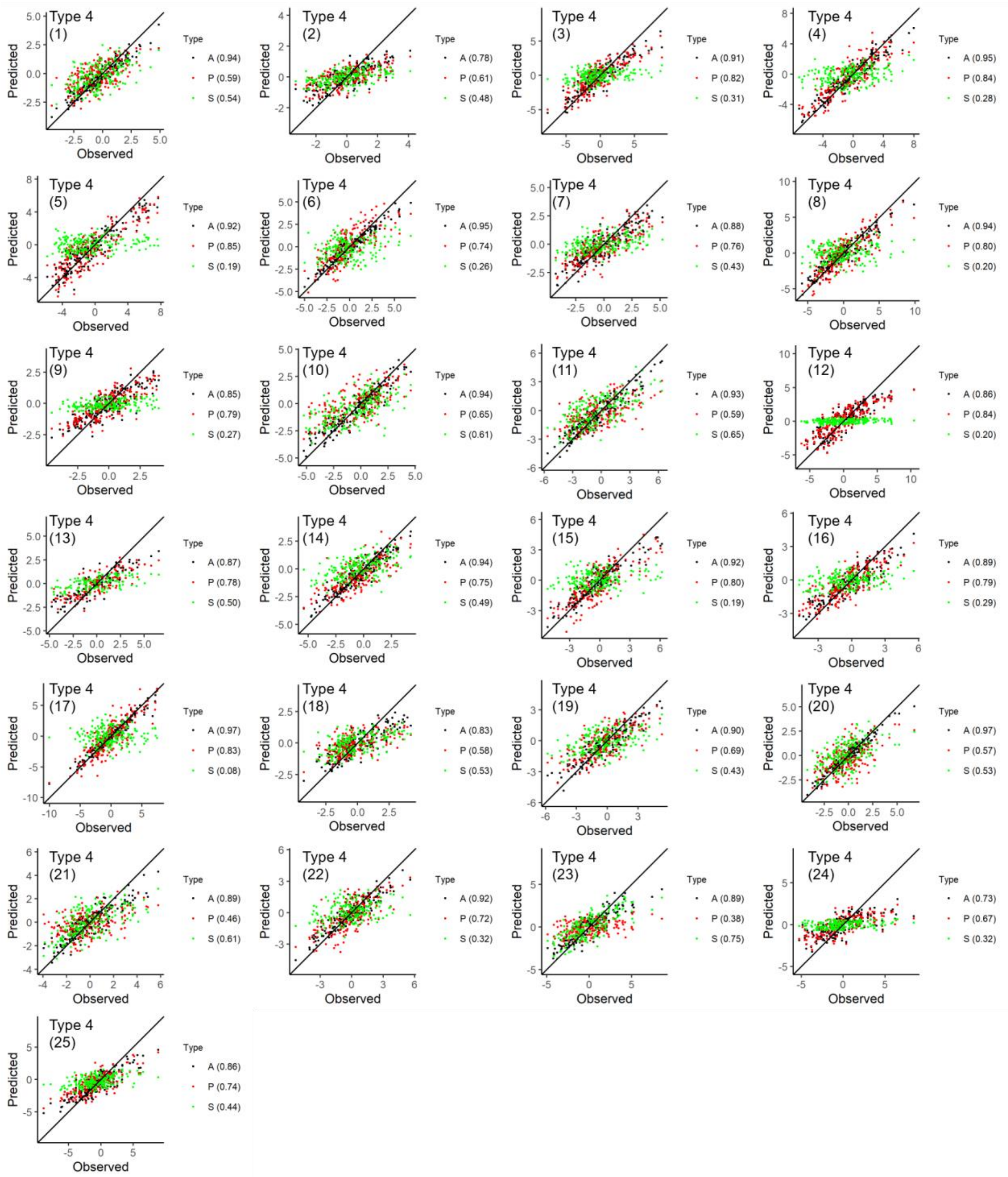


**Supplementary Figure 16. Prediction accuracies on DTS (maize) for analysis type 4 using LASSO**. Predicted values from all markers (A), favourable primary alleles (P) and favourable secondary alleles (S) are plotted against the observed trait values. Correlations between the predicted and observed values are annotated on each individual plot. Type 1 combines all 25 NAM families, type 2 combines all 25 NAM families while accounting for fixed family effect, and type 3 excludes the testing family in its training set and the testing family is indicated on each individual plot.

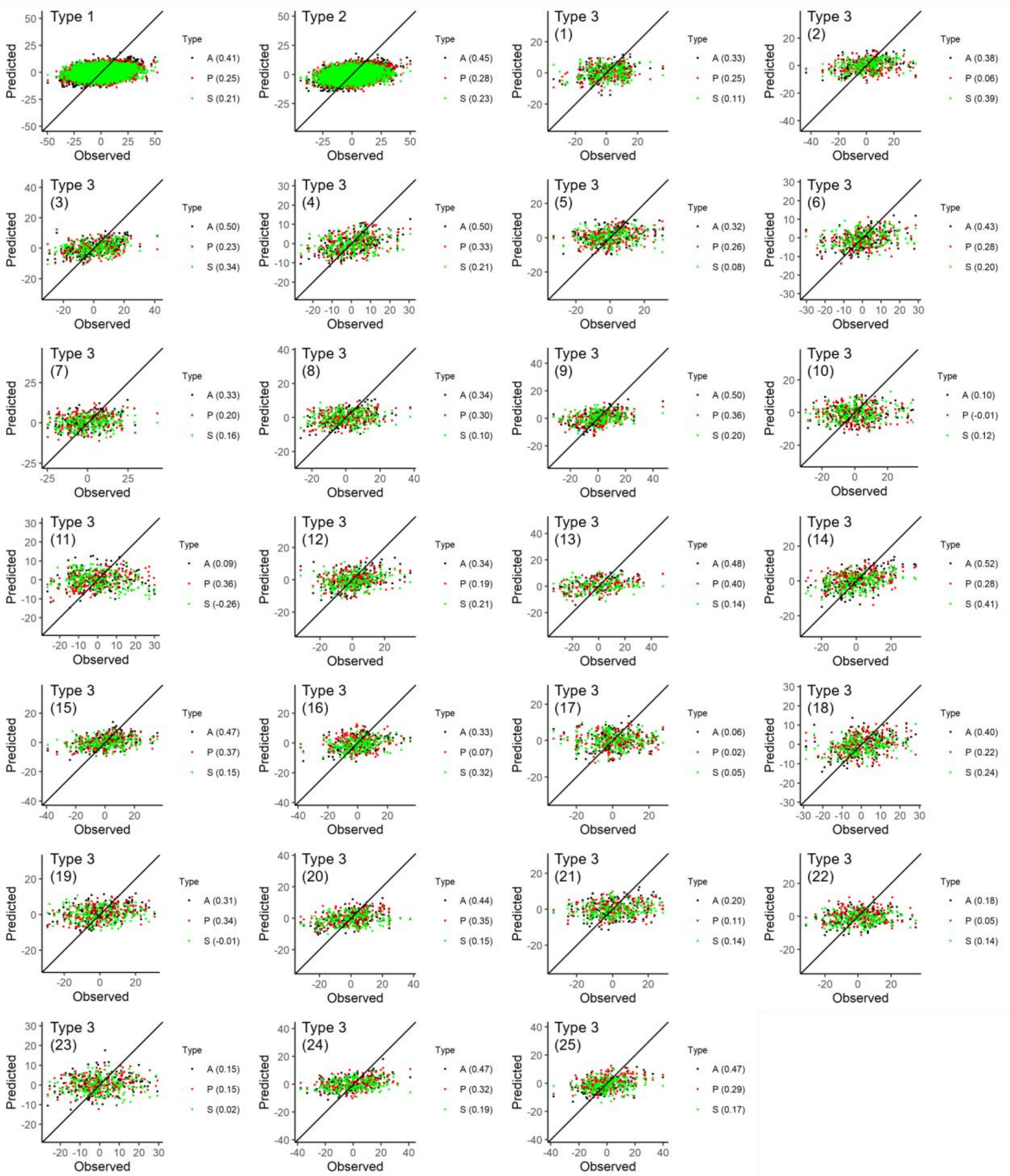


**Supplementary Figure 17. Prediction accuracies on CL (maize) for analysis type 1-3 using LASSO**. Predicted values from all markers (A), favourable primary alleles (P) and favourable secondary alleles (S) are plotted against the observed trait values. Correlations between the predicted and observed values are annotated on each individual plot. Type 1 combines all 25 NAM families, type 2 combines all 25 NAM families while accounting for fixed family effect, and type 3 excludes the testing family in its training set and the testing family is indicated on each individual plot.

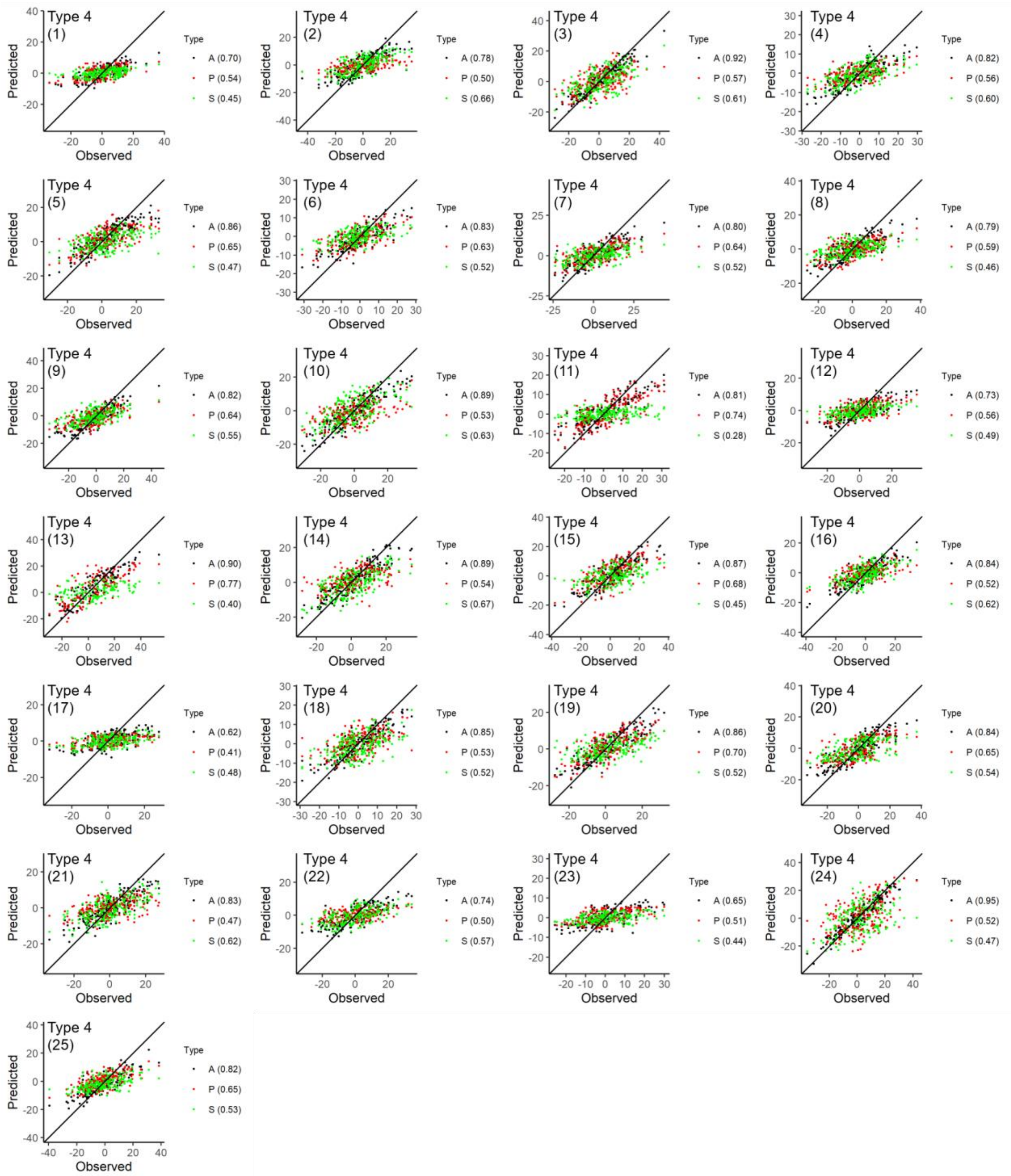


**Supplementary Figure 18. Prediction accuracies on CL (maize) for analysis type 4 using LASSO**. Predicted values from all markers (A), favourable primary alleles (P) and favourable secondary alleles (S) are plotted against the observed trait values. Correlations between the predicted and observed values are annotated on each individual plot. Type 1 combines all 25 NAM families, type 2 combines all 25 NAM families while accounting for fixed family effect, and type 3 excludes the testing family in its training set and the testing family is indicated on each individual plot.

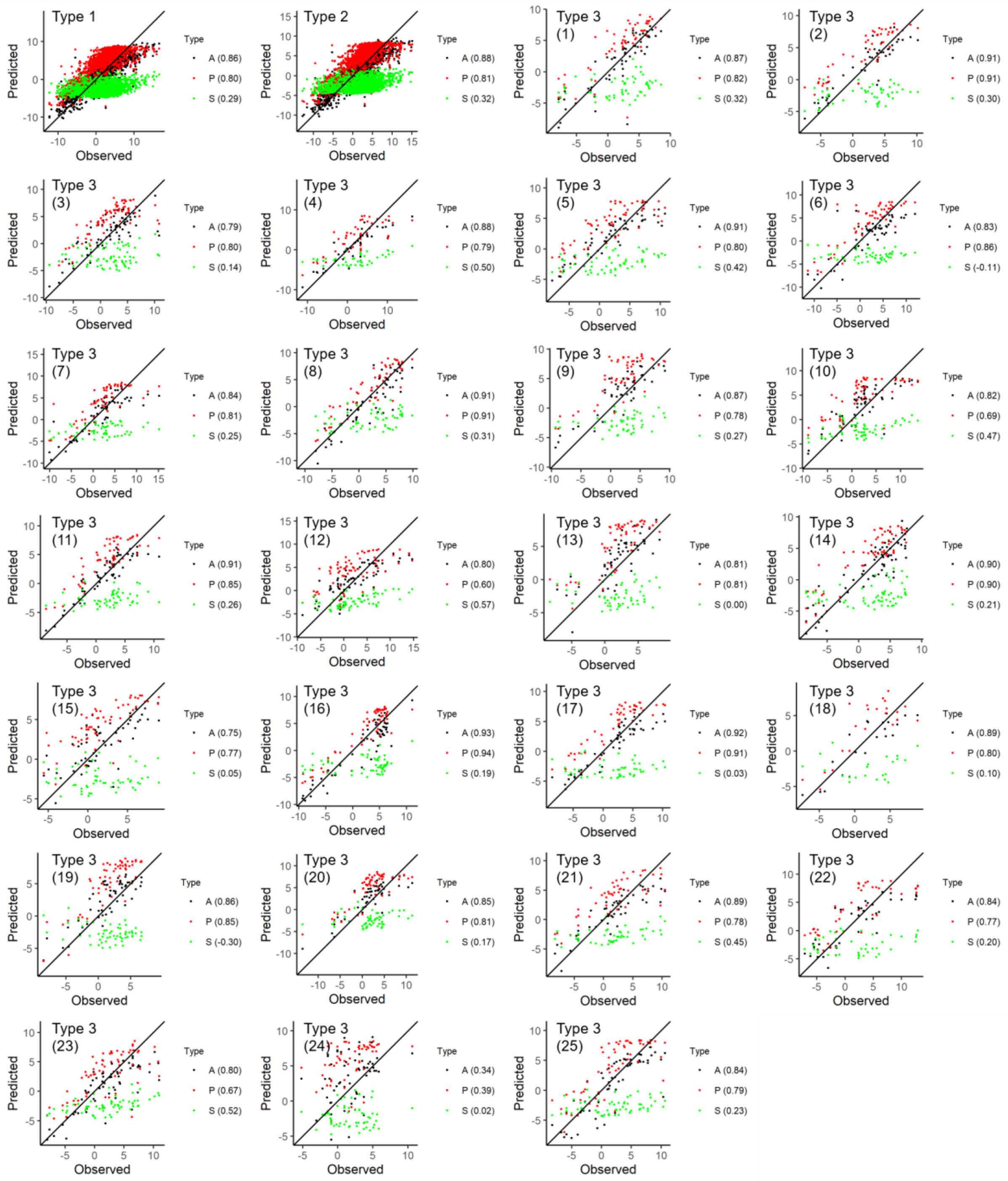


**Supplementary Figure 19. Prediction accuracies on DTH (barley) for analysis type 1-3 using BayesCπ**. Predicted values from all markers (A), favourable primary alleles (P) and favourable secondary alleles (S) are plotted against the observed trait values. Correlations between the predicted and observed values are annotated on each individual plot. Type 1 combines all 25 NAM families, type 2 combines all 25 NAM families while accounting for fixed family effect, and type 3 excludes the testing family in its training set and the testing family is indicated on each individual plot.

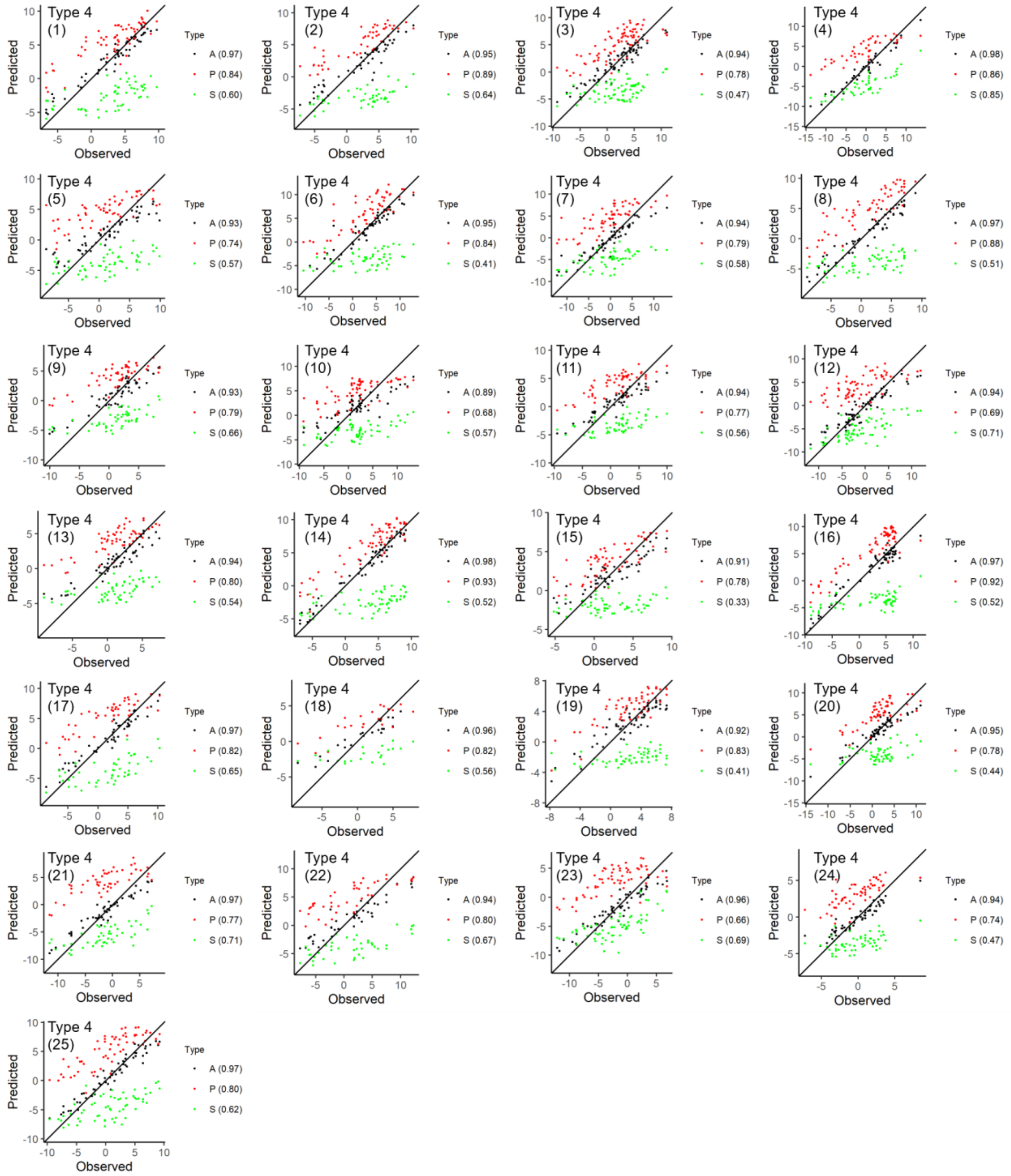


**Supplementary Figure 20. Prediction accuracies on DTH (barley) for analysis type 4 using BayesCπ**. Predicted values from all markers (A), favourable primary alleles (P) and favourable secondary alleles (S) are plotted against the observed trait values. Correlations between the predicted and observed values are annotated on each individual plot. Type 1 combines all 25 NAM families, type 2 combines all 25 NAM families while accounting for fixed family effect, and type 3 excludes the testing family in its training set and the testing family is indicated on each individual plot.

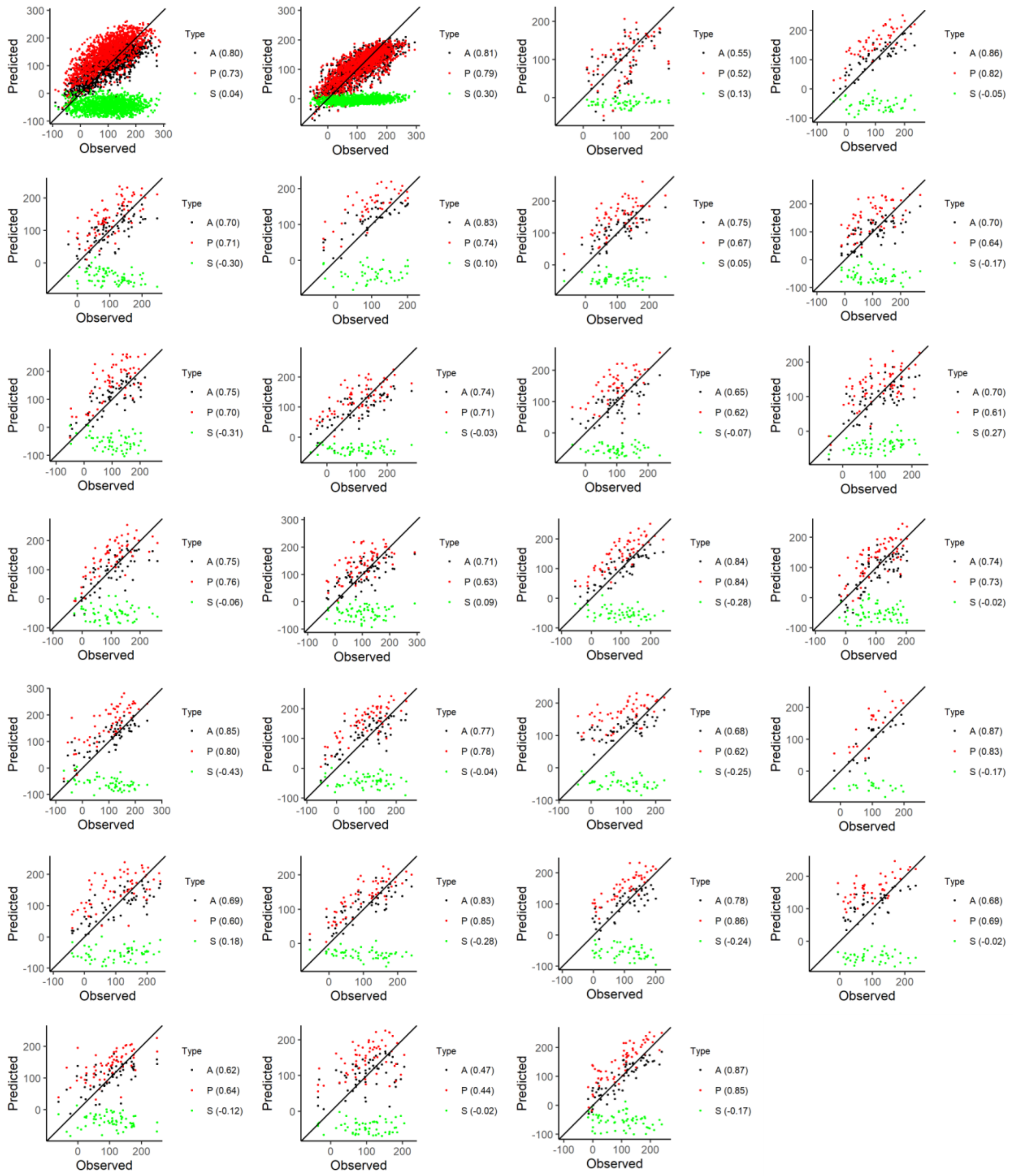


**Supplementary Figure 21. Prediction accuracies on YLD (barley) for analysis type 1-3 using BayesCπ**. Predicted values from all markers (A), favourable primary alleles (P) and favourable secondary alleles (S) are plotted against the observed trait values. Correlations between the predicted and observed values are annotated on each individual plot. Type 1 combines all 25 NAM families, type 2 combines all 25 NAM families while accounting for fixed family effect, and type 3 excludes the testing family in its training set and the testing family is indicated on each individual plot.

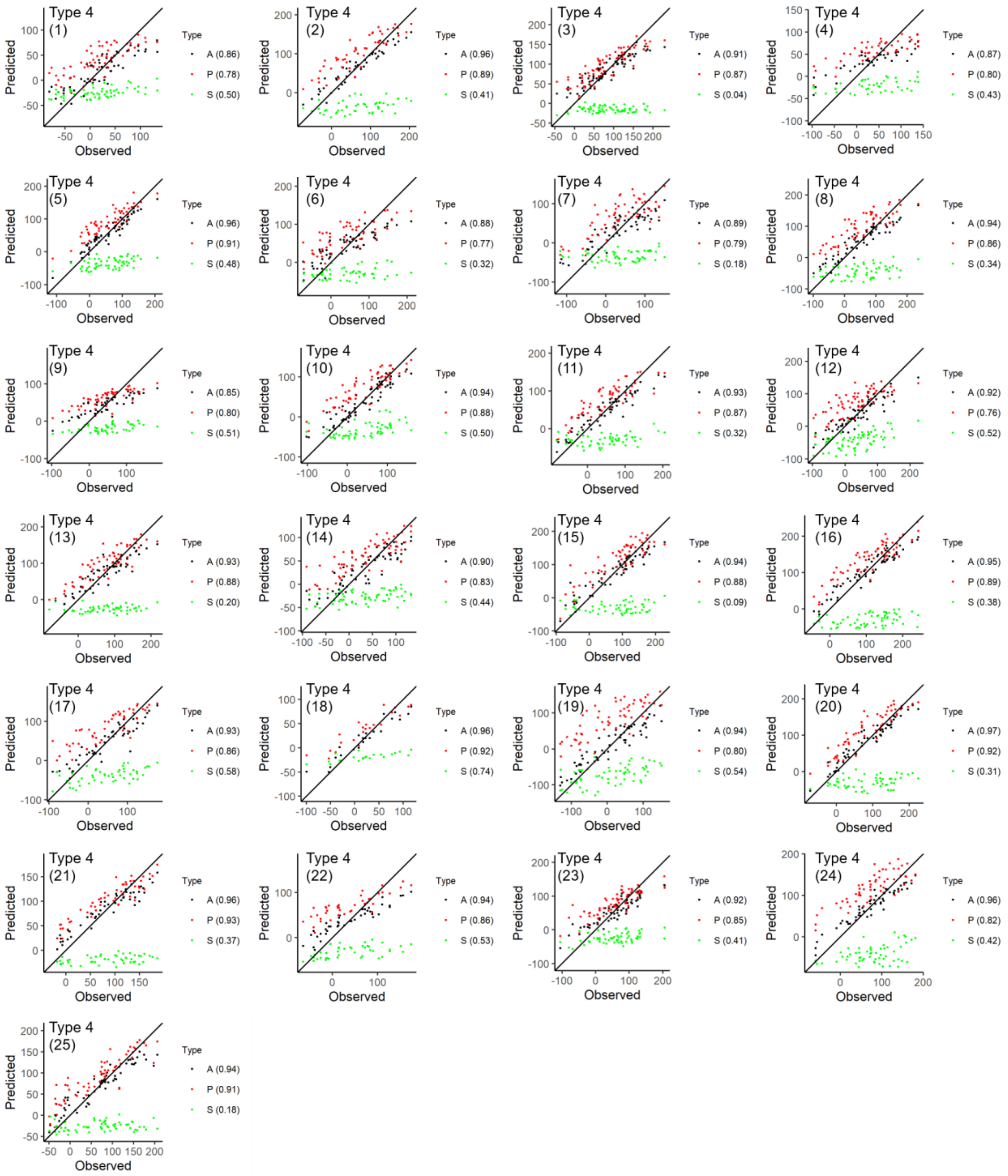


**Supplementary Figure 22. Prediction accuracies on YLD (barley) for analysis type 4 using BayesCπ**. Predicted values from all markers (A), favourable primary alleles (P) and favourable secondary alleles (S) are plotted against the observed trait values. Correlations between the predicted and observed values are annotated on each individual plot. Type 1 combines all 25 NAM families, type 2 combines all 25 NAM families while accounting for fixed family effect, and type 3 excludes the testing family in its training set and the testing family is indicated on each individual plot.

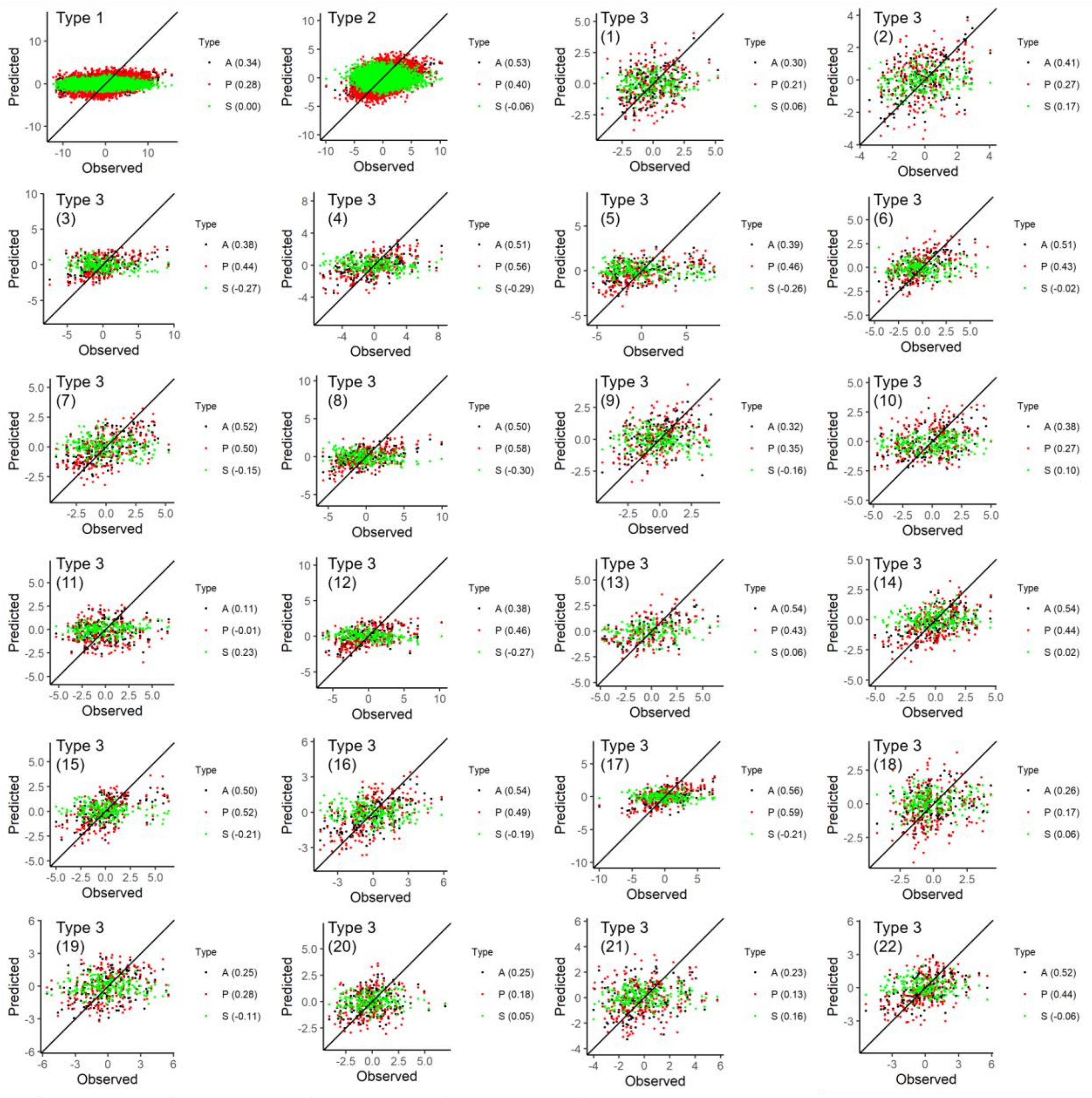


**Supplementary Figure 23. Prediction accuracies on DTS (maize) for analysis type 1-3 using BayesCπ**. Predicted values from all markers (A), favourable primary alleles (P) and favourable secondary alleles (S) are plotted against the observed trait values. Correlations between the predicted and observed values are annotated on each individual plot. Type 1 combines all 25 NAM families, type 2 combines all 25 NAM families while accounting for fixed family effect, and type 3 excludes the testing family in its training set and the testing family is indicated on each individual plot.

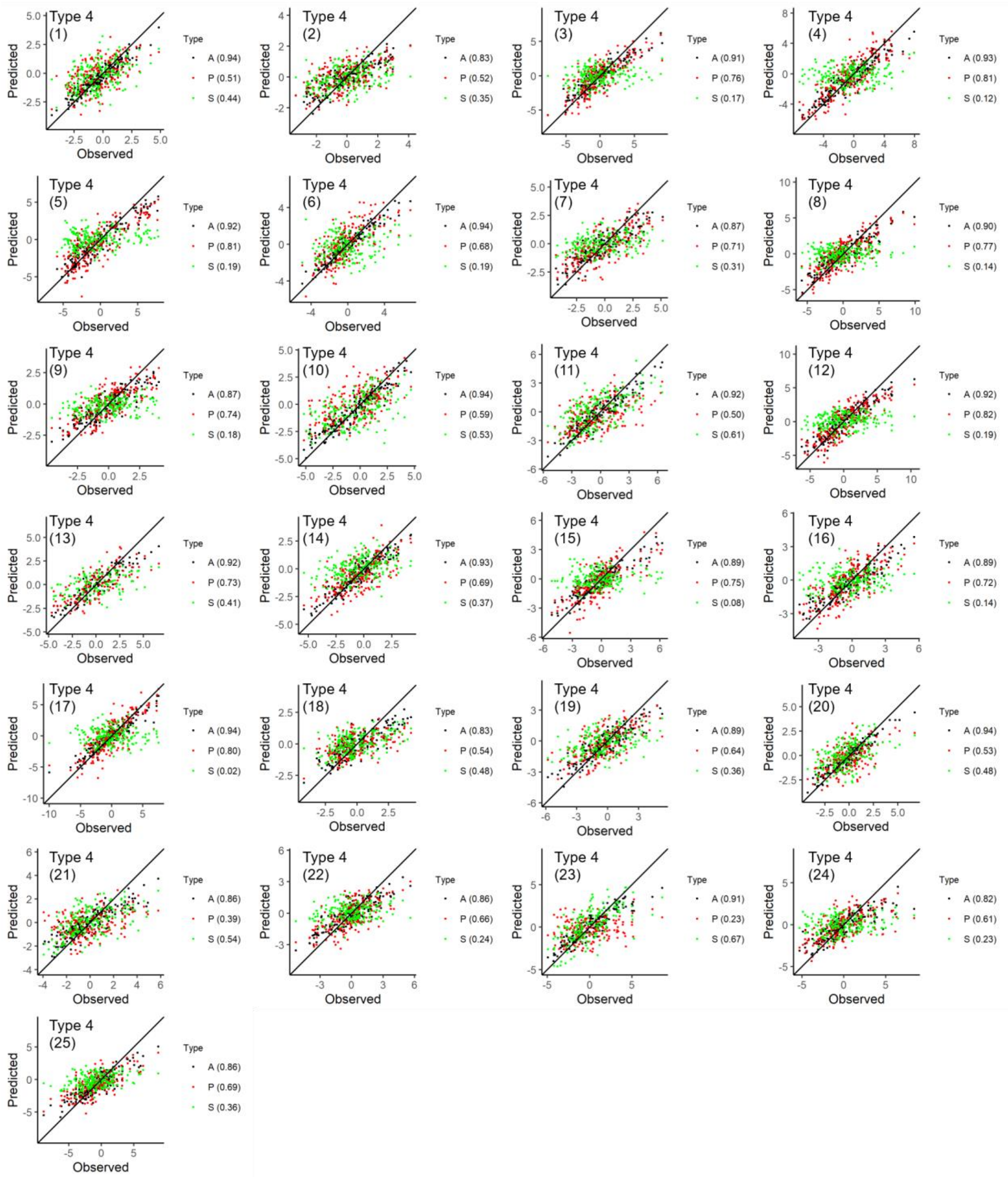


**Supplementary Figure 24. Prediction accuracies on DTS (maize) for analysis type 4 using BayesCπ**. Predicted values from all markers (A), favourable primary alleles (P) and favourable secondary alleles (S) are plotted against the observed trait values. Correlations between the predicted and observed values are annotated on each individual plot. Type 1 combines all 25 NAM families, type 2 combines all 25 NAM families while accounting for fixed family effect, and type 3 excludes the testing family in its training set and the testing family is indicated on each individual plot.

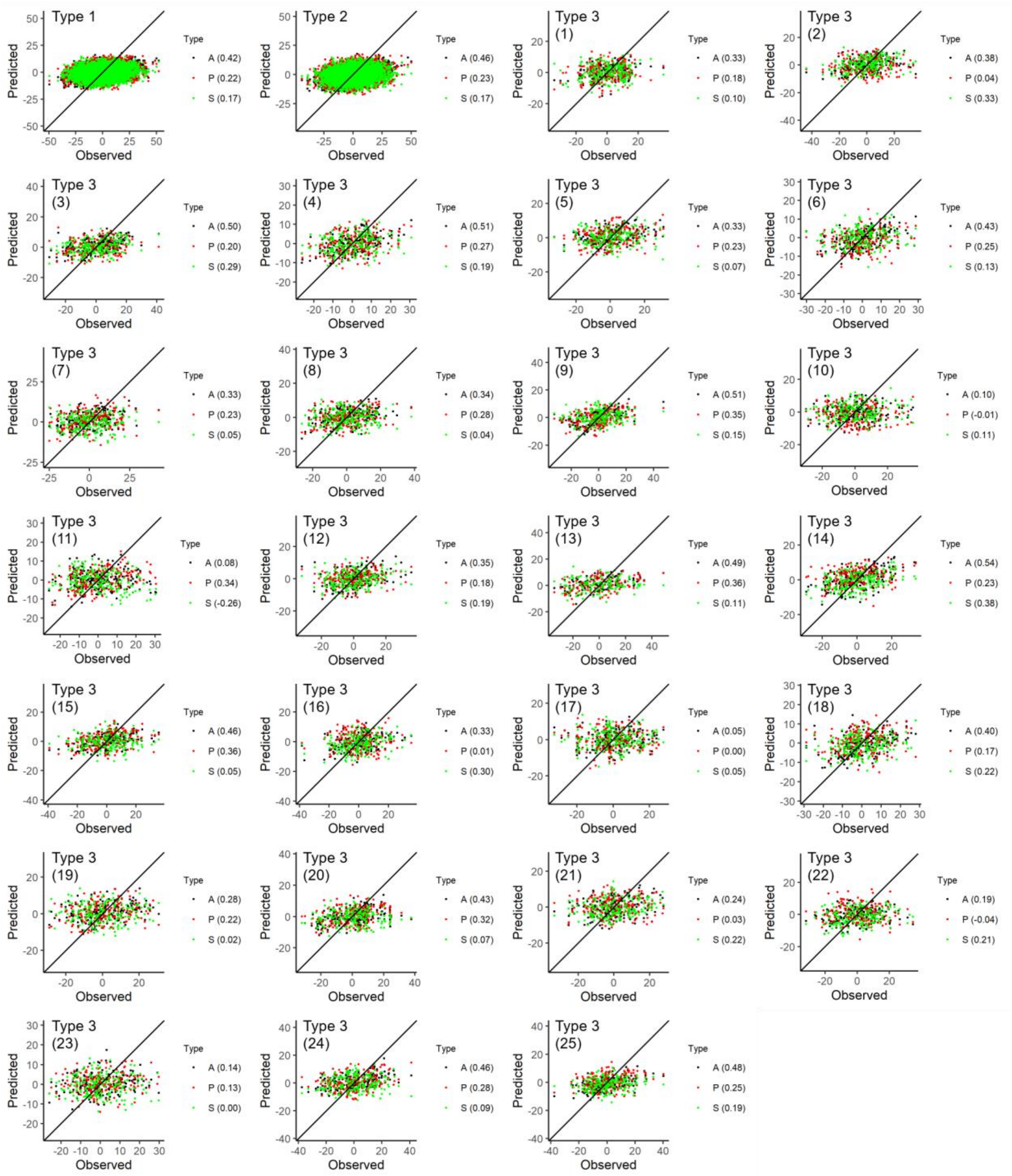


**Supplementary Figure 25. Prediction accuracies on CL (maize) for analysis type 1-3 using BayesCπ**. Predicted values from all markers (A), favourable primary alleles (P) and favourable secondary alleles (S) are plotted against the observed trait values. Correlations between the predicted and observed values are annotated on each individual plot. Type 1 combines all 25 NAM families, type 2 combines all 25 NAM families while accounting for fixed family effect, and type 3 excludes the testing family in its training set and the testing family is indicated on each individual plot.

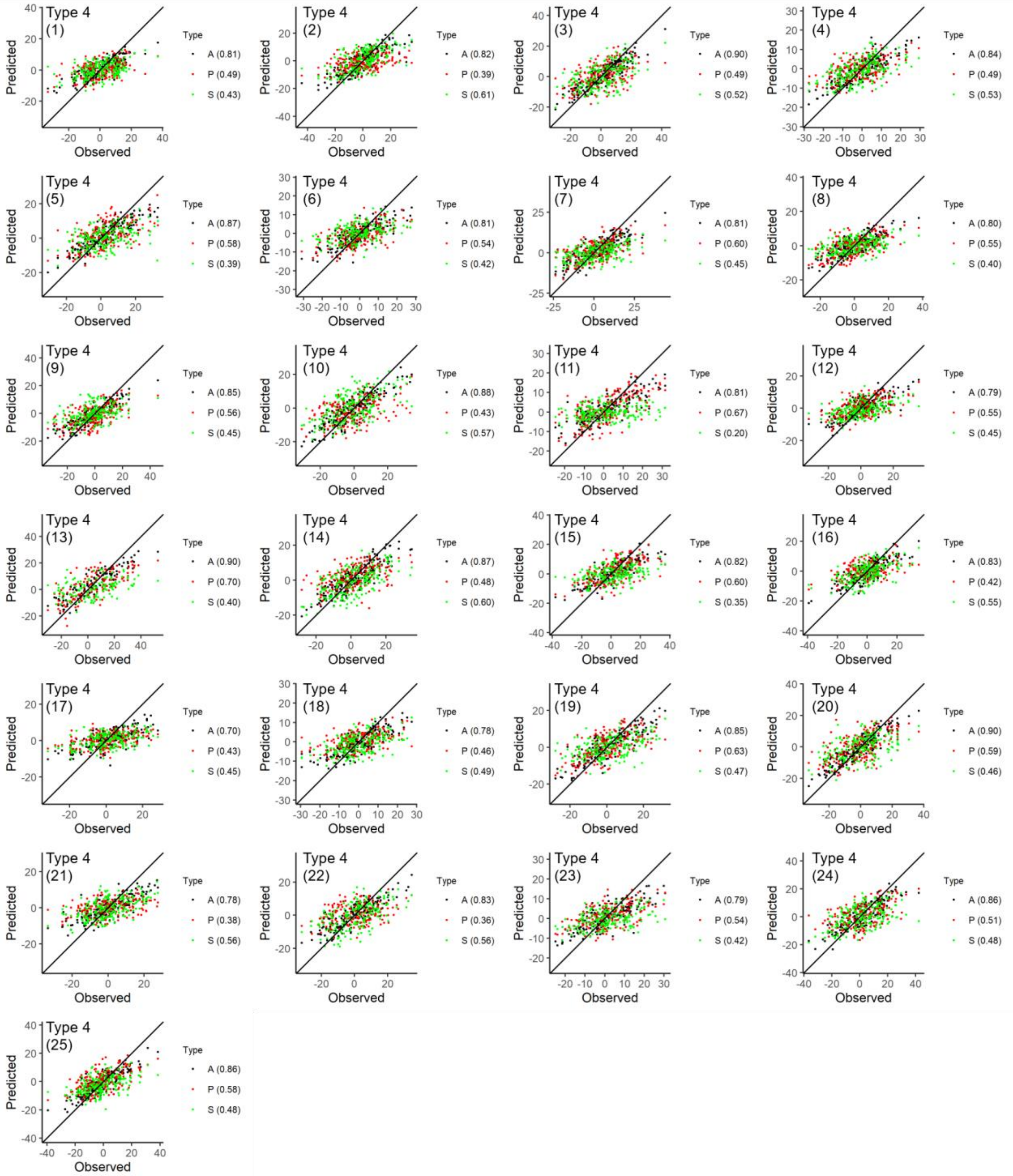


**Supplementary Figure 26. Prediction accuracies on CL (maize) for analysis type 4 using BayesCπ**. Predicted values from all markers (A), favourable primary alleles (P) and favourable secondary alleles (S) are plotted against the observed trait values. Correlations between the predicted and observed values are annotated on each individual plot. Type 1 combines all 25 NAM families, type 2 combines all 25 NAM families while accounting for fixed family effect, and type 3 excludes the testing family in its training set and the testing family is indicated on each individual plot.

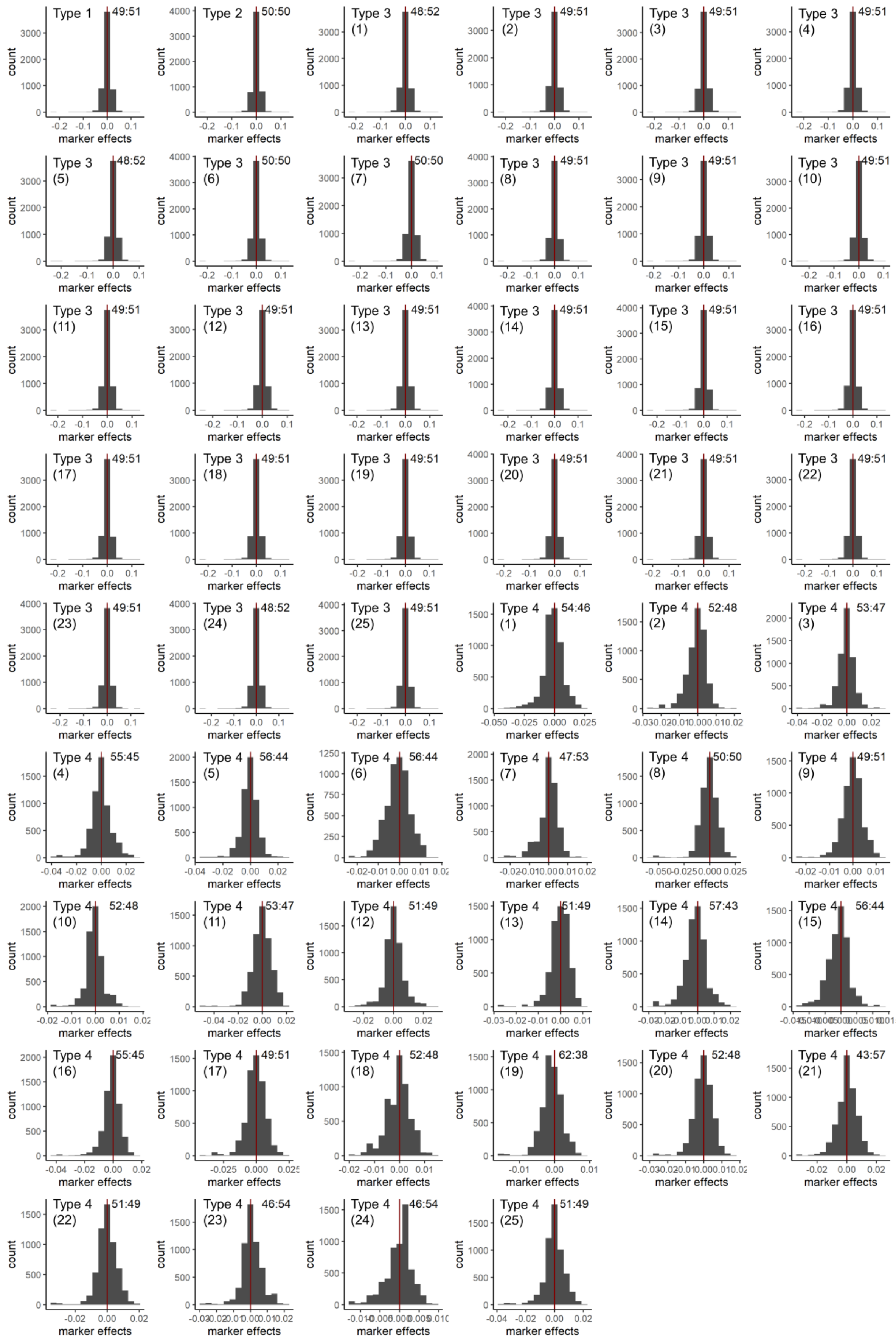


**Supplementary Figure 27. Distributions of non-zero DTH (barley) marker effects for analysis type 1-4 using rrBLUP**.

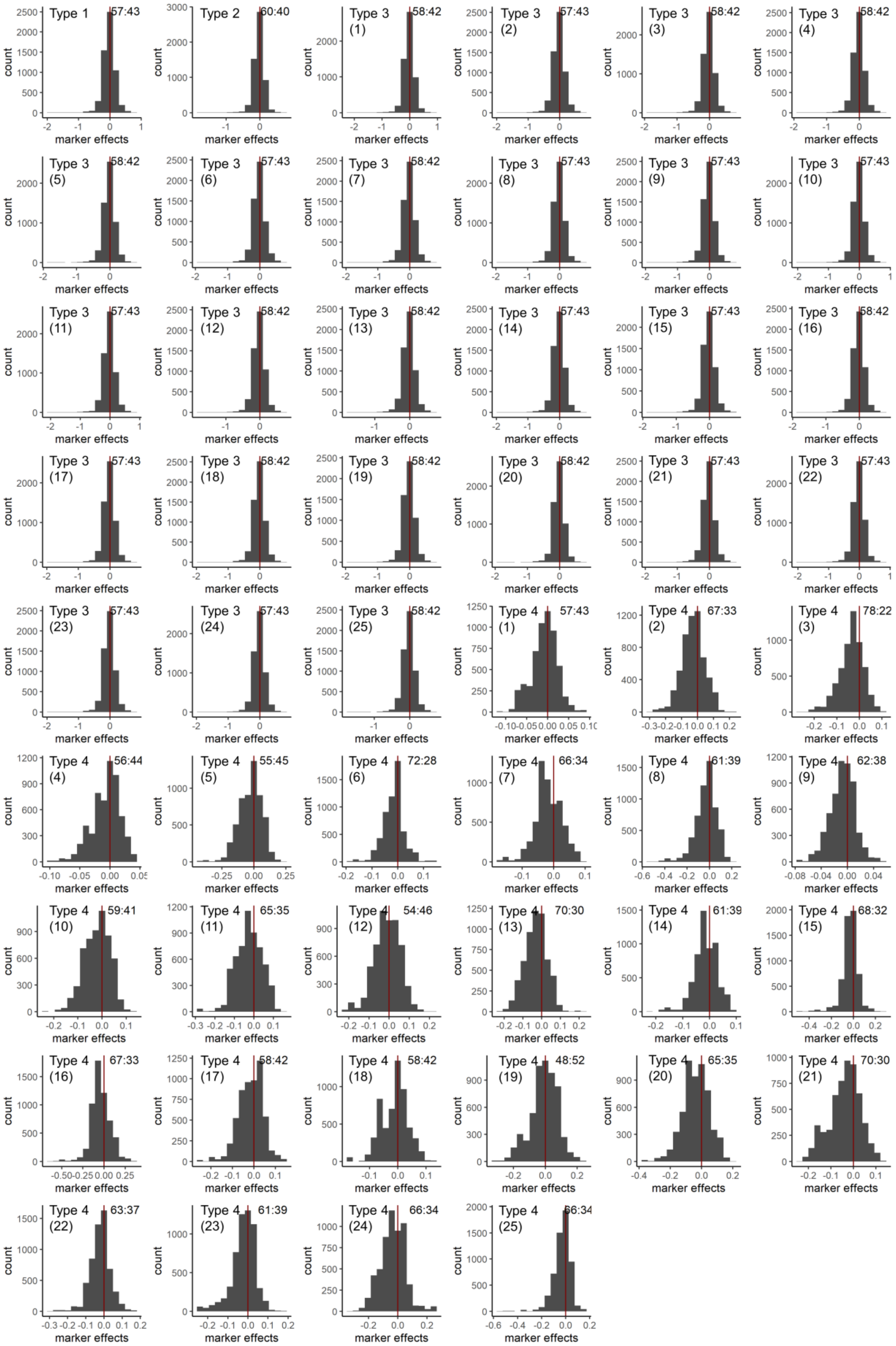


**Supplementary Figure 28. Distributions of non-zero YLD (barley) marker effects for analysis type 1-4 using rrBLUP**.

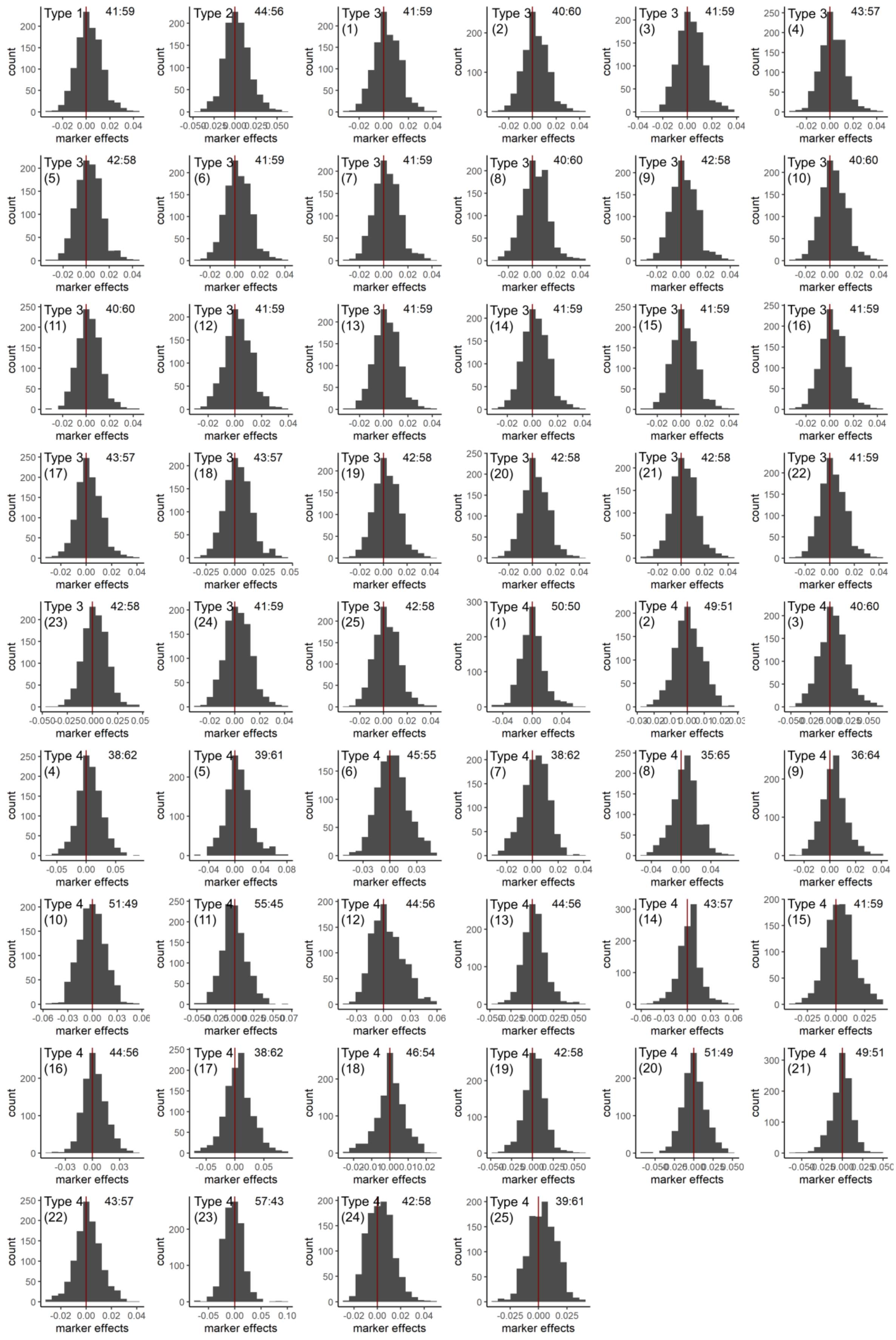

**Supplementary Figure 29. Distributions of non-zero DTS (maize) marker effects for analysis type 1-4 using rrBLUP**.

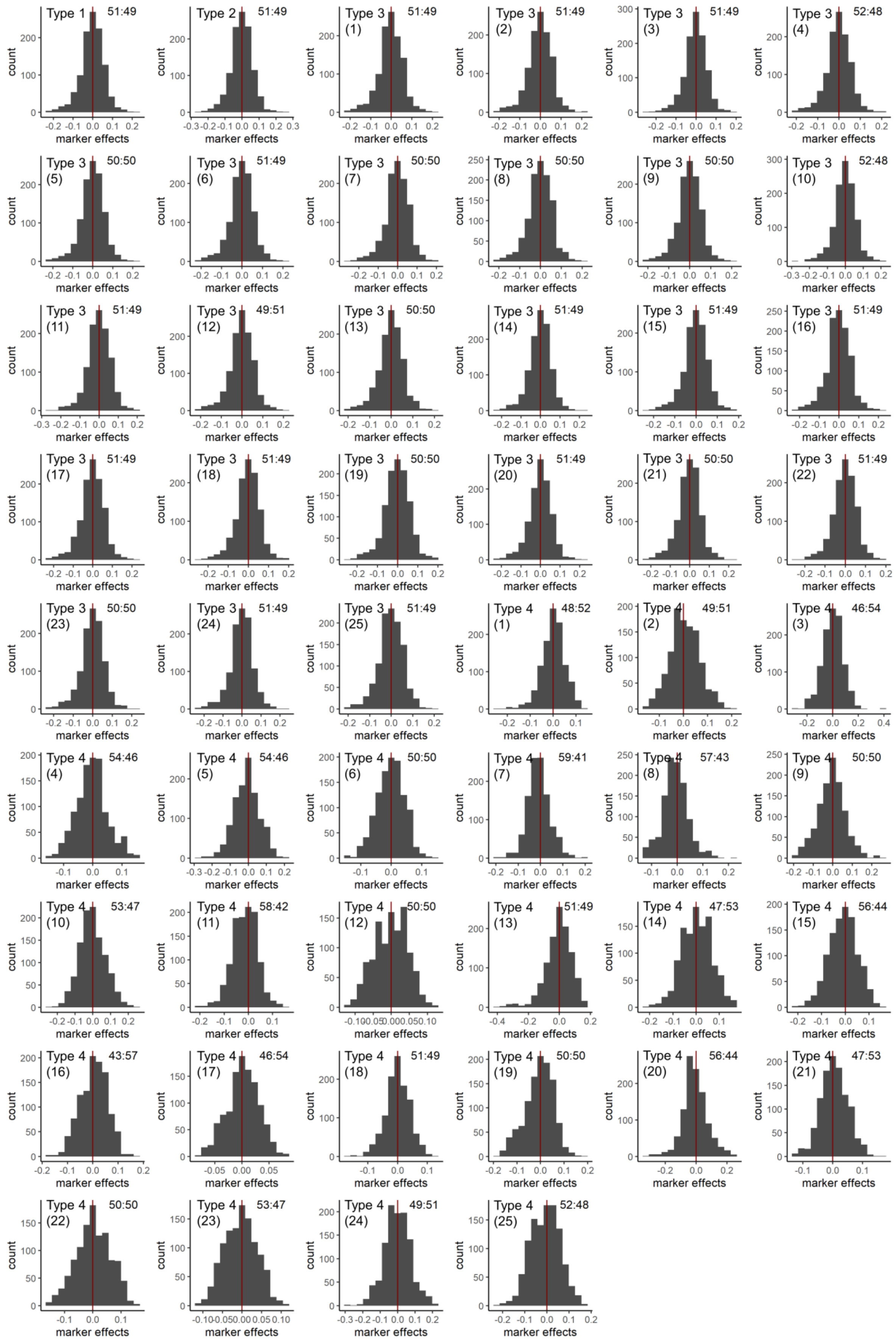

**Supplementary Figure 30. Distributions of non-zero CL (maize) marker effects for analysis type 1-4 using rrBLUP**.

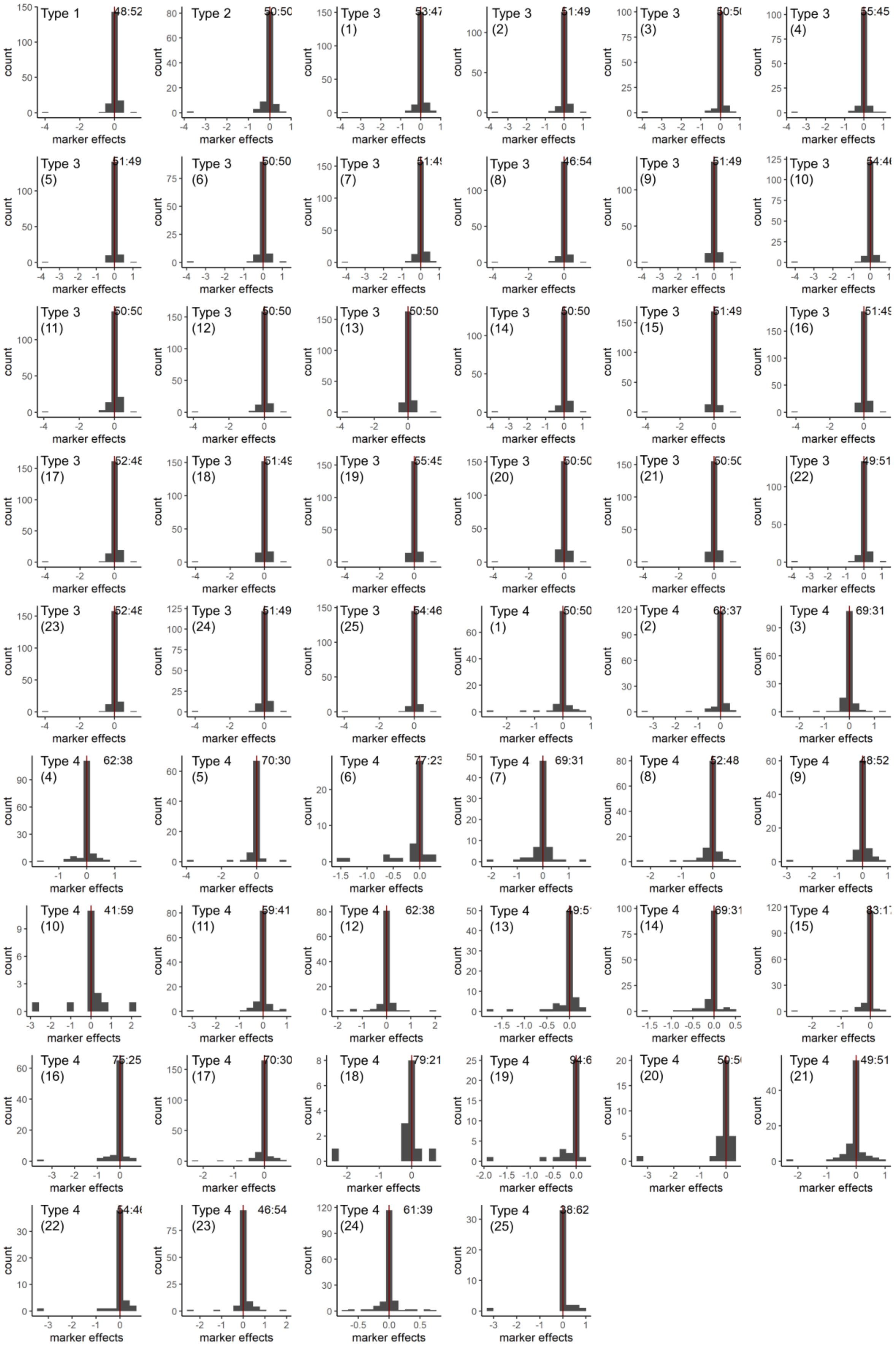


**Supplementary Figure 31. Distributions of non-zero DTH (barley) marker effects for analysis type 1-4 using LASSO**.

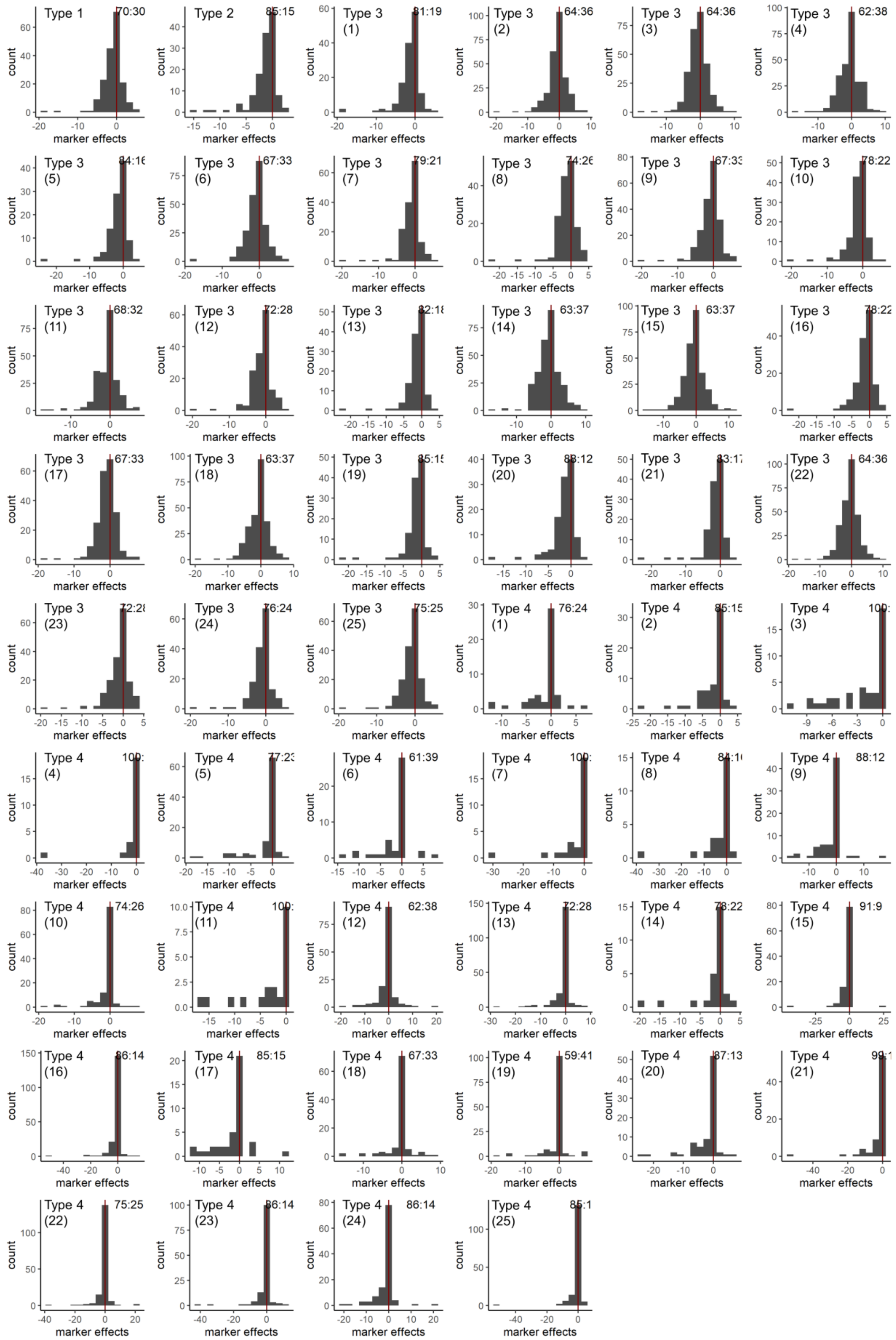

**Supplementary Figure 32. Distributions of non-zero YLD (barley) marker effects for analysis type 1-4 using LASSO**.

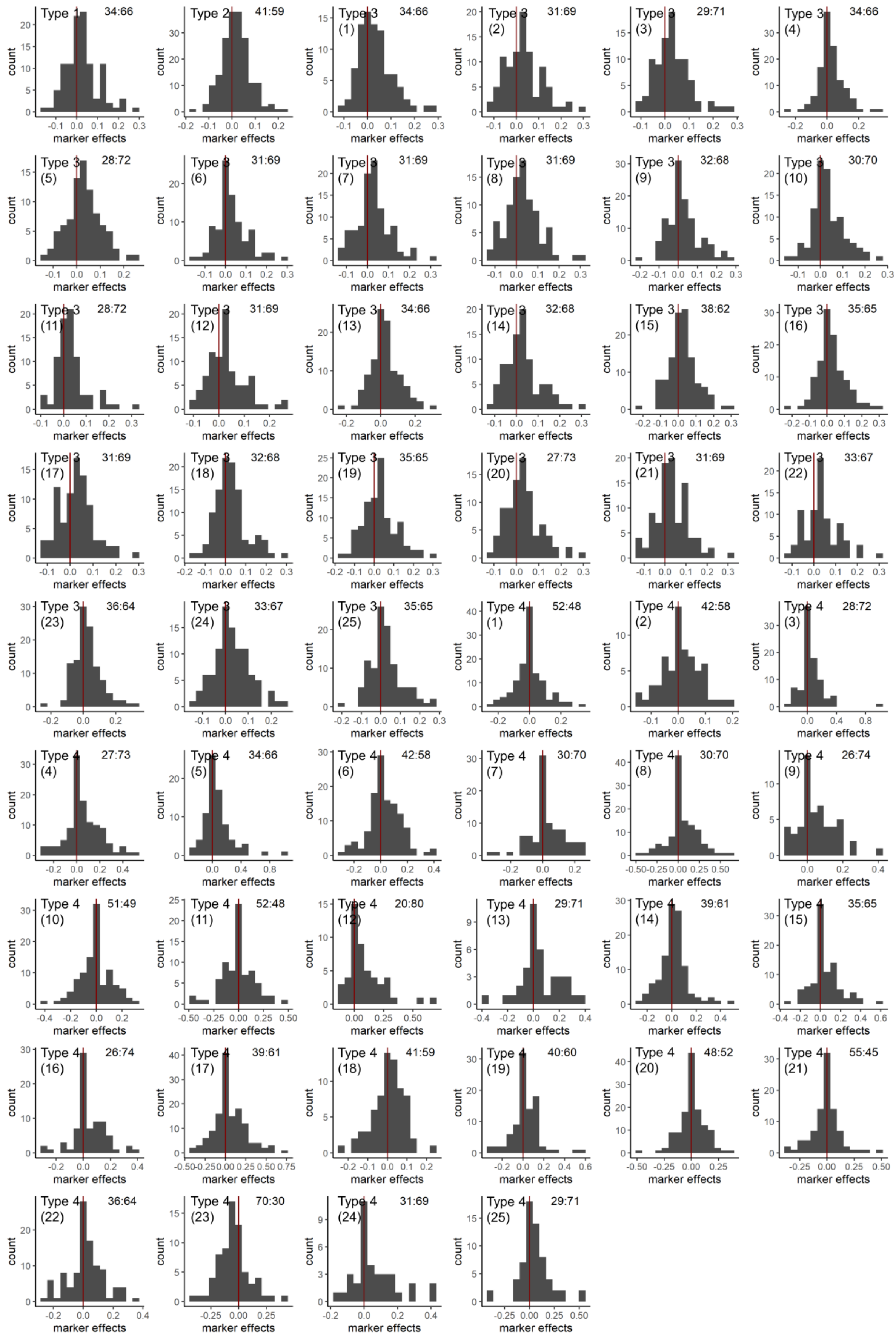


**Supplementary Figure 33. Distributions of non-zero DTS (maize) marker effects for analysis type 1-4 using LASSO**.

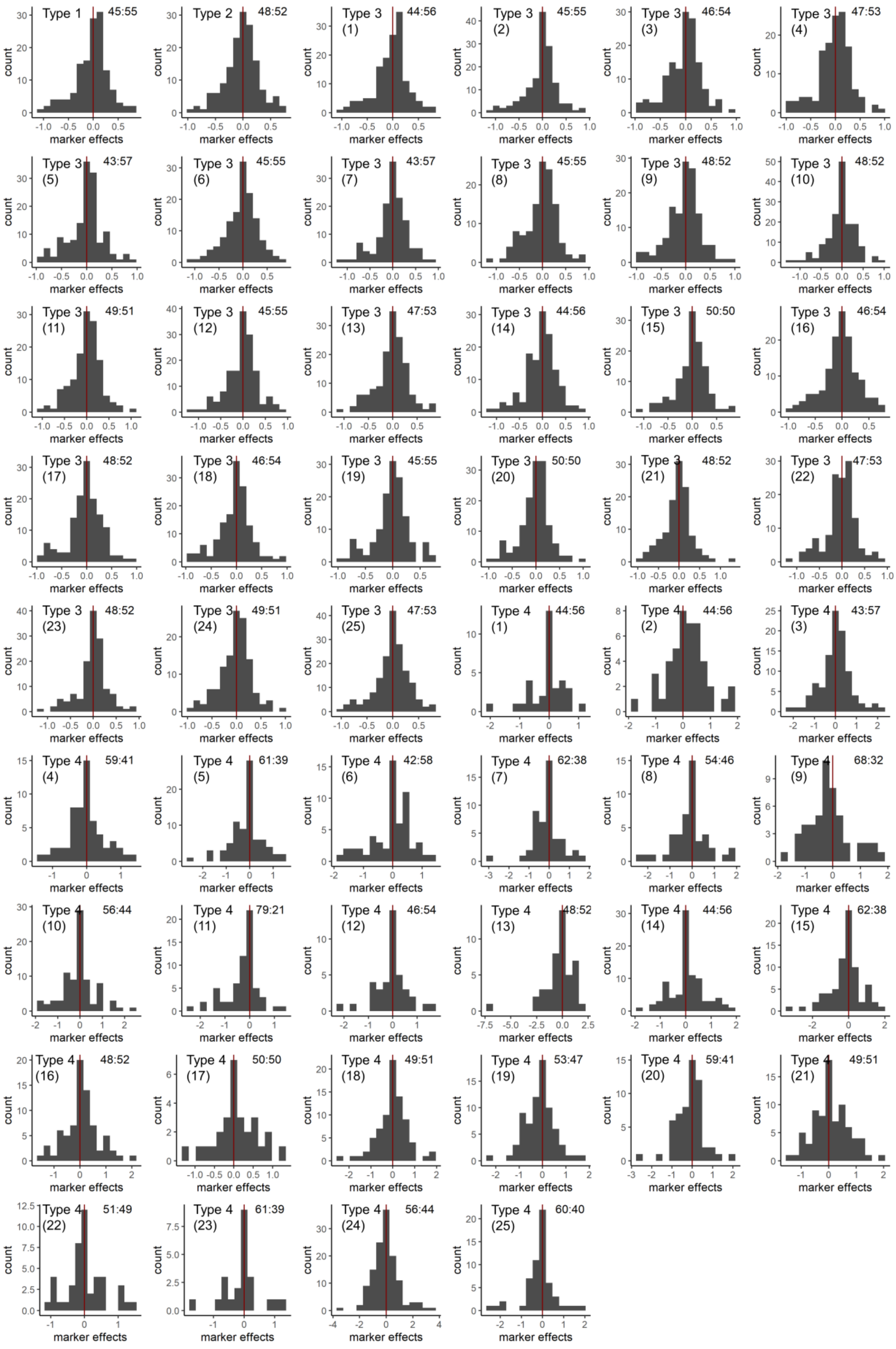


**Supplementary Figure 34. Distributions of non-zero CL (maize) marker effects for analysis type 1-4 using LASSO**.

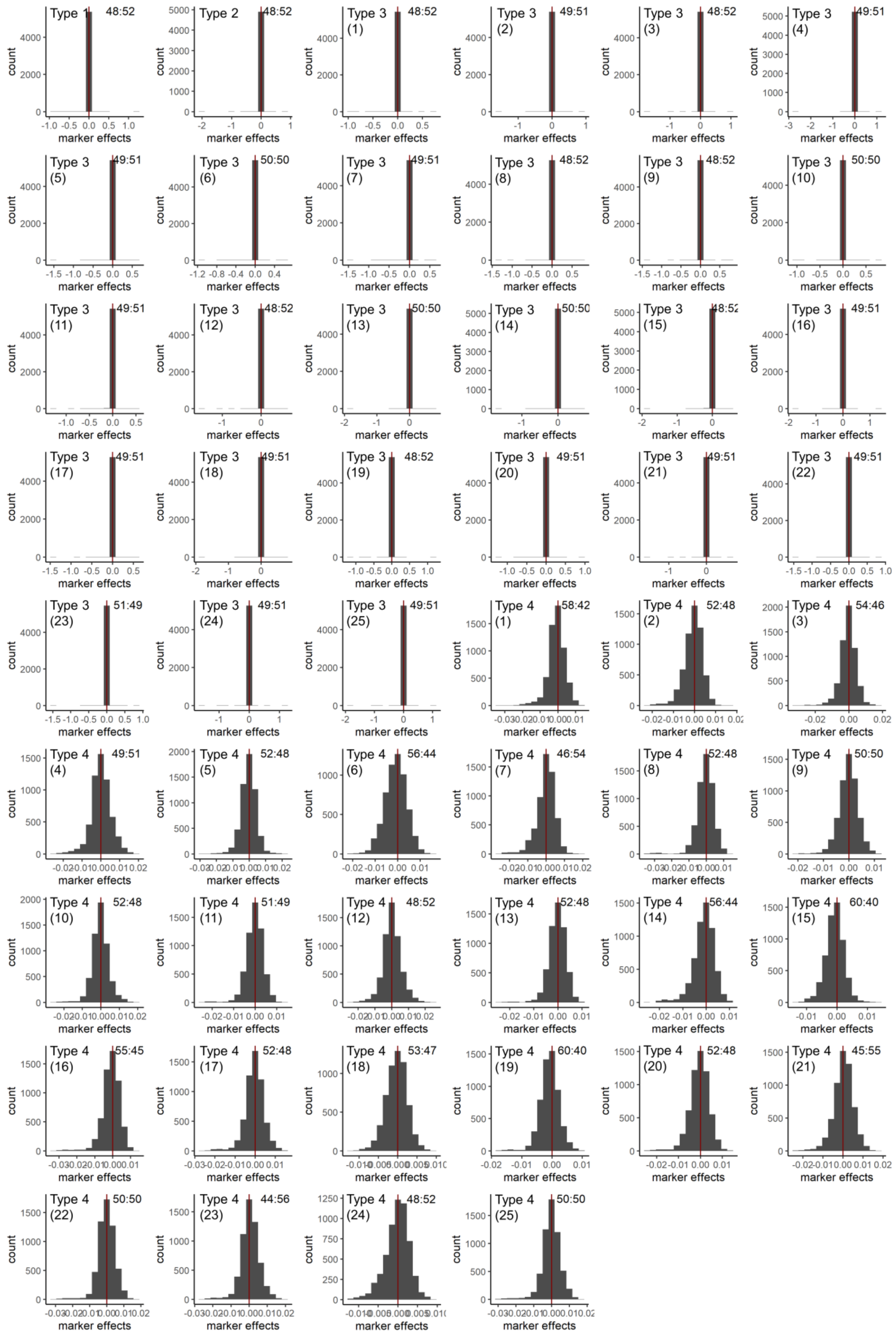


**Supplementary Figure 35. Distributions of non-zero DTH (barley) marker effects for analysis type 1-4 using BayesCπ.**

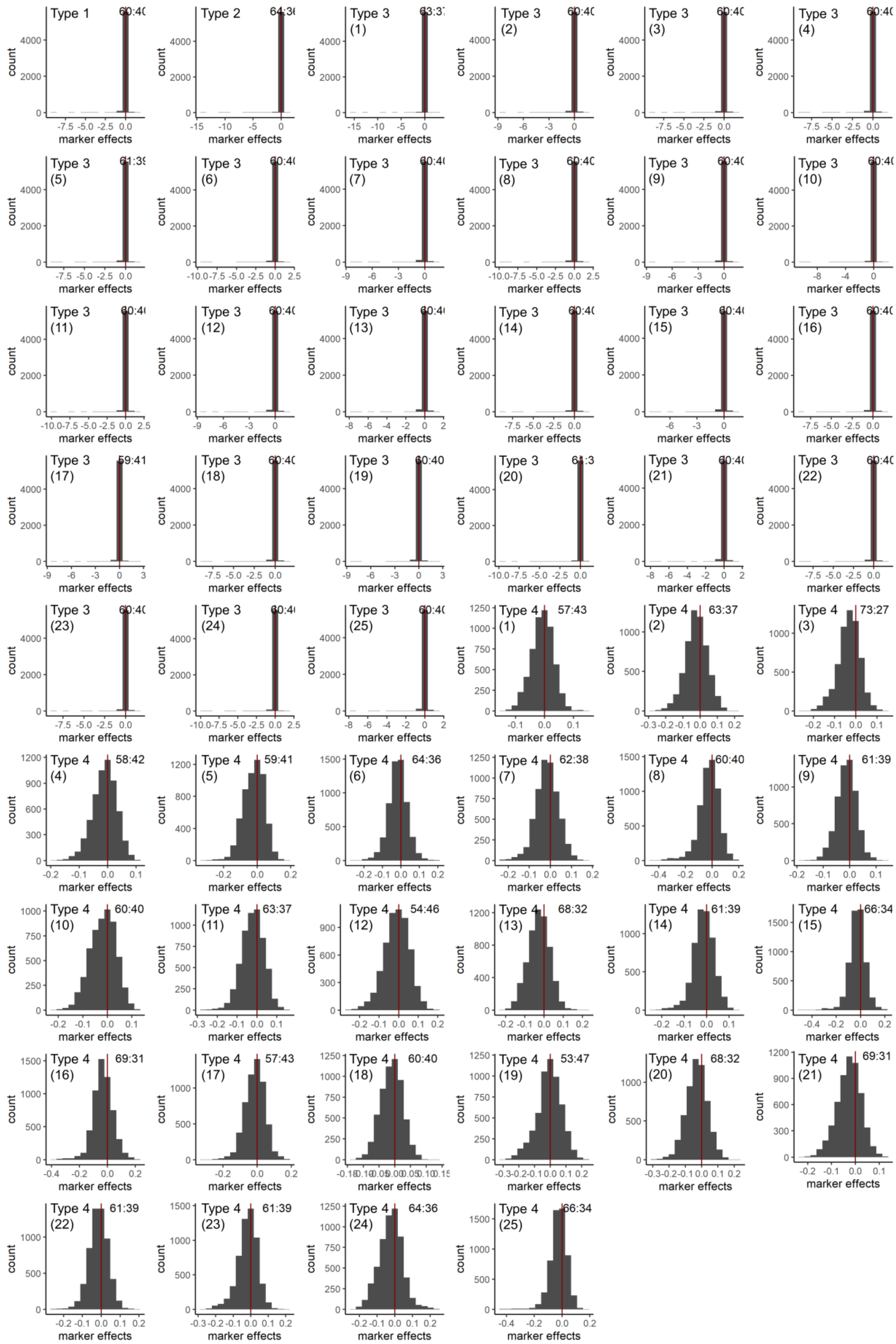

**Supplementary Figure 36. Distributions of non-zero YLD (barley) marker effects for analysis type 1-4 using BayesCπ.**

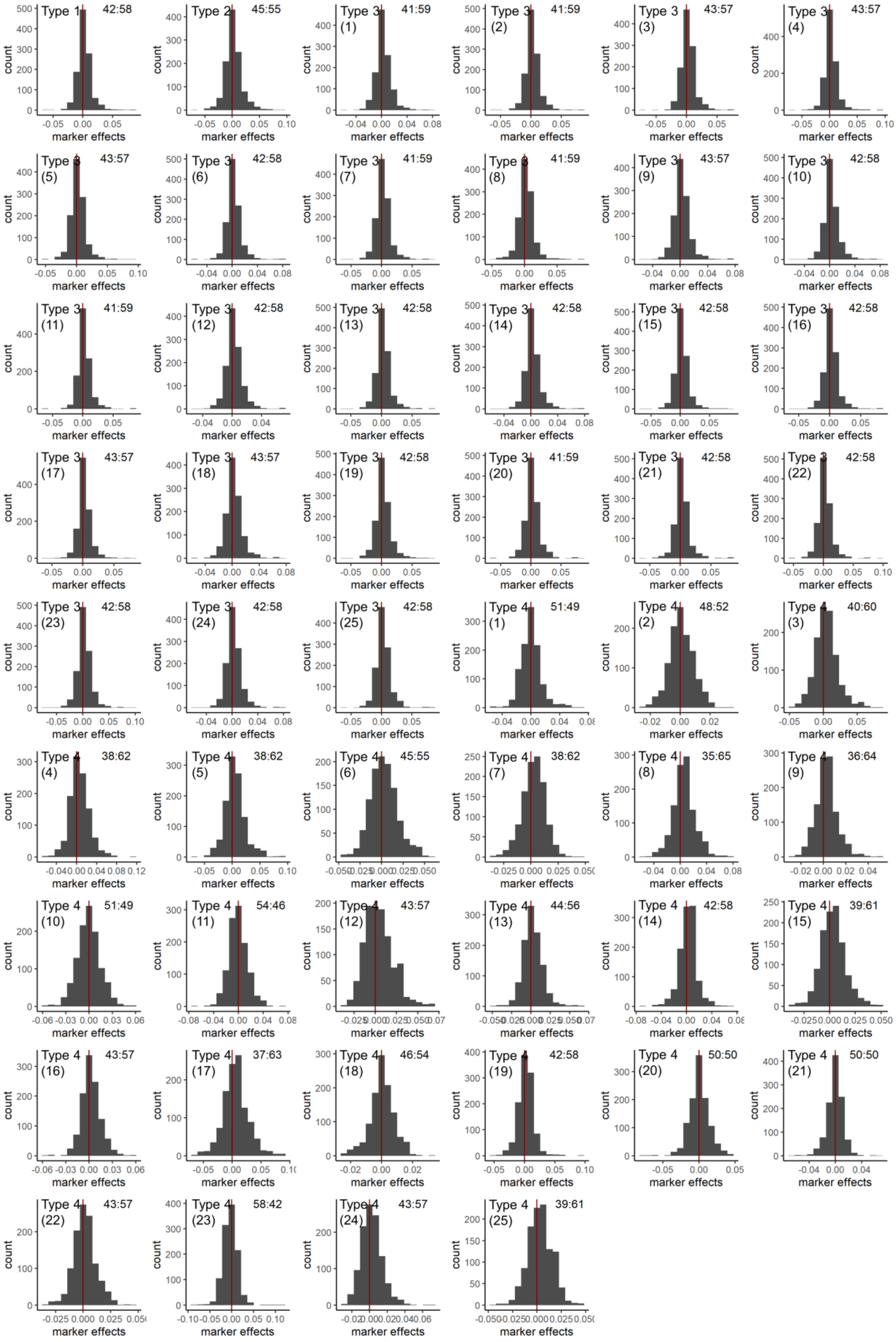


**Supplementary Figure 37. Distributions of non-zero DTS (maize) marker effects for analysis type 1-4 using BayesCπ**.

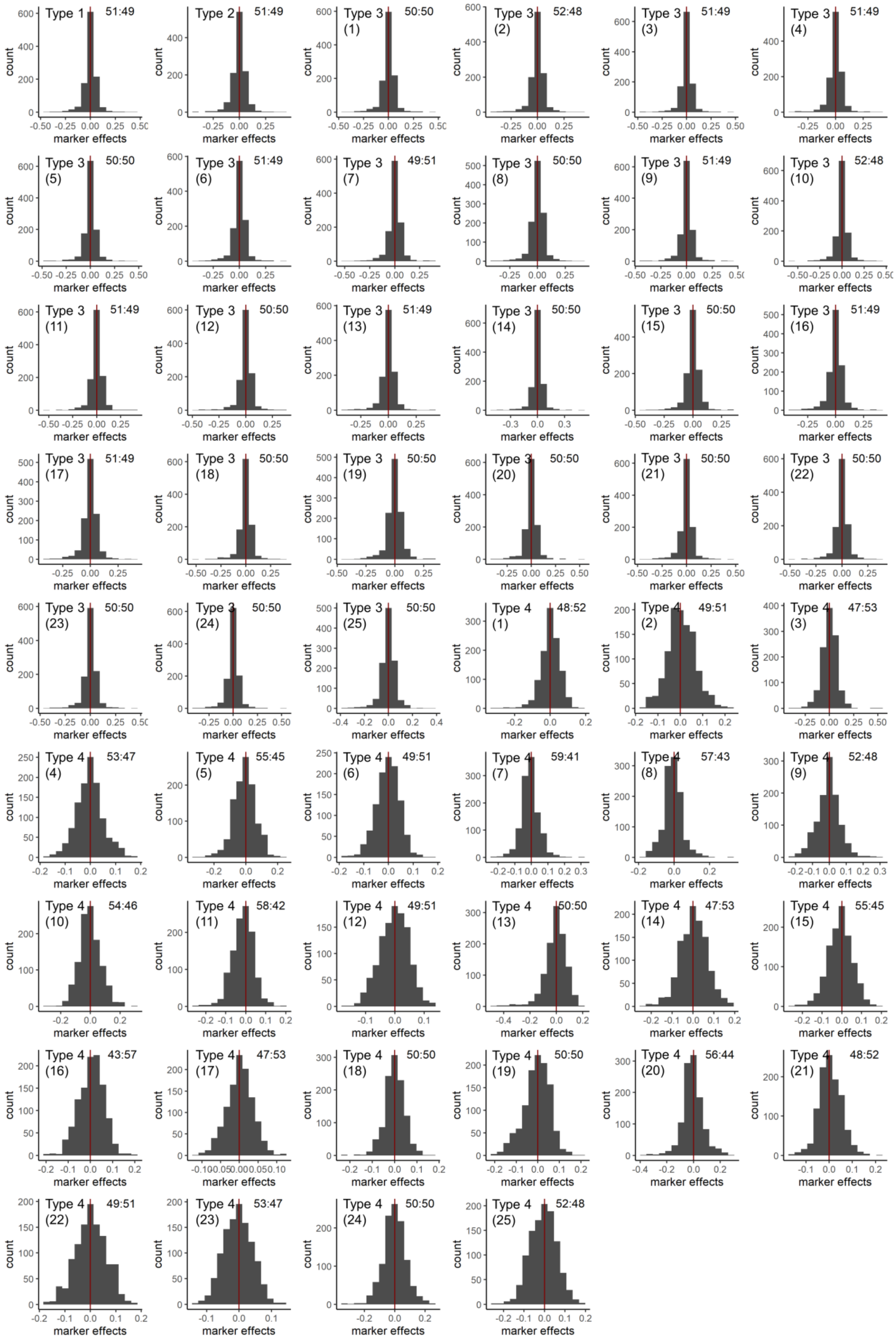


**Supplementary Figure 38. Distributions of non-zero CL (maize) marker effects for analysis type 1-4 using BayesCπ**.

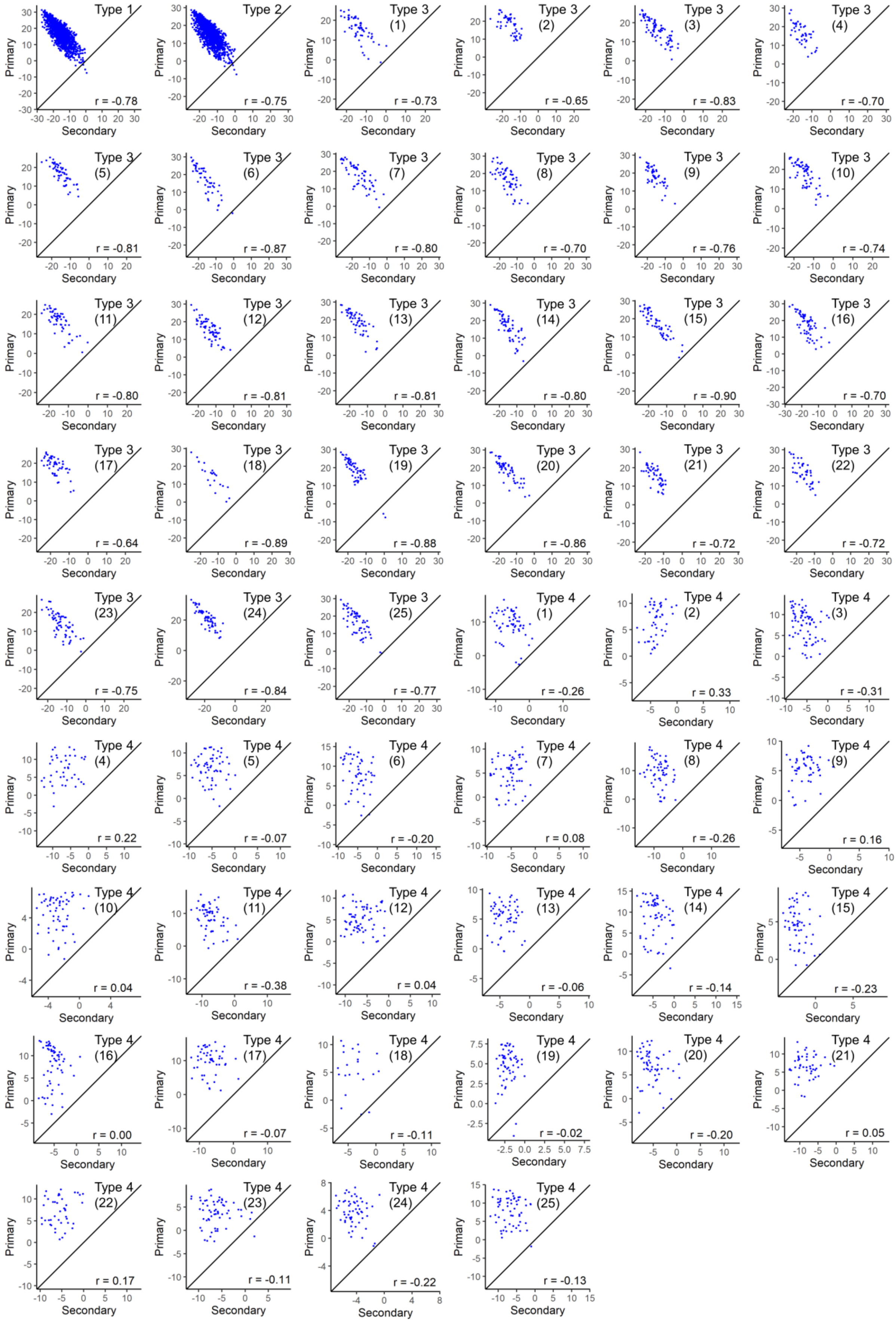


**Supplementary Figure 39. Predictions from favourable primary (P) vs secondary (S) markers of DTH (barley) for analysis type 1-4 using rrBLUP.**

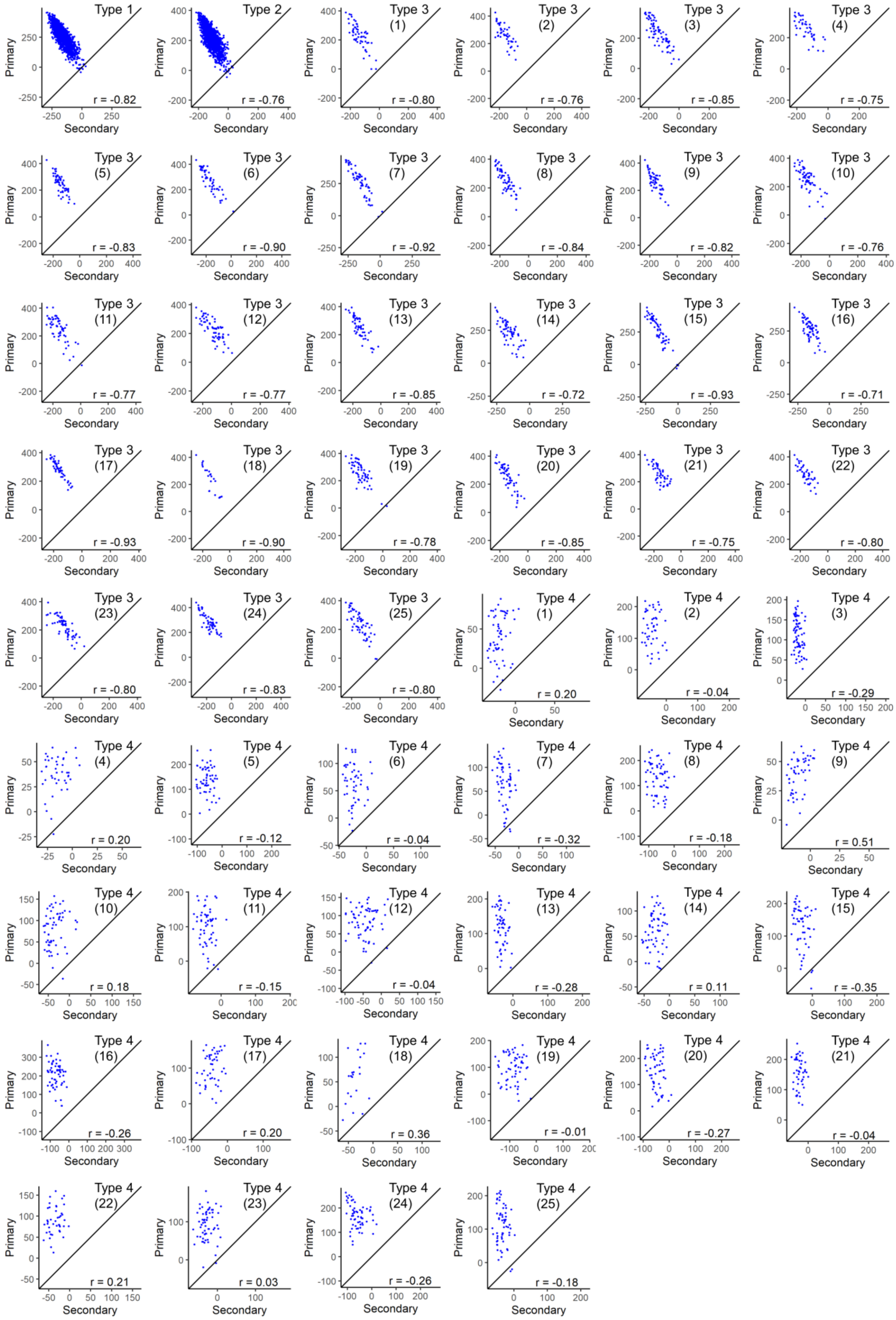

**Supplementary Figure 40. Predictions from favourable primary (P) vs secondary (S) markers of YLD (barley) for analysis type 1-4 using rrBLUP**.

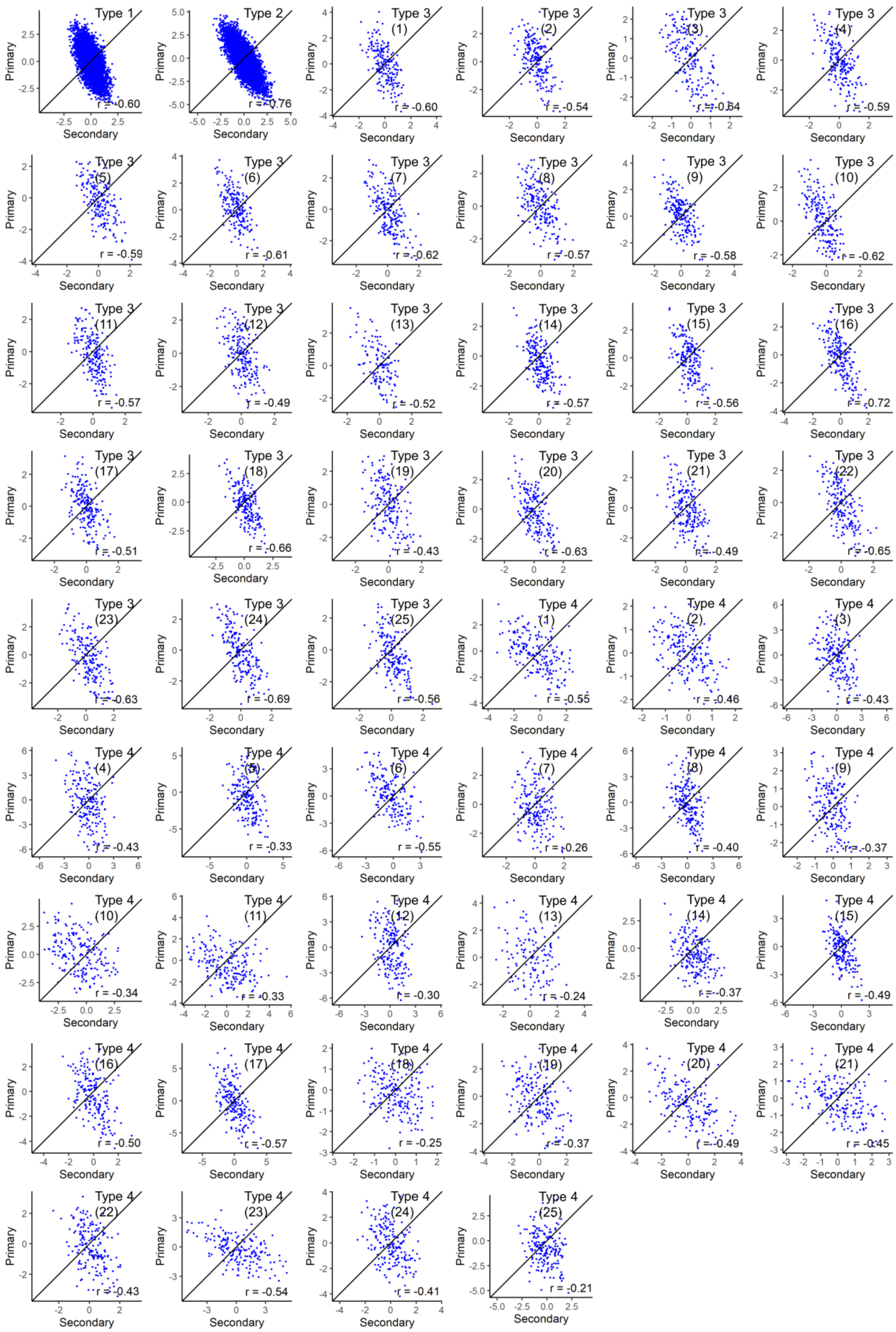

**Supplementary Figure 41. Predictions from favourable primary (P) vs secondary (S) markers of DTS (maize) for analysis type 1-4 using rrBLUP.**

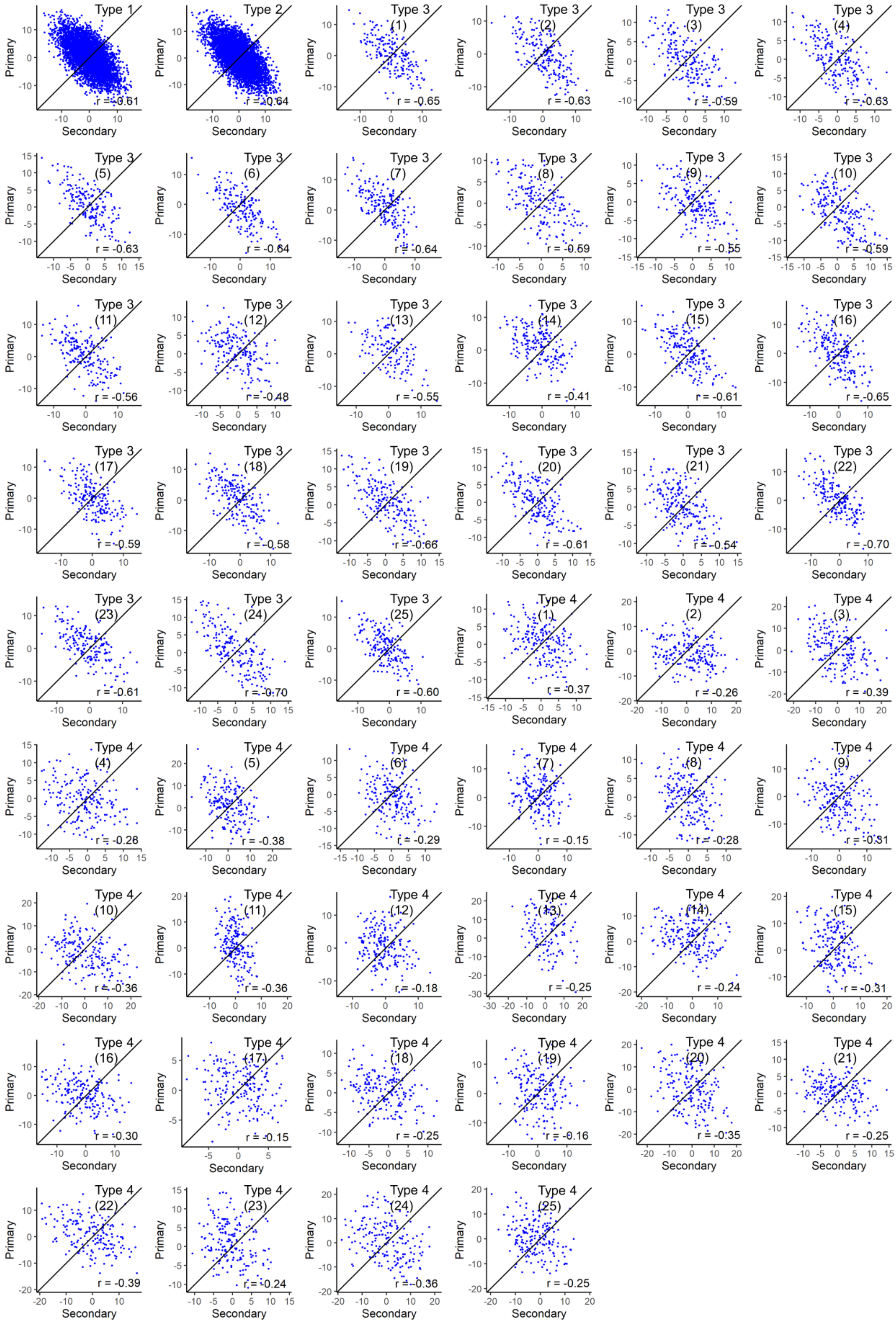


**Supplementary Figure 42. Predictions from favourable primary (P) vs secondary (S) markers of CL (maize) for analysis type 1-4 using rrBLUP**.

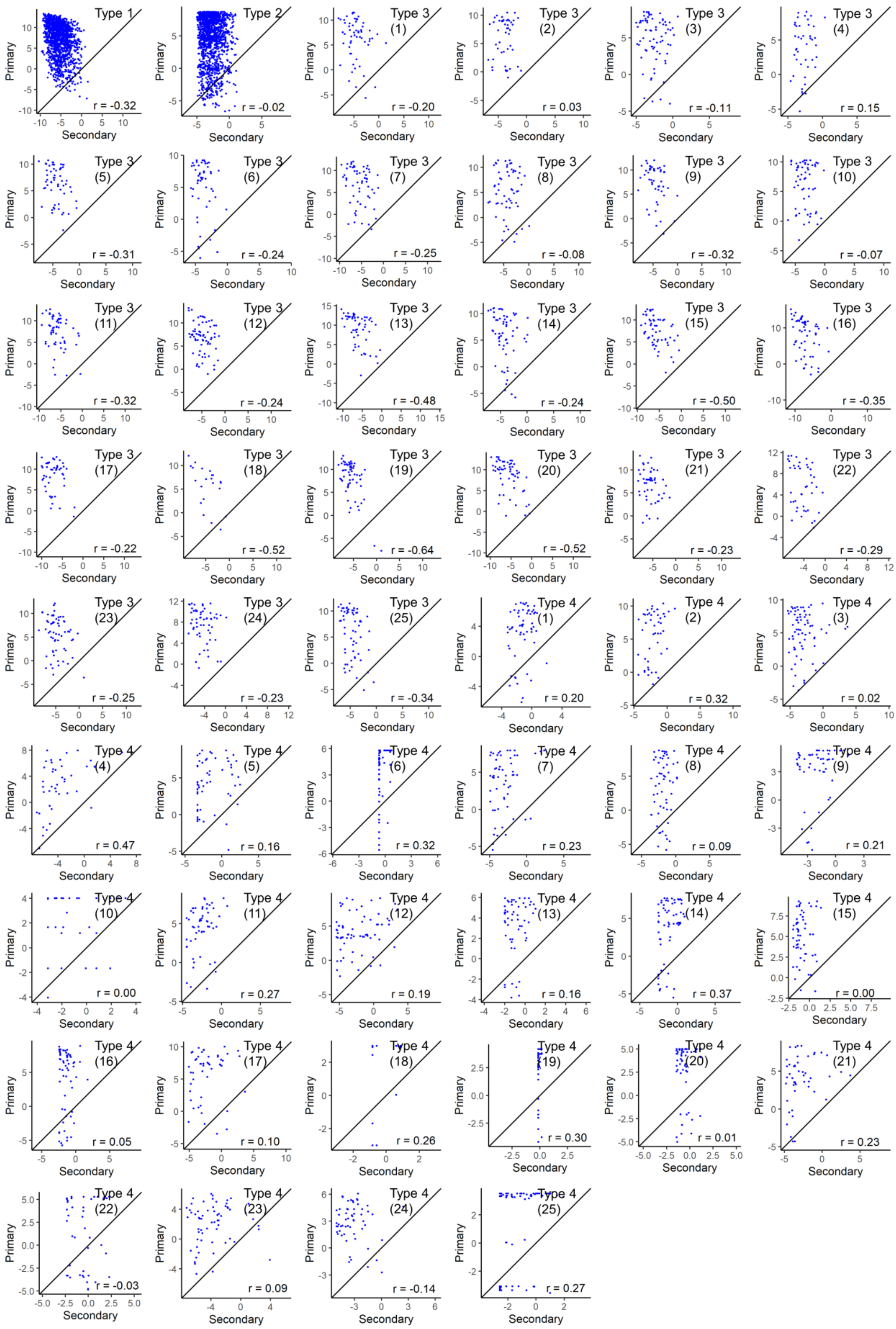


**Supplementary Figure 43. Predictions from favourable primary (P) vs secondary (S) markers of DTH (barley) for analysis type 1-4 using LASSO.**

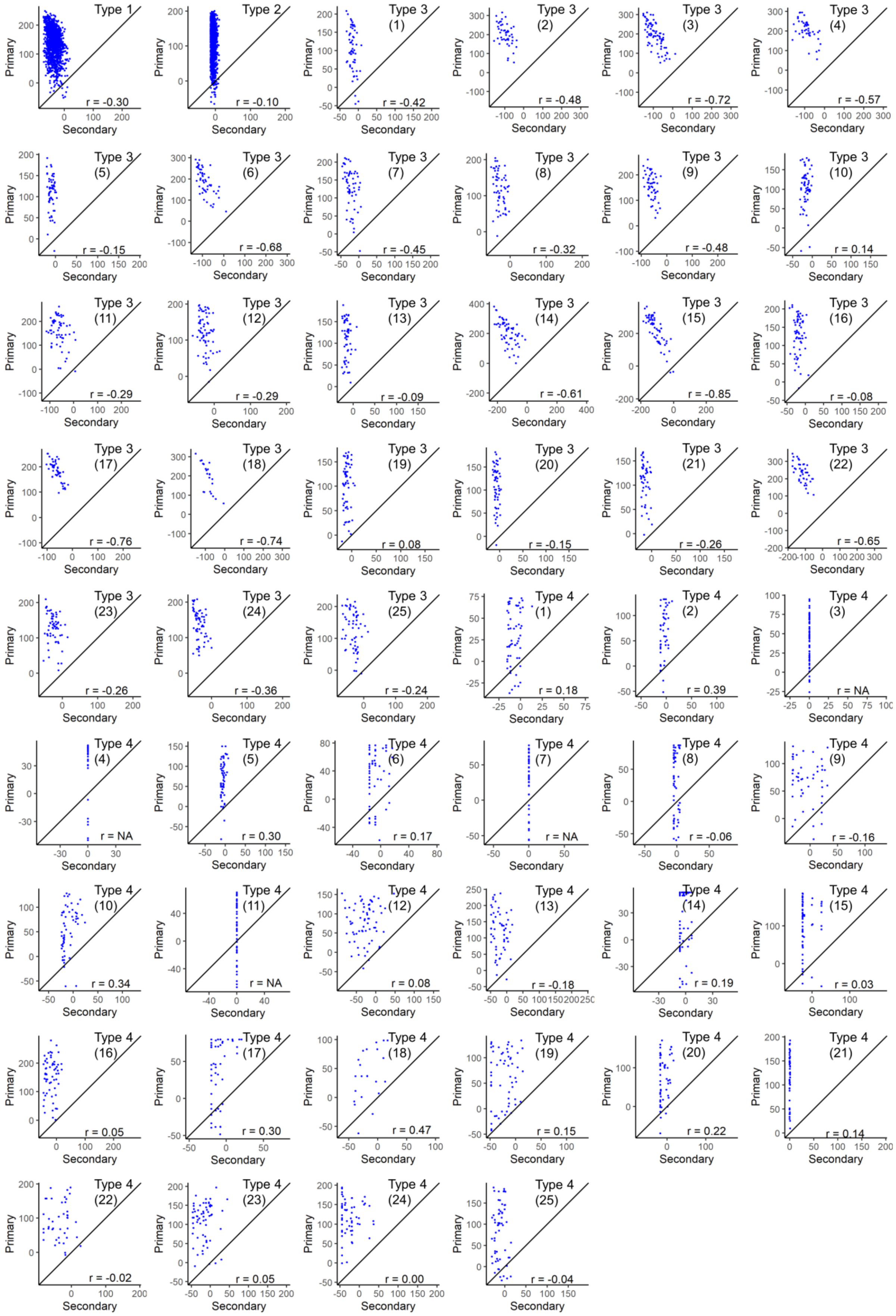


**Supplementary Figure 44. Predictions from favourable primary (P) vs secondary (S) markers of YLD (barley) for analysis type 1-4 using LASSO.**

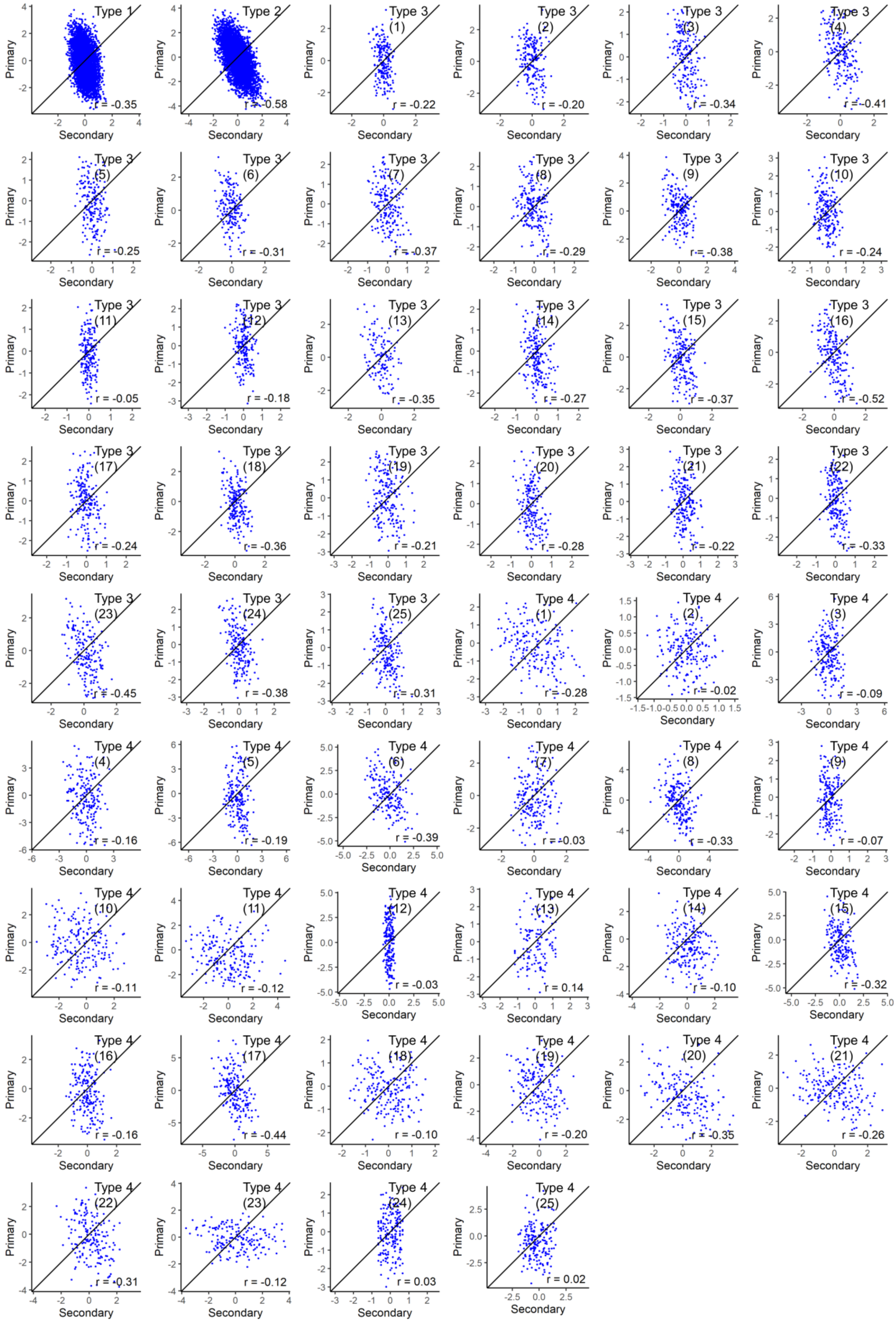


**Supplementary Figure 45. Predictions from favourable primary (P) vs secondary (S) markers of DTS (maize) for analysis type 1-4 using LASSO**.

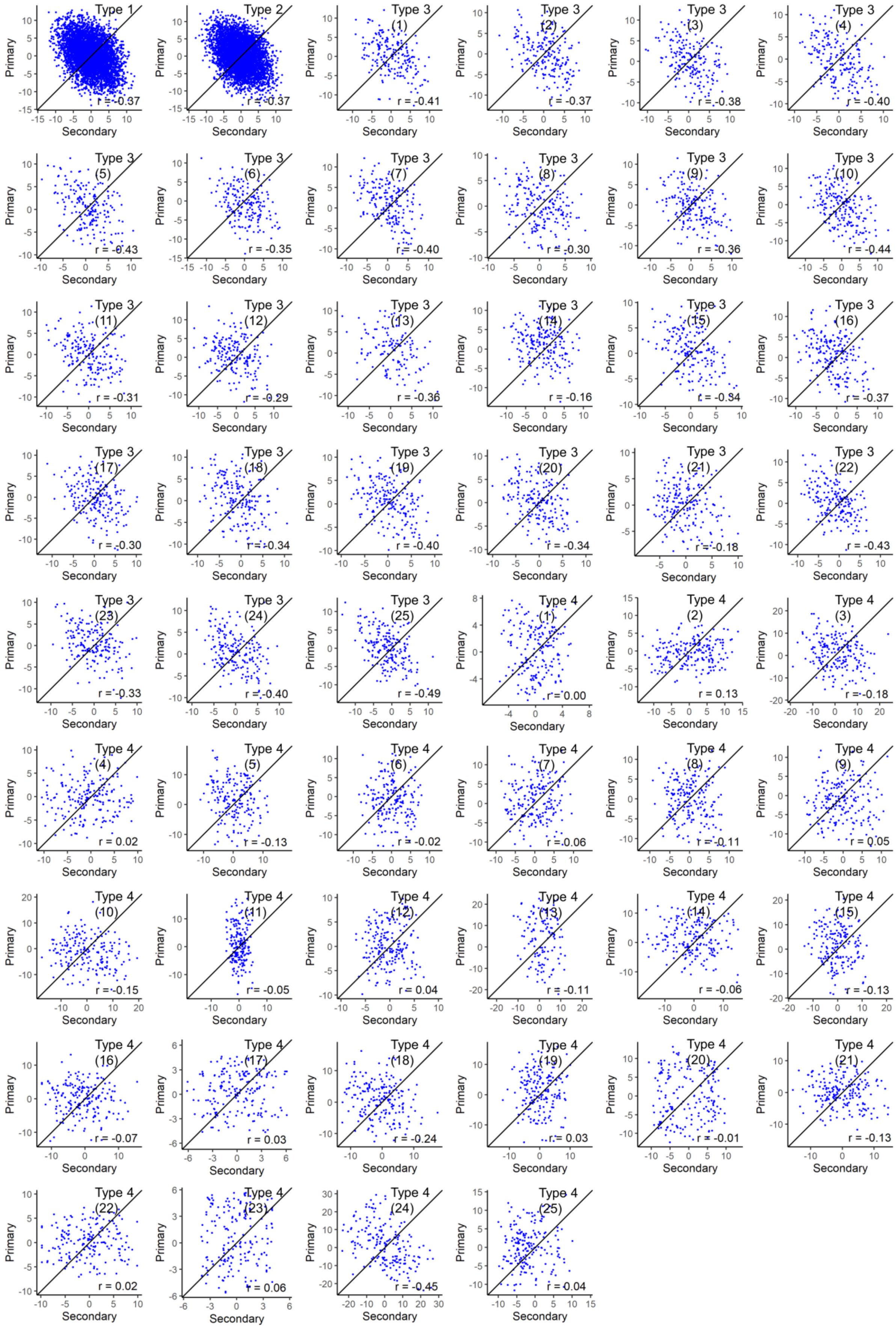


**Supplementary Figure 46. Predictions from favourable primary (P) vs secondary (S) markers of CL (maize) for analysis type 1-4 using LASSO.**

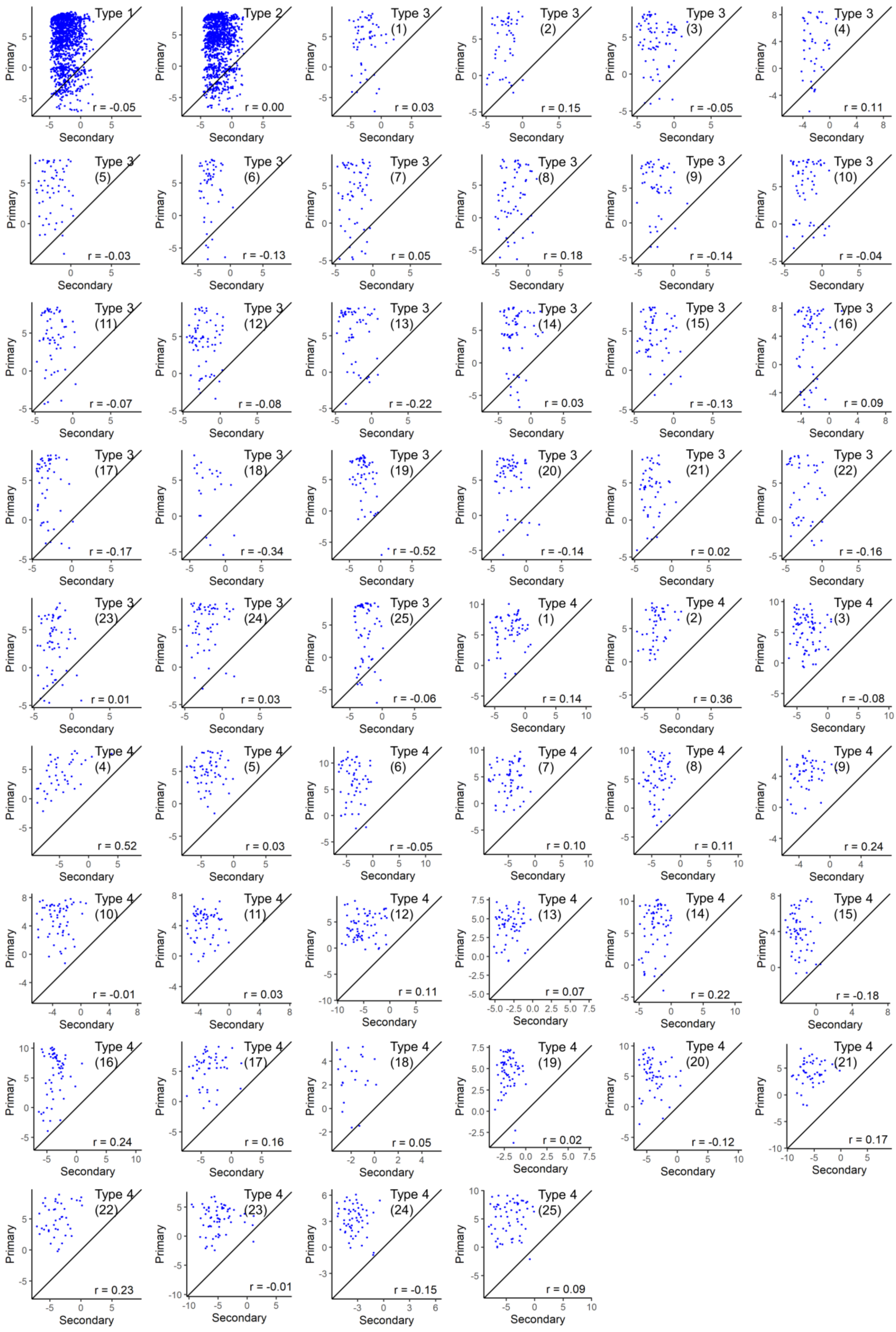


**Supplementary Figure 47. Predictions from favourable primary (P) vs secondary (S) markers of DTH (barley) for analysis type 1-4 using BayesCπ.**

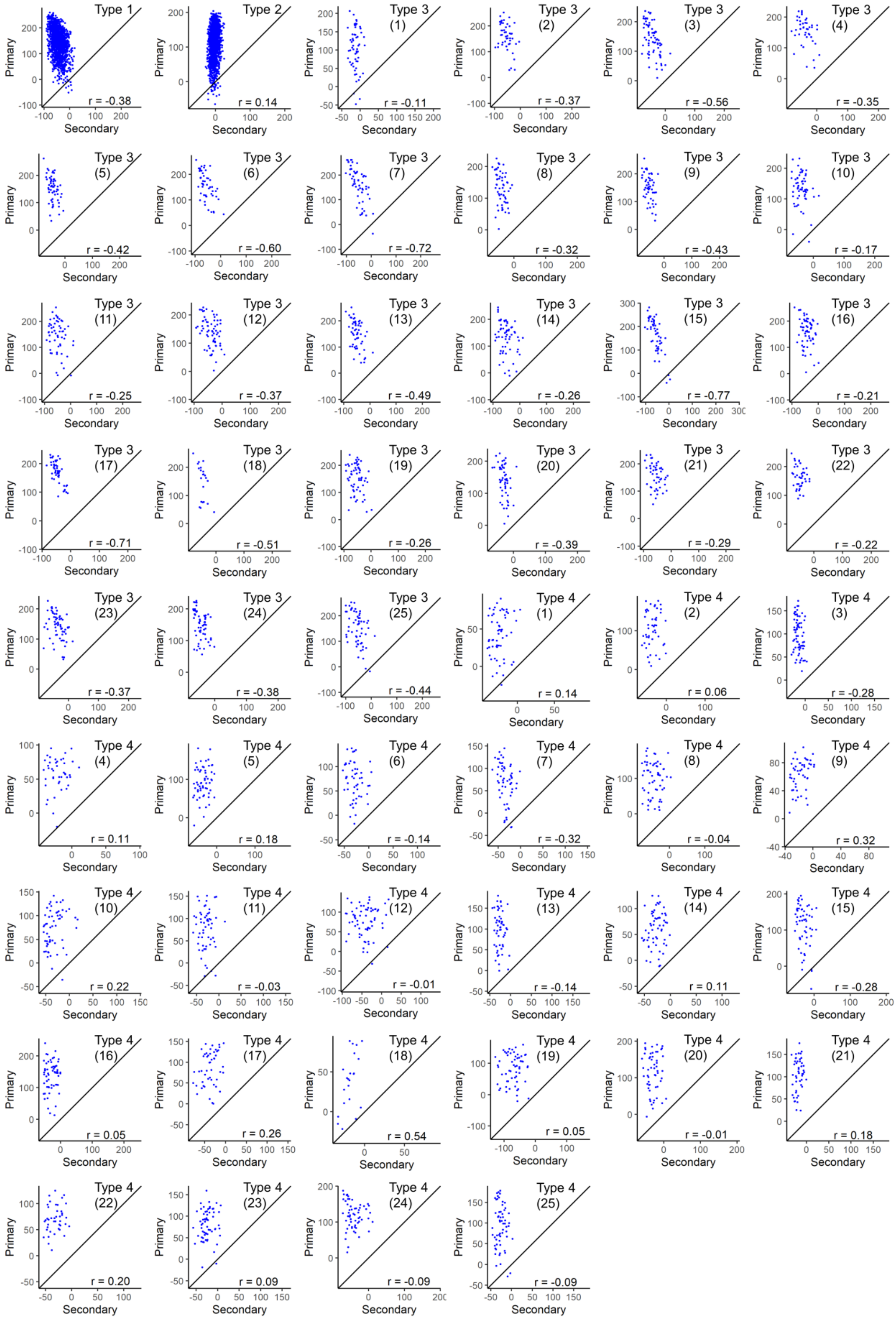


**Supplementary Figure 48. Predictions from favourable primary (P) vs secondary (S) markers of YLD (barley) for analysis type 1-4 using BayesCπ.**

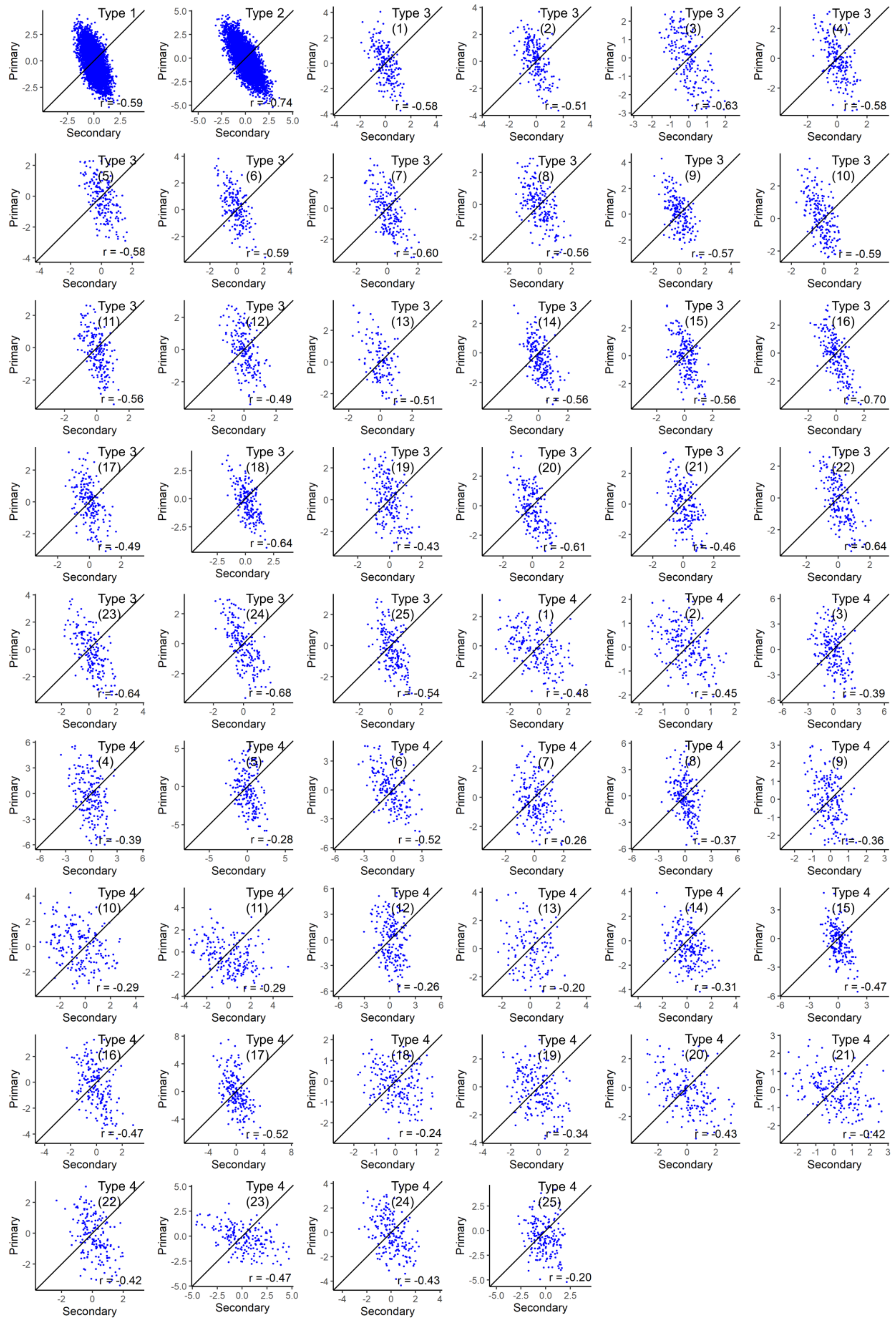


**Supplementary Figure 49. Predictions from favourable primary (P) vs secondary (S) markers of DTS (maize) for analysis type 1-4 using BayesCπ.**

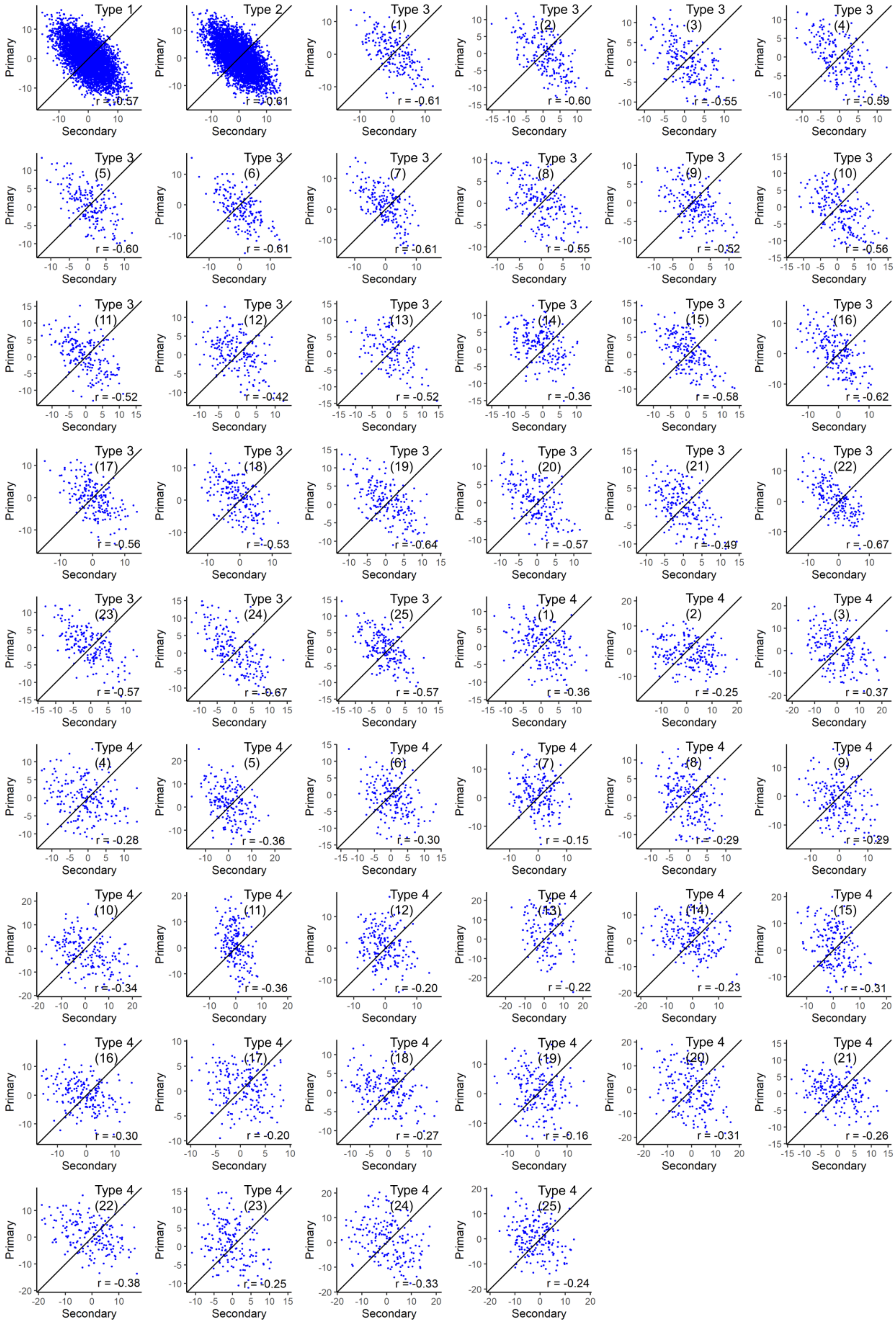


**Supplementary Figure 50. Predictions from favourable primary (P) vs secondary (S) markers of CL (maize) for analysis type 1-4 using BayesCπ.**

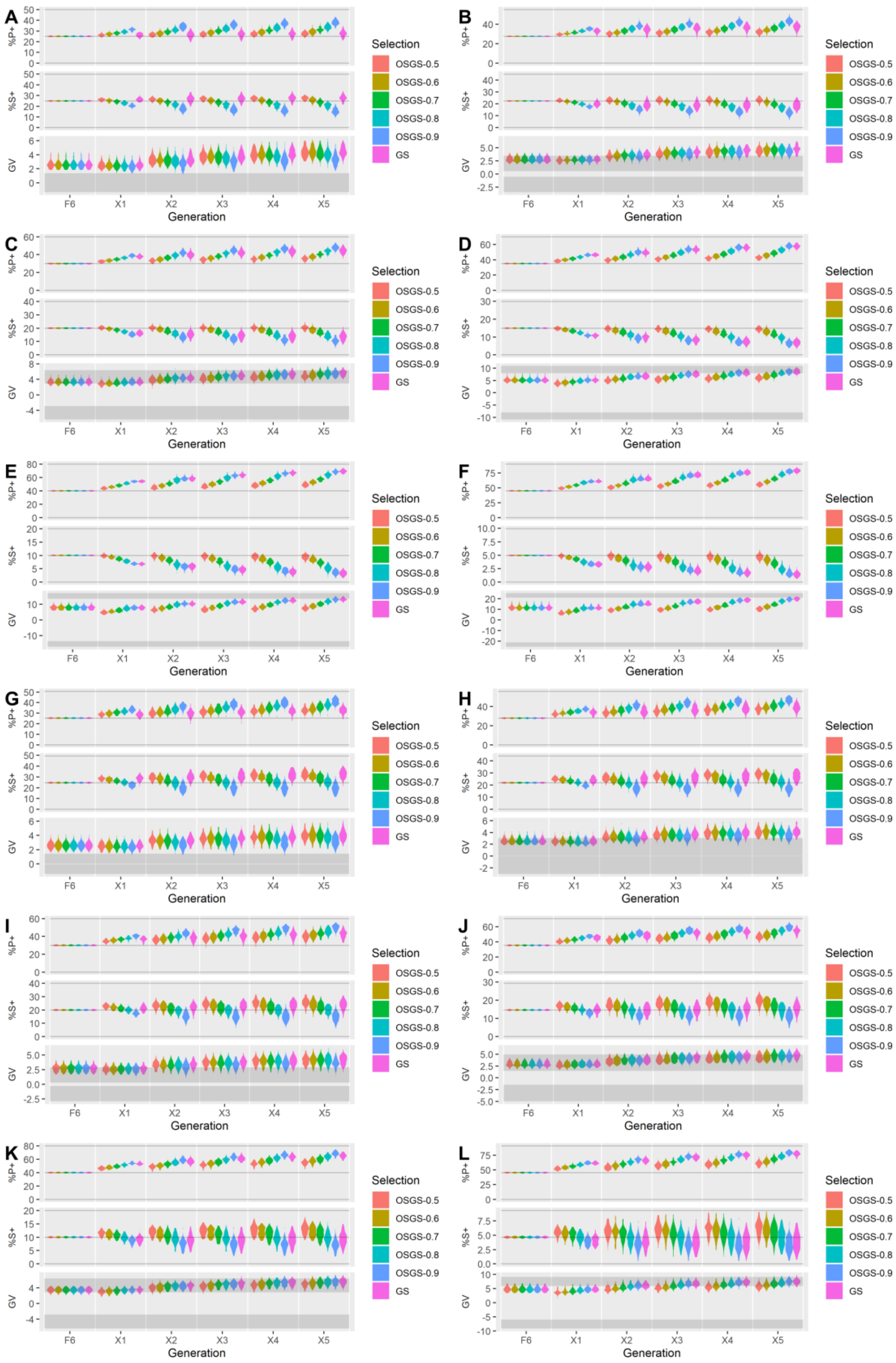


**Supplementary Figure 51. Simulation results comparing OSGS to GS in $F_2$-derived populations**. In each plot, %P+ is the percentage of favourable primary alleles, %S+ is percentage of favourable secondary alleles, and GV is the genetic values. Densities of %P+ and %S+ are plotted for the mean %P+ and %S+ over 100 simulations. Densities of GV are plotted for the best (maximum) GV over 100 simulations. [**A**] QTL density of 2cM/QTL & P:S ratio of 50:50. [**B**] 2 & 55:45. [**C**] 2 & 60:40. [**D**] 2 & 70:30. [**E**] 2 & 80:20. [**F**] 2 & 90:10. [**G**] 20 & 50:50. [H] 20 & 55:45. [**I**] 20 & 60:40. [**J**] 20 & 70:30. [**K**] 20 & 80:20. [**L**] 20 & 90:10.

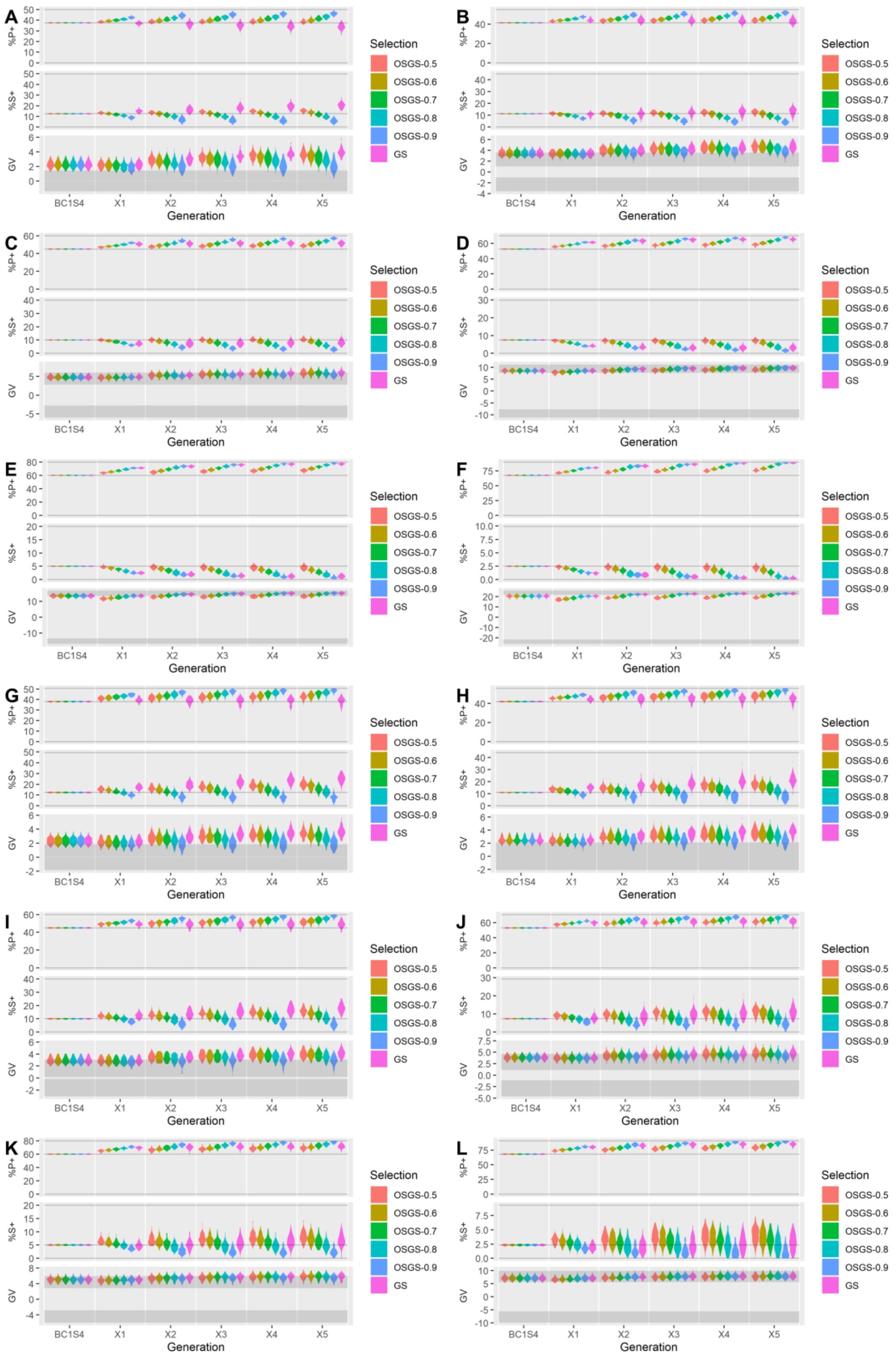


**Supplementary Figure 52. Simulation results comparing OSGS to GS in $BC_1$-derived populations**. In each plot, %P+ is the percentage of favourable primary alleles, %S+ is percentage of favourable secondary alleles, and GV is the genetic values. Densities of %P+ and %S+ are plotted for the mean %P+ and %S+ over 100 simulations. Densities of GV are plotted for the best (maximum) GV over 100 simulations. [**A**] QTL density of 2cM/QTL & P:S ratio of 50:50. [**B**] 2 & 55:45. [**C**] 2 & 60:40. [**D**] 2 & 70:30. [**E**] 2 & 80:20. [**F**] 2 & 90:10. [**G**] 20 & 50:50. [H] 20 & 55:45. [**I**] 20 & 60:40. [**J**] 20 & 70:30. [**K**] 20 & 80:20. [**L**] 20 & 90:10.

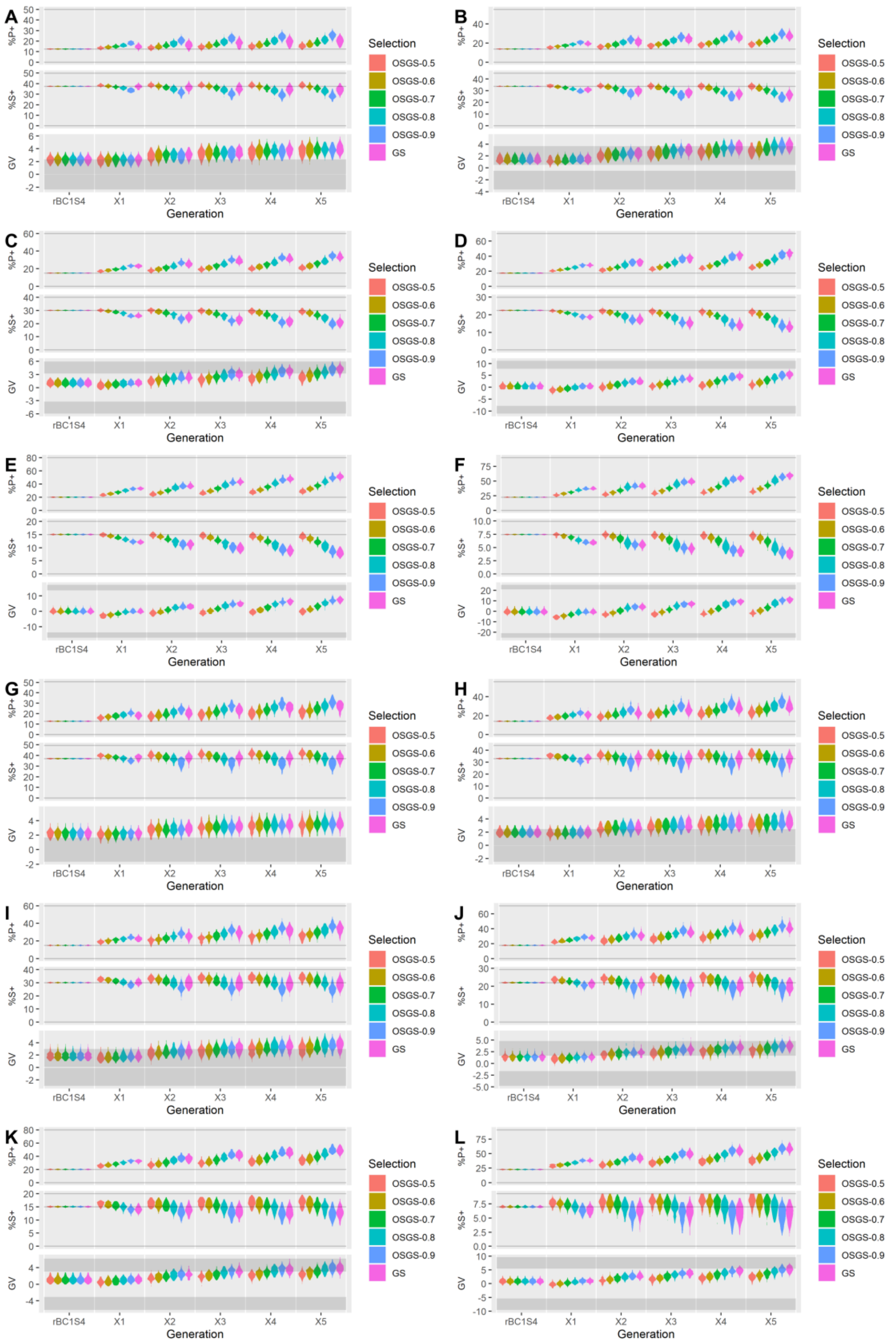


**Supplementary Figure 53. Simulation results comparing OSGS to GS in rBC$_1$-derived populations**. In each plot, %P+ is the percentage of favourable primary alleles, %S+ is percentage of favourable secondary alleles, and GV is the genetic values. Densities of %P+ and %S+ are plotted for the mean %P+ and %S+ over 100 simulations. Densities of GV are plotted for the best (maximum) GV over 100 simulations. [**A**] QTL density of 2cM/QTL & P:S ratio of 50:50. [**B**] 2 & 55:45. [**C**] 2 & 60:40. [**D**] 2 & 70:30. [**E**] 2 & 80:20. [**F**] 2 & 90:10. [**G**] 20 & 50:50. [H] 20 & 55:45. [**I**] 20 & 60:40. [**J**] 20 & 70:30. [**K**] 20 & 80:20. [**L**] 20 & 90:10.
